# RIPK2 Stabilizes c-Myc and is an Actionable Target for Inhibiting Prostate Cancer Metastasis

**DOI:** 10.1101/2020.05.14.096867

**Authors:** Yiwu Yan, Bo Zhou, Chen Qian, Alex Vasquez, Avradip Chatterjee, Xiaopu Yuan, Edwin Posadas, Natasha Kyprianou, Beatrice S. Knudsen, Ramachandran Murali, Arkadiusz Gertych, Sungyong You, Michael R. Freeman, Wei Yang

## Abstract

Despite advances in diagnosis and treatment, metastatic prostate cancer remains incurable and is associated with high mortality rates. Thus, novel actionable drug targets are urgently needed for therapeutic interventions in advanced prostate cancer. Here we report receptor-interacting protein kinase 2 (RIPK2) as an actionable drug target for suppressing prostate cancer metastasis. RIPK2 is frequently amplified in lethal prostate cancers and its overexpression is associated with disease progression and aggressiveness. Genetic and pharmacological inhibition of RIPK2 significantly suppressed prostate cancer progression *in vitro* and metastasis *in vivo.* Multi-level proteomic analysis revealed that RIPK2 strongly regulates c-Myc protein stability and activity, largely by activating the MKK7/JNK/c-Myc phosphorylation pathway—a novel, non-canonical RIPK2 signaling pathway. Targeting RIPK2 inhibits this phosphorylation pathway, and thus promotes the degradation of c-Myc—a potent oncoprotein for which no drugs have been approved for clinical use yet. These results support targeting RIPK2 for personalized therapy in prostate cancer patients towards improving survival.

## Main

Prostate cancer is the second most common cancer in men worldwide and causes about 360,000 deaths each year^1^. Mortality is predominantly caused by metastases that almost invariably become resistant to castration and other therapies^2^. Despite significant advances in prostate cancer treatment in the past decade^3^, the relative five-year survival rate of patients with metastatic disease remains about 30%^4^. Hence, there is an urgent need to identify novel actionable drug targets for combating metastatic progression to lethal disease.

To address this problem, we analyzed three large-scale clinical omics databases: the cBioPortal for Cancer Genomics^5^, the Prostate Cancer Transcriptome Atlas (PCTA)^6^, and the Pharos for human druggable genome^7^. We applied three stringent criteria to filter these databases: 1) genes are recurrently (>10%) amplified in lethal metastatic castration-resistant prostate cancer (mCRPC) (n=655)^8–10^, 2) mRNA levels have a high correlation (rho>0.9) with prostate cancer progression from normal to lethal disease (n=2,115), and 3) proteins can be readily targeted by T_clin_- or T_chem_-grade inhibitors, which are approved by the Food and Drug Administration (FDA) or have an activity cutoff of < 30 nM^7^. A total of 574, 1,655, and 2,208 human genes meet the three criteria, respectively, with an overlap of seven genes (Fig. 1a). Among these seven candidate druggable driver genes, receptor-interacting protein kinase 2 (RIPK2) has the highest mRNA level increase along with prostate cancer progression from benign to lethal disease (Fig. 1b). Currently, little is known about the importance and functions of RIPK2 in prostate cancer, although it has been well characterized in inflammation and innate immunity and was recently found to be involved in several other cancer types by activating the canonical RIPK2/NF-κB pathway^11,12^.

**Fig. 1.**
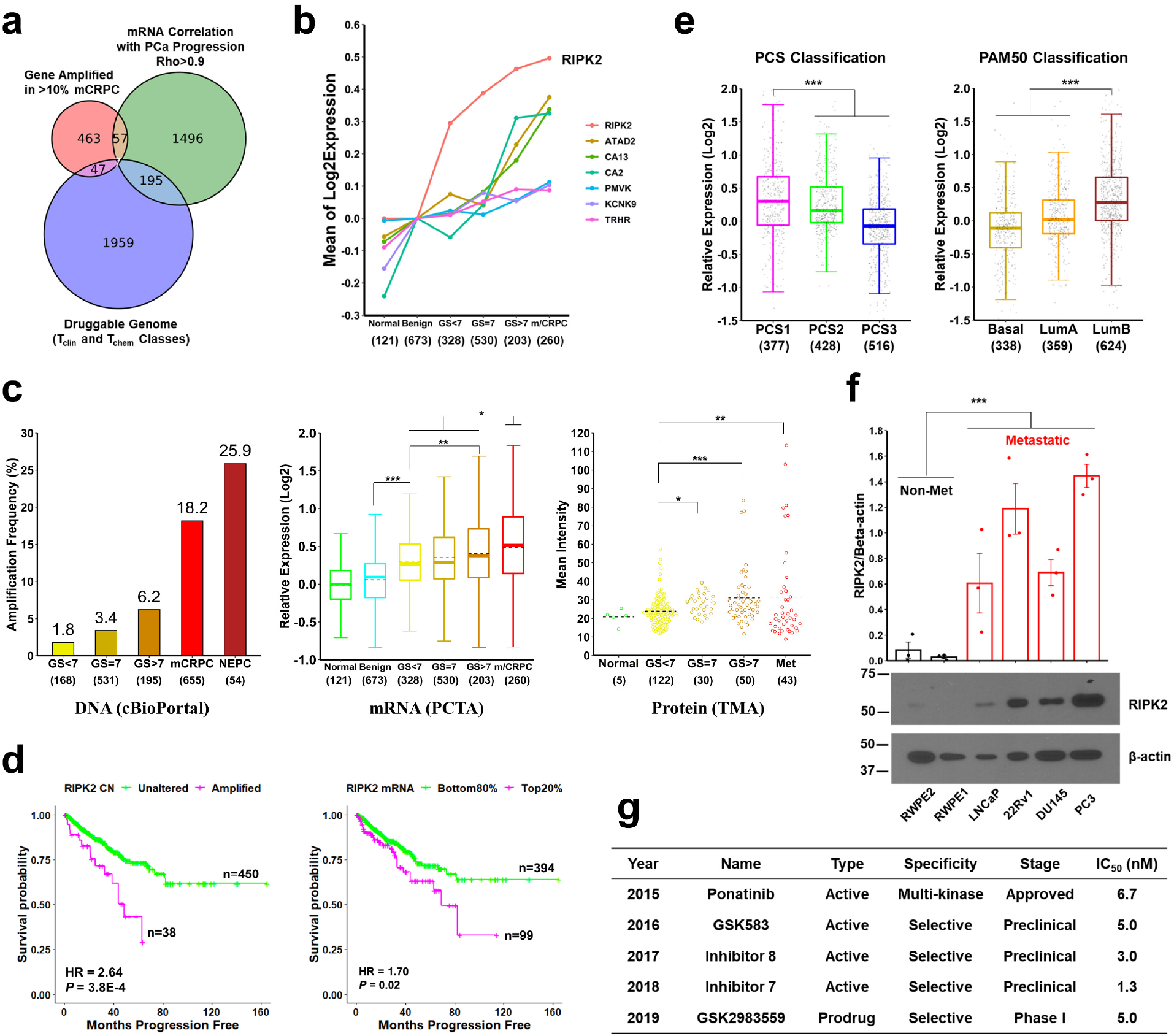
The overexpression of RIPK2—a druggable protein—is associated with prostate cancer progression and aggressiveness. **a,** Identification of candidate druggable drivers of prostate cancer progression to lethal disease. mCRPC, metastatic castration-resistant prostate cancer; PCa, prostate cancer. **b,** mRNA level changes of the seven overlapping genes along with prostate cancer progression. GS, Gleason score; m/CRPC, metastatic or castration-resistant prostate cancer; numbers in parentheses, sample sizes. **c,** Comparison of RIPK2 gene amplification (left), mRNA expression (middle), and protein expression (right) levels under different pathological conditions. Dotted lines indicate mean values. Box-and-whisker plots show the median values (solid line inside box), 25^th^ and 75^th^ percentiles (bottom and top of box, respectively), 25^th^ percentile – 1.5× interquartile range (bottom whisker), and 75^th^ percentile + 1.5× interquartile range (top whisker). *P* values were determined by unpaired two-tailed Student’s *t*-test. NEPC, neuroendocrine prostate cancer; Met, metastatic prostate cancer; TMA, tissue microarray. **d,** Kaplan-Meier progression-free survival analysis of prostate cancer patients (the TCGA PanCancer Atlas cohort) stratified based on whether RIPK2 is amplified (left) or highly expressed (right). Two-sided log-rank test. CN, copy number. **e,** Boxplot of RIPK2 mRNA levels in three PCS (left) and PAM50 (right) subtypes of prostate tumors in the PCTA cohort. Unpaired two-tailed Student’s *t*-test. **f,** Comparison of the protein expression levels of RIPK2 in six commonly used prostate cell line models. Unpaired two-tailed Student’s *t*-test. **g,** A list of RIPK2 inhibitors with the half-maximal inhibitory concentration (IC50) of < 10 nM. For the whole figure, * *P* < 0.05, ** *P* < 0.01, *** *P* < 0.001.

To determine whether RIPK2 is associated with prostate cancer progression in clinical tissue specimens at multiple molecular levels *(i.e.,* DNA, RNA, and protein), we analyzed publicly accessible prostate cancer genomics and transcriptomics datasets and performed immunohistochemical analysis of prostate cancer tissue microarrays. Indeed, RIPK2 expression increases with Gleason grade as well as along with the natural history of prostate cancer progression at all the three molecular levels (Fig. 1c, Extended Data Fig. 1). Moreover, the gene amplification and high mRNA abundance of RIPK2 are significantly associated with worse progression-free survival of prostate cancer patients in the Cancer Genome Atlas (TCGA) PanCancer Atlas^13^ cohort (Fig. 1d). In addition, RIPK2 mRNA levels are significantly higher in aggressive prostate cancer subtypes such as Prostate Cancer Subtype 1 (PCS1) and Luminal B (LumB), compared with the other two less aggressive PCS or PAM50 (prediction analysis of microarray 50) subtypes, in both PCTA and TCGA cohorts (Fig. 1e, Extended Data Fig. 2). Of note, PCS1 tumors progress more rapidly to metastatic disease than PCS2 and PCS3 tumors (hazard ratio = 4.8)^6^, while LumB tumors exhibit poorer clinical prognosis than LumA and Basal tumors^14^. Consistent with the clinical findings, RIPK2 protein abundance largely correlates with the aggressiveness of preclinical prostate cancer cell line models (Fig. 1f). Importantly, RIPK2 can be targeted by multiple potent small-molecule inhibitors such as GSK583 (a preclinical RIPK2-selective inhibitor) and Ponatinib (a multi-kinase inhibitor approved for the treatment of leukemia) (Fig. 1g, Extended Data Table 1)^15,16^. Collectively, these findings suggest that RIPK2 overexpression is associated with prostate cancer aggressiveness and that RIPK2 is potentially an actionable drug target for prostate cancer.

Using CRISPR/Cas9^17,18^, we stably knocked out RIPK2 from three androgen-independent prostate cancer cell lines: androgen receptor-negative PC3 and DU145 and androgen receptor-positive 22Rv1 (Fig. 2a). To determine whether RIPK2-knockout (RIPK2-KO) suppresses prostate cancer progression *in vitro,* we performed cell proliferation, transwell migration, matrigel invasion, clonogenic, and soft agar colony formation assays. The results showed that RIPK2-KO significantly reduced prostate cancer cell invasion, anchorage-dependent colony formation, and anchorage-independent soft agar colony formation (Fig. 2b-d). Nevertheless, RIPK2-KO only had a negligible effect on cell proliferation or migration in 2D culture *in vitro* (Extended Data Fig. 3). To determine whether RIPK2-KO suppresses prostate cancer metastasis *in vivo,* we conducted intracardiac injection of luciferase-labeled control and RIPK2-KO 22Rv1 cells in male SCID/Beige mice. Of note, 22Rv1 cells express both full-length and constitutively active truncated variants of androgen receptor, recapitulating aggressive mCRPC tumors^19^. Compared with the control group, mice harboring RIPK2-KO 22Rv1 cells showed significantly lower metastatic burden (84% reduction at week 4) (Fig. 2e-f, Extended Data Fig. 4). To determine whether RIPK2-KO affects tumor growth *in vivo*, we performed subcutaneous injection of control and RIPK2-KO 22Rv1 cells in male SCID/Beige mice. Compared with control tumors, RIPK2-KO 22Rv1 xenografts had a significantly reduced tumor growth rate and formed smaller tumors (Fig. 2g-h, Extended Data Fig. 5).

**Fig. 2.**
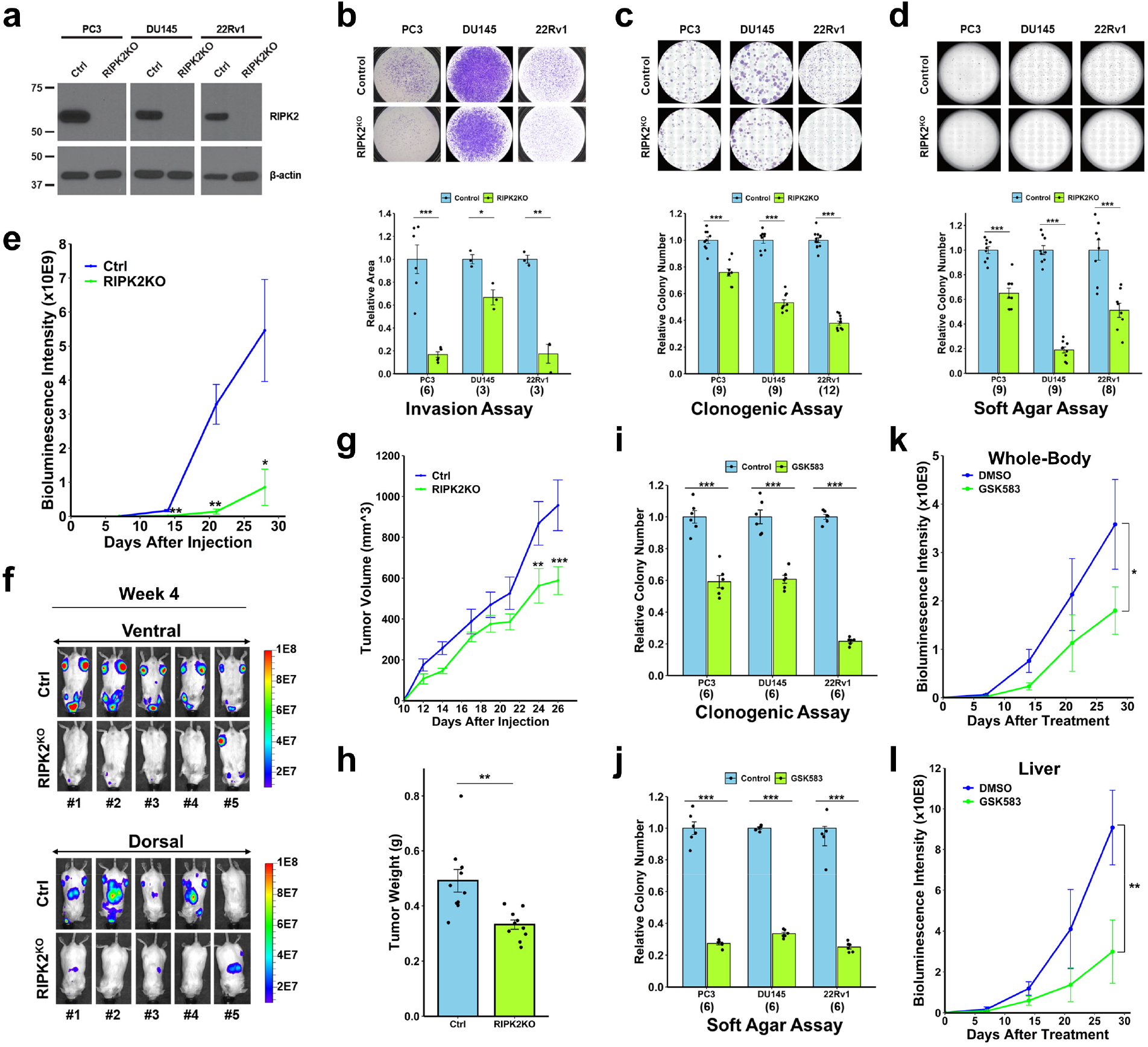
Genetic and pharmacological inhibition of RIPK2 suppresses prostate cancer progression and metastasis. **a,** Western blot analysis of RIPK2 protein levels in control and RIPK2-KO PC3, DU145, and 22Rv1 cells. Similar results were obtained in three independent experiments. **b-d,** Invasion (**b**), anchorage-dependent colony formation (**c**), and anchorage-independent soft agar colony formation (**d**) of control and RIPK2-KO PC3, DU145, and 22Rv1 cells. Numbers in parentheses represent the numbers of biological replicates. Unpaired two-tailed Student’s *t*-test. **e,** Effect of RIPK2-KO on 22Rv1 metastasis following intracardiac injection into male SCID/beige mice (n=5). Mann-Whitney *U* test. **f,** Bioluminescence images of the ventral (top) and dorsal (bottom) sides of mice bearing 22Rv1 control or RIPK2-KO tumors four weeks after intracardiac injection (n=5). **g,** Effect of RIPK2-KO on 22Rv1 xenograft tumor growth following subcutaneous injection into both flanks of male SCID/beige mice (tumor n=10). Mann-Whitney *U* test. **h,** The weights of 22Rv1 control and RIPK2-KO tumors (n=10) 26 days after subcutaneous injection. Unpaired two-tailed Student’s *t*-test. **i-j,** Anchorage-dependent (**i**) and -independent (**j**) colony formation of PC3, DU145, 22Rv1 cells (n=6) treated with GSK583 (10 μM) or vehicle. Unpaired two-tailed Student’s *t*-test. **k-l,** Effect of GSK583 treatment (10 mg per kg) on 22Rv1 metastasis in the whole body (**k**) or the mid-body (**l**) of male SCID/beige mice (vehicle, n=5; GSK583, n=7), starting 7 days after intracardiac injection. Two-way ANOVA. For the whole figure, data represent the mean ± standard error of the mean; * *P* < 0.05, ** *P* < 0.01, and *** *P* < 0.001.

Consistent with the *in vitro* results of genetic ablation, pharmacological inhibition of RIPK2 by GSK583 significantly suppressed anchorage-dependent and -independent colony formations of PC3, DU145, and 22Rv1 cells (Fig. 2i-j, Extended Data Fig. 6). To determine whether GSK583 can suppress prostate cancer metastasis in a setting that mimics the clinical scenario, we injected the aforementioned luciferase-tagged control 22Rv1 cells intracardially in male SCID/Beige mice. When metastases were observed by bioluminescence imaging at day 7, we randomized the mice into two groups (Extended Data Fig. 7), followed by daily treatment with GSK583 (10 mg per kg) or vehicle administered by oral gavage. In line with the KO result, GSK583 significantly suppressed the metastatic progression of 22Rv1 cells *in vivo* (50% reduction at week 4) (Fig. 2k and Extended Data Fig. 8). Of interest, GSK583 more significantly decreased the bioluminescence intensities in the ventral-side mid-body, which largely (if not exclusively) correspond to liver metastasis as supported by histopathological analysis (Fig. 2l, Extended Data Fig. 9). Notably, studies have shown that, among common visceral metastases, prostate cancer patients with liver metastases had the worst median overall survival^20^. It remains unknown why GSK583 has a higher efficacy on prostate cancer liver metastasis than overall metastasis. One possibility is that GSK583 has a better distribution in the liver, similar to nutlin-3a^21^. Importantly, the GSK583 treatment had no significant effect on mouse weight (Extended Data Fig. 10). Collectively, the results suggest that targeting RIPK2 is a viable strategy to inhibit the metastatic progression of prostate cancer.

As we did not detect significant changes in the activity of NF-κB (the canonical RIPK2 downstream effector) by RIPK2-KO in prostate cancer cells, we performed an unbiased label-free proteomic analysis to comprehensively identify RIPK2 downstream effectors (Extended Data Fig. 11). Here, we chose the control and RIPK2-KO PC3 cells for comparison, because PC3 has the highest RIPK2 protein expression level among all the cell lines that we screened (Fig. 1f). Database searching analysis identified 5,237 protein groups with a false discovery rate (FDR) of <1%. After quality assessment and statistical analysis (Extended Data Fig. 12-13, Table 2), we found that 243 and 409 protein groups are consistently downregulated or upregulated in all the three RIPK2-KO PC3 clones, compared with control PC3 cells, respectively *(q* < 0.005) (Fig. 3a, Extended Data Fig. 14-15, Table 3). ToppFun gene ontology enrichment analysis showed that the downregulated protein set is most significantly associated with ribosome biogenesis and mitochondria, whereas the upregulated protein set are most significantly associated with cytoskeleton organization and focal adhesion (Extended Data Fig. 16). Using the downregulated and upregulated protein sets, we respectively computed the Z scores of RIPK2-induced and RIPK2-repressed activities in each sample of the PCTA and TCGA cohorts. In line with the aforementioned RIPK2 mRNA findings (Fig. 1c, 1e, Extended Data Fig. 2), the RIPK2-induced activity scores are positively, whereas the RIPK2-repressed activity scores are negatively, associated with prostate cancer progression and aggressiveness in both cohorts (Extended Data Fig. 17).

**Fig. 3.**
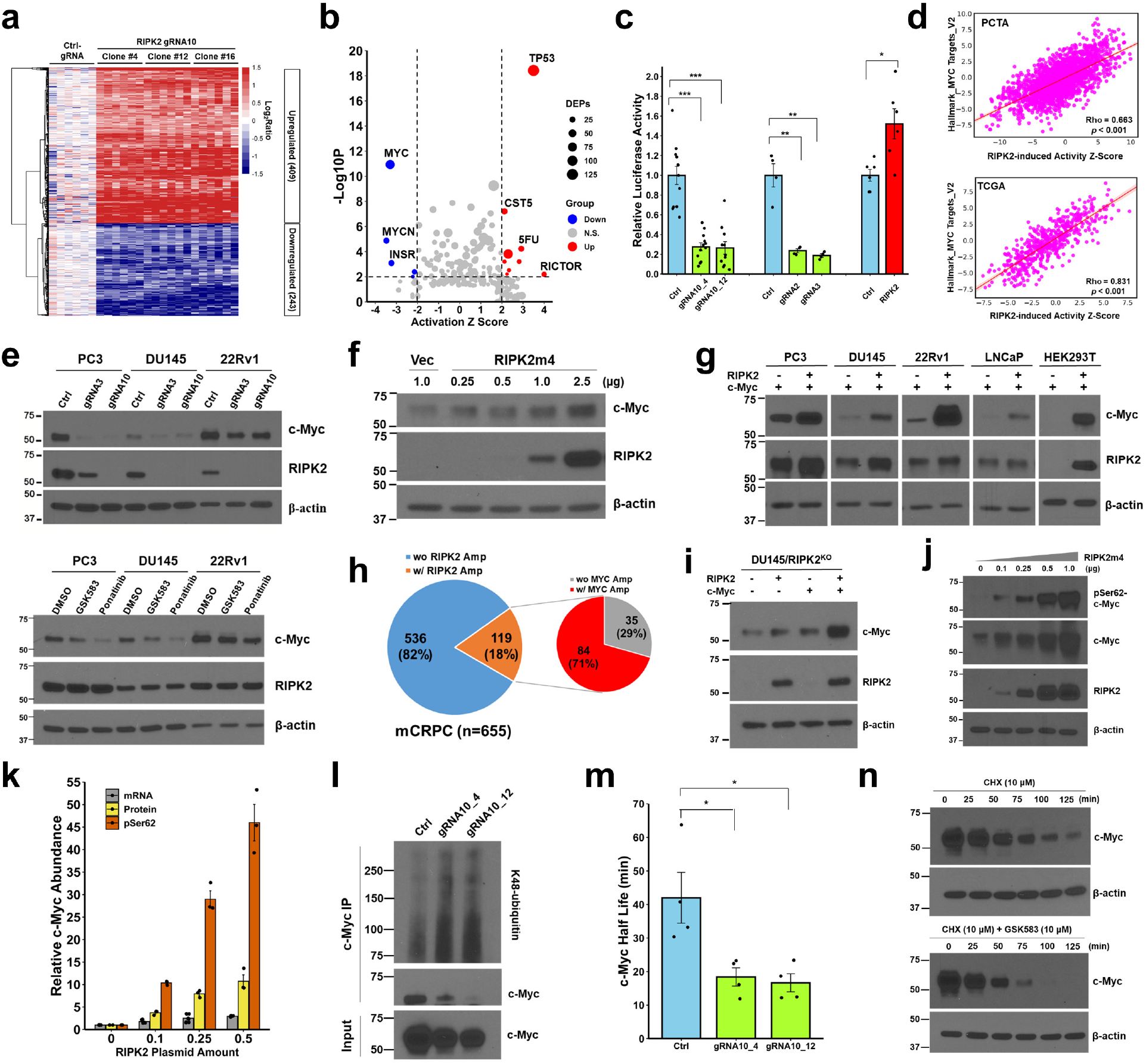
RIPK2 activates c-Myc by phosphorylating and stabilizing the c-Myc protein. **a,** Heatmap of the 652 protein groups that were consistently differentially expressed in three RIPK2-KO PC3 single-cell clones, compared with control PC3 cells. **b,** Scatter plot showing the -Log_10_*P* values against the activation Z scores (computed by Ingenuity Pathway Analysis) of putative regulators upstream of the 652 differentially expressed proteins (DEPs). Each dot represents a putative regulator. Dot sizes are proportional to the numbers of DEPs downstream of corresponding putative regulators. Red and blue represent putative regulators significantly activated and inactivated by RIPK2-KO, respectively. **c,** Bar graph of the relative c-Myc luciferase reporter activity. Data represent the mean ± standard error of the mean; unpaired twotailed Student’s *t*-test; * *P*<0.05, ** *P*<0.01, *** *P*<0.001. **d,** Spearman’s correlation between RIPK2-induced activity Z scores and Hallmark_MYC_Targets_V2 gene set activity Z scores in the PCTA (top) and TCGA Firehose Legacy (bottom) prostate cancer cohorts. **e,** Western blot analysis of the changes of c-Myc protein abundance in response to RIPK2-KO by CRISPR/Cas9 (top) as well as RIPK2 inhibitors GSK583 (10 μM) and Ponatinib (5 μM) (bottom). Similar results were obtained in three independent replicates. **f,** Dose-dependent increase of endogenous c-Myc protein abundance in RIPK2-KO PC3 cells in response to ectopic RIPK2 expression. **g,** Upregulation of exogenous c-Myc protein abundance in five different cell lines in response to ectopic RIPK2 expression, compared with the vector control. For each plasmid, 0.5 μg was used in transient transfection. **h,** Frequency of RIPK2 gene amplification (Amp) as well as RIPK2 and MYC co-amplification in mCRPC tissue specimens. **i,** Different levels of c-Myc protein increase in RIPK2-KO DU145 cells in response to the ectopic expression of RIPK2 and/or c-Myc (0.5 μg per plasmid). **j,** Dose-dependent increase of protein abundance of pS62-c-Myc and c-Myc in response to RIPK2 overexpression in RIPK2-KO HEK293T cells, which were co-transfected with 0.5 μg c-Myc plasmid. **k,** Quantification of the relative increase of c-Myc at the mRNA, protein, and pSer62 levels. Data represent the mean ± standard error of the mean. **l,** Increased K48-linked ubiquitination of c-Myc in response to RIPK2-KO in PC3 cells. **m,** Bar graph of the half-lives of c-Myc in control and RIPK2-KO PC3 cells. Data represent the mean ± standard error of the mean; unpaired two-tailed Student’s *t*-test; * *P*<0.05. **n,** Increased c-Myc degradation in response to GSK583 (10 μM) in PC3 cells.

To identify key mediators of RIPK2 signaling in prostate cancer, we conducted an upstream regulator analysis of the 652 differentially expressed protein groups using Ingenuity Pathway Analysis. The *in-silico* procedure showed that, among all putative upstream regulators, c-Myc was the most significantly suppressed by RIPK2-KO (Fig. 3b). In addition, as mentioned above, ribosome biogenesis—a key biological process regulated by c-Myc^22^—is the most significantly downregulated biological process by RIPK2-KO. Thus, c-Myc is potentially a central mediator of RIPK2 signaling in prostate cancer. Of note, c-Myc has long been recognized as a “most wanted” target for cancer therapy, as it contributes to the pathogenesis and progression of many human cancers including prostate cancer^23,24^. However, no direct c-Myc-targeting drugs have ever been approved by the FDA. Therefore, our study raises the possibility that targeting RIPK2 may be useful to treat c-Myc-driven prostate cancer. To the best of our knowledge, c-Myc has not yet been reported to be functionally associated with RIPK2.

To confirm that RIPK2 activates c-Myc signaling, we performed c-Myc luciferase reporter assays in RIPK2-manipulated PC3 cells. RIPK2-KO by three different gRNAs substantially decreased, whereas RIPK2 overexpression significantly increased, c-Myc activity in PC3 cells compared with their respective controls (Fig. 3c). Strikingly, in both PCTA and TCGA prostate cancer cohorts, RIPK2-induced activity scores are highly correlated with MYC activity scores, even though the overlap between the RIPK2 and MYC signature genes is modest (Fig. 3d; Extended Data Fig. 18). In comparison, RIPK2 mRNA levels have lower correlations with MYC activity scores, but still at levels comparable to those between c-Myc mRNA levels and MYC activity scores (Extended Data Fig. 18b). Interestingly, among all the human kinases, RIPK2 mRNA levels have the strongest correlation with MYC activity scores in the PCTA and TCGA cohorts on average (Extended Data Fig. 19). In addition, out of the 50 Hallmark gene sets in the Molecular Signatures Database (MSigDB)^25^, the MYC_Targets gene sets (V1 and V2) are among the most strongly associated with RIPK2 mRNA and activity levels in both cohorts (Extended Data Fig. 20). Taken together, RIPK2 is a strong activator of c-Myc in prostate cancer.

To investigate whether RIPK2 modulates c-Myc protein abundance, we performed functional inhibition: RIPK2-KO by two independent gRNAs and pharmacological inhibition of RIPK2 by GSK583 and Ponatinib. Our results showed that RIPK2 inhibition downregulated c-Myc protein levels in PC3, DU145, and 22Rv1 cells (Fig. 3e). In addition, ectopic expression of RIPK2m4, whose gRNA10-targeting spacer region was silently mutated to disable Cas9 recognition, increased the protein abundance of endogenous c-Myc in a dose-dependent fashion in RIPK2-KO PC3 cells (Fig. 3f), as well as that of exogenous c-Myc in five different cell lines (Fig. 3g).

Notably, the RIPK2 and MYC genes are located about 38 mega-bases apart on chromosome 8q, whose gains are among the most frequent cytogenetic alterations in prostate cancer^26^. Our analysis of three large-scale genomic profiling studies of mCRPC tissue specimens (n=655) showed that 18.2% of samples had RIPK2 amplification, of which 70.6% had RIPK2 and MYC co-amplification (Fig. 3h, Extended Data Fig. 21a). Interestingly, compared with the forced expression of RIPK2 or c-Myc alone, the co-transfection of both RIPK2 and c-Myc—a mimic of RIPK2 and MYC co-amplification—resulted in much higher c-Myc protein levels in RIPK2-KO DU145, 22Rv1, and HEK293T cells (Fig. 3i, Extended Data Fig. 21b). Thus, the co-amplification of RIPK2 and MYC, a relatively frequent event in mCRPC tumors, synergistically contributes to c-Myc protein abundance.

The regulation of c-Myc protein abundance by RIPK2 is mainly not at the transcriptional level (Extended Data Fig. 22). In human cells, the c-Myc protein can be stabilized through Ser62 phosphorylation, preventing its targeted degradation by the ubiquitin-proteasome system^27^. As expected, ectopic overexpression of RIPK2 increased the abundance of phospho-Ser62-c-Myc (pSer62-c-Myc) and c-Myc in a dose-dependent fashion (Fig. 3j). Moreover, the abundance of pSer62-c-Myc increased much faster than mRNA and protein abundance of RIPK2 (Fig. 3k), suggesting that RIPK2 regulates c-Myc largely by phosphorylating and stabilizing c-Myc. Consistently, RIPK2-KO significantly increased the K48-linked ubiquitination of c-Myc, a signal for proteasomal degradation^27^ (Fig. 3l). In addition, both RIPK2-KO and RIPK2 inhibition promoted proteasomal degradation and reduced the half-life of the c-Myc protein (Fig. 3m, 3n, Extended Data Fig. 23). Collectively, RIPK2 stabilizes the c-Myc protein by phosphorylating its Ser62 residue and preventing it from proteasomal degradation.

To determine whether RIPK2 binds to c-Myc, a prerequisite of direct RIPK2 phosphorylation of c-Myc, we performed proximity ligation assay and immunoprecipitation analysis. However, both failed to detect the protein-protein association between RIPK2 and c-Myc (Extended Data Fig. 24). Therefore, to identify the protein(s) linking RIPK2 to c-Myc-Ser62, we comprehensively profiled the interactome and the downstream phosphoproteome of RIPK2, followed by an integration analysis.

Prior to the interactome analysis, we discovered that the cytoplasmic location and the intact kinase domain are essential for RIPK2’s regulation of c-Myc (Fig. 4a, Extended Data Fig. 25). Thus, we postulated that certain protein(s) binding to the kinase domain, but not other regions, of cytoplasmic RIPK2 are critical for mediating RIPK2’s indirect phosphorylation of c-Myc-Ser62. To identify such protein(s), we performed a rigorously controlled interactome analysis by immunoprecipitation-mass spectrometry (IP-MS) (Extended Data Fig. 26-27). The analysis identified 1,189 proteins with an FDR of <1 %. After quality assessment and statistical analysis, we identified 219 protein candidates—including two known RIPK2-binding partners XIAP and RPL38—that associate with the kinase domain (and not other regions) of cytoplasmic RIPK2 (Fig. 4b, Extended Data Fig. 28-31). These candidates include six kinases (Extended Data Table 4), yet none of them was reported to be able to directly phosphorylate c-Myc-Ser62.

**Fig. 4.**
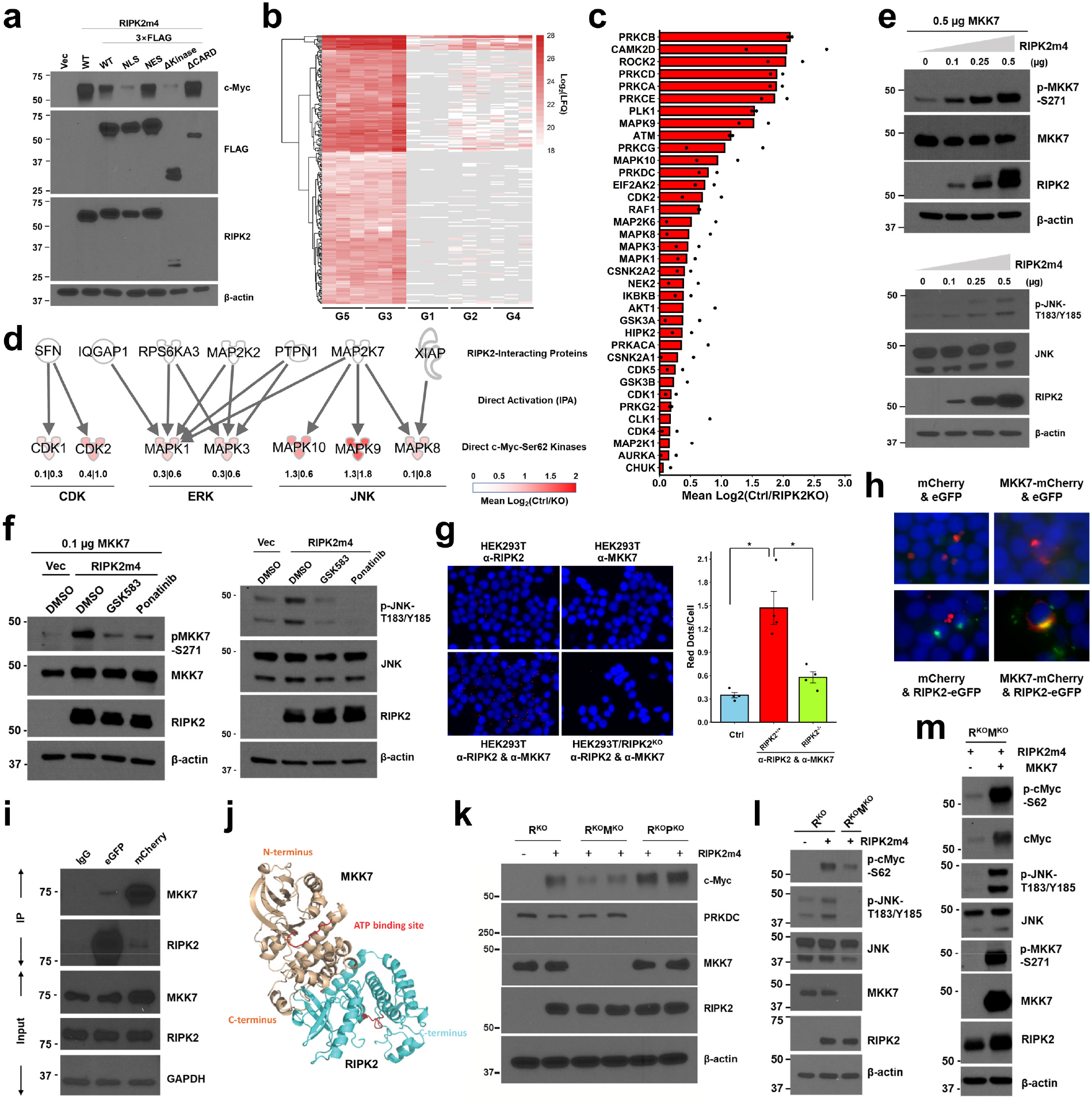
RIPK2 activates the MKK7/JNK/c-Myc signaling axis. **a,** Western blot analysis of c-Myc, FLAG-tag, and RIPK2 in RIPK2-KO HEK293T cells under the indicated conditions. β-actin was used as a loading control. Vec, vector control; WT, wild-type; NLS, nuclear localization signal; NES, nuclear export signal; Δkinase, deletion of the kinase domain; ΔCARD, deletion of the CARD domain. **b,** Heatmap of the 219 protein candidates specifically associated with the kinase domain of cytoplasmic RIPK2. G5 and G3 were experimental groups, whereas G1, G2, and G4 were control groups (see Extended Data Fig. 26 for more details). The heatmap cells in gray indicate zero label-free quantification (LFQ) intensities. **c,** Bar graph of the average activity change of inferred kinases downstream of RIPK2. A dot represents kinase activity in control PC3 cells relative to that in a RIPK2-KO PC3 single-cell clone (#4 or #12). **d,** Activation network connecting RIPK2-interacting proteins with direct c-Myc-Ser62 kinases downstream of RIPK2. The numbers indicate kinase activities in control PC3 cells relative to those in RIPK2-KO PC3 clone#4 (left) and clone#12 (right). **e,** Western blot analysis of p-MKK7-S271, total MKK7, p-JNK-T183/Y185, total JNK, and RIPK2 in RIPK2-KO HEK293T cells under the indicated conditions. β-actin was used as a loading control. In the upper panel, cells were co-transfected with MKK7 to enable the detection of p-MKK7-S271. **f,** Western blot analysis of target proteins in RIPK2-KO HEK293T cells under the indicated conditions. β-actin was used as a loading control. Vec and RIPK2m4, 0.1 μg; GSK583, 10 μM; Ponatinib, 5 μM. In the left panel, cells were co-transfected with MKK7 to enable the detection of p-MKK7-S271. **g,** Representative images (left) and a bar graph (right) of proximity ligation assay detecting the association of endogenous RIPK2 and MKK7 in HEK293T cells. **h,** Representative images of fluorescence colocalization assay of RIPK2 and MKK7 in HEK293T cells. **i**, Representative Western blots of RIPK2 and MKK7 in co-immunoprecipitation products and input samples. GAPDH was used as a loading control. **j**, Structural model showing that RIPK2 directly binds to MKK7 in a head-to-tail fashion. Three-dimensional structures of kinase domains are shown as ribbon representation. **k-m**, Representative Western blots of target proteins in HEK293T cells under the indicated conditions. Cells were co-transfected with 0.5 μg c-Myc plasmid to facilitate the detection of p-c-Myc-Ser62. β-actin was used as a loading control. R^KO^, RIPK2-knockout; R^KO^M^KO^, knockout of both RIPK2 and MKK7 genes; R^KO^P^KO^, knockout of both RIPK2 and PRKDC genes.

Next, to identify kinase(s) that link RIPK2 to c-Myc-Ser62, we performed a phosphoproteomic comparison of two RIPK2-KO PC3 clones with control PC3 cells (Extended Data Fig. 32). The analysis identified 6,749 phosphosites, which correspond to 2,716 phosphoproteins, with an FDR of <1% and a localization probability of >0.75. Following quality assessment and statistical analysis (Extended Data Fig. 33, Table 5), we quantified the relative activities of 47 kinases based on ≥ 5 substrate phosphosites per kinase, using the kinase-substrate enrichment analysis (KSEA) App^28^ (Extended Data Table 6). RIPK2 appeared to activate 36 kinases (Fig. 4c), of which nine were reported as direct c-Myc-Ser62 kinases but not as direct RIPK2 substrates.

The interactome and phosphoproteomics findings raised a possibility that RIPK2 binds to and activates a protein, which in turn activates a direct c-Myc-Ser62 kinase. Thus, we applied Ingenuity Pathway Analysis to connect the 219 RIPK2-interacting proteins with the nine direct c-Myc-Ser62 kinases, based on whether the former can directly activate the latter. Interestingly, seven RIPK2-interacting proteins can activate seven direct c-Myc-Ser62 kinases (Fig. 4d). Among the latter, c-Jun N-terminal kinases (JNKs) downstream of MAP2K7 (also known as MKK7), especially JNK2 (MAPK9), were most activated by RIPK2 (Fig. 4d). Of note, studies have shown that JNKs can directly phosphorylate c-Myc-Ser62 and that JNK2 deletion reduces c-Myc-Ser62 phosphorylation^29,30^. Thus, the MKK7/JNK pathway is potentially a central pathway mediating RIPK2’s phosphorylation and stabilization of c-Myc. In support of this hypothesis, RIPK2 activates MKK7 and JNK in a dose-dependent fashion, and this activation was abolished by GSK583 and Ponatinib (Fig. 4e, 4f, Extended Data Fig. 34-35).

To confirm the interactome finding that MKK7 associates with RIPK2 (Extended Data Fig. 36), we performed proximity ligation assay, fluorescence colocalization, and co-immunoprecipitation (Fig. 4g-i). Collectively, the results confirmed that MKK7 colocalizes and associates with RIPK2. In addition, structural modeling suggested that the N-terminal region of RIPK2 may directly bind to the C-terminal region of MKK7 (Fig. 4j). To validate that MKK7 is a key mediator of RIPK2’s regulation of c-Myc, we knocked out MKK7 from RIPK2-KO HEK293T cells with two different gRNAs by CRISPR/Cas9. Western blot analysis showed that MKK7-KO largely, although not completely, abolished RIPK2’s regulation of c-Myc (Fig. 4k, Extended Data Fig. 37). In comparison, the knockout of PRKDC, which encodes another RIPK2-interacting kinase (Extended Data Fig. 38), did not significantly affect RIPK2’s regulation of c-Myc (Fig. 4k, Extended Data Fig. 37). Moreover, MKK7-KO abrogated RIPK2-enhanced c-Myc and JNK phosphorylation, and the effects could be rescued by ectopic expression of MKK7 (Fig. 4l, 4m, Extended Data Fig. 39). Taken together, the MKK7/JNK axis is a major—albeit not the only—mediator of RIPK2’s phosphorylation and stabilization of c-Myc.

In summary, via integrated multi-omics analysis, we identified RIPK2 as an actionable drug target for the treatment of advanced prostate cancer, operating via a novel non-canonical RIPK2 signaling pathway. Our findings demonstrate that targeting RIPK2 inhibits the MKK7/JNK/c-Myc phosphorylation cascade, leading to the destabilization of the c-Myc protein and impairing prostate cancer progression to metastasis (Extended Data Fig. 40). Notably, RIPK2 is much more frequently amplified and overexpressed than any direct c-Myc-Ser62 kinases in prostate cancer (Extended Data Fig. 41). It is also frequently amplified or overexpressed in several other cancer types, and its overexpression is associated with poor clinical outcomes (Extended Data Fig. 42-43). Strikingly, RIPK2 activity scores are highly correlated with MYC activity scores across all cancer types in the TCGA PanCancer Atlas (Extended Data Fig. 44). Thus, clinical trials of targeting RIPK2—either alone or in combination with established therapies—are warrantied for personalized treatment of prostate cancer and potentially many other cancer types towards improving patient outcomes.

## METHODS

### Reagents and chemicals

SYBR Green PCR Master Kit (#4309155) was obtained from Applied Biosystems. iScript cDNA synthesis kit (#1708891), 2× or 4× Laemmli Sample Buffer (#1610737, #1610747), Mini-protean TGX gels (#4561096) were from Bio-Rad. MYC reporter kit (#60519) was from BPS Bioscience. GSK583 (#19739) and Ponatinib (#11494) were from Cayman Chemical. Collagen type I (#354249), Matrigel (#356231), and cell culture dishes and plates were from Corning. Fetal bovine serum (FBS) was from Denville. Transwell inserts (#353097) were from Falcon. BD Gram Crystal Violet (#212526) and Difco Agar Noble (#214220) were from Fisher Scientific. Iodoacetamide (#RPN6302) was from GE Healthcare. DMEM, RPMI-1640, fetal bovine serum (FBS), L-glutamine, penicillin, streptomycin, blasticidin, puromycin, and Opti-MEM (#51895) were from Gibco. Titansphere Titanium dioxide (5020-75000) was from GL Sciences. Microcon-30kDa (MRCF0R030) and Amicon Ultra-4 (#UFC803024) centrifugal filter units were from Millipore. Turbofectin (#TF81001) was from Origene. Dual-Luciferase Reporter Assay (#E1910) and Trypsin Gold (#V5280) were from Promega. Cycloheximide (#7698), MG132 (#474790), Cell Counting Kit-8 (#96992), Tris, SDS, sodium chloride, ammonium bicarbonate, and urea were from Sigma Aldrich. TRIzol Reagent (#15596026), 660nm Protein Assay Reagent (#22660), SuperSignal West Pico PLUS (#34577) and Femto (#34096) chemiluminescent substrates, dithiothreitol (DTT) (#20290), MS-grade trypsin (90059), LC-MS grade formic acid (#85178), 2-cm Acclaim PepMap 100 trap columns (#164564), and 50-cm EASY-Spray columns (ES803) were from Thermo Scientific. Autoradiography films were from Thomas Scientific.

### Constructs

pLenti-puro (#39481)^31^, LentiCas9-Blast (#52962)^32^, LentiGuide-Puro (#52963)^32^, single guide RNA (sgRNA) targeting human *RIPK2* (gRNA#10) (#76910)^33^, non-targeting guide RNA (#80255)^33^, pLJM-EGFP (#19319)^34^, mCherry2-N1 (#54517), pBMN (CMV-copGFP-Luc2-Puro) (#80389)^35^, pMD2.G (#12259), and psPAX2 (#12260) were obtained from Addgene. The canonical RIPK2 coding sequence (NM_003821) and MKK7 *(i.e.,* MAP2K7) coding sequence (NM_145185.4) were amplified by PCR from a RIPK2 vector (Origene, #RG202530) and a MKK7 vector (GenScript, Ohu28940), respectively, and cloned into the pLenti-puro vector.

### Generation of mutant and fusion cDNA

Primer sequences for molecular cloning were listed in Extended Data Table 7. Site-directed mutagenesis was conducted with the QuickChange II XL Site-Directed Mutagenesis Kit (Agilent, #200521). RIPK2m4 was generated by synonymous substitution of 648 A->T, 651 T->A, 654 C->A, and 657 A->C. The following fusion cDNAs were generated by PCR and cloned into the pLenti-puro vector: 3×FLAG-tag was fused to the C-terminus of target genes; NLS (CCAAAAAAGAAAAGAAAAGTT) and NES (CTACCGCCGCTGGAAAGACTGACTCTG) to the N-terminus of RIPK2m4; 6×His-tag to the C-terminus of MKK7. The RIPK2m4 sequence was cloned into the pLJM-eGFP vector to form RIPK2m4-eGFP. The MKK7 sequence was cloned into the mCherry2-N1 vector to form MKK7-mCherry. All plasmids were validated by sequencing (Laragen).

### RNA extraction and quantitative real-time polymerase chain reaction

Total RNA was extracted from cells using the TRIZOL (Invitrogen). Messenger RNA was converted to the first-strand cDNA using iScript cDNA synthesis kit (Bio-Rad), followed by RT-PCR reaction using SYBR Green PCR Master Kit in QuantStudio 5 Systems (Applied Biosystems). Gene expression was normalized to GAPDH using the comparative CT method.

### Cell lines

PC3, DU145, 22Rv1, LNCaP, RWPE-1, RWPE-2, and HEK293T cells were obtained from the American Type Culture Collection (ATCC). The cell lines were authenticated using the Promega PowerPlex 16 system DNA typing (Laragen). Unless otherwise specified, the cells were cultured in the media as follows. PC3, DU145, and HEK293T cells were cultured in DMEM, while 22Rv1 and LNCaP cells were cultured in RPMI-1640, supplemented with 10% fetal bovine serum (FBS), 2 mM L-glutamine, 100 units/mL penicillin and 100 μg/mL streptomycin. RWPE1 and RWPE2 cells were cultured in Keratinocyte Serum-Free Medium supplemented with 50 μg/mL bovine pituitary extract and 5 ng/mL human recombinant epidermal growth factor. All cells were cultured at 37°C in a 95% air, 5% CO_2_ humidified atmosphere. Mycoplasma contamination was routinely monitored using the MycoAlert PLUS Mycoplasma Detection Kit (Lonza, #LT07-118).

### Gene knockout by CRISPR/Cas9

sgRNAs targeting human *RIPK2* (#2 and #3), *MAP2K7* (#1 and #2), and *PRKDC* (#1 and #2) genes were designed using the sgRNA designer (Broad Institute). Their sequences and targeting domains are shown in Extended Data Table 8. All guide RNAs were cloned into LentiGuide-Puro (Addgene) according to the Zhang laboratory protocol, followed by sequencing validation (Laragen). Transfer (LentiGuide-Puro or LentiCas9-Blast), packing (psPAX2), and envelope (pMD2.G) plasmids were co-transfected into HEK293T cells to produce lentivirus containing sgRNAs and Cas9. After 72 h, cell supernatants were filtered with 0.45 μm filters and stored at −80°C for future use. To generate stable cell lines with gene knockout, cells were cultured in 6-well plates for 24 h to reach ~50% confluency, followed by infection with lentivirus of Cas9 and sgRNA supplemented with 10 μg/mL polybrene. After 72 h, medium was replaced by fresh medium supplemented with 10% FBS, 2mM L-glutamine, 100 units/mL penicillin, 100μg/mL streptomycin, 10μg/mL blasticidin and 1μg/mL puromycin. The selective medium was replaced every other day.

### Isolation of stable single cell clone

Single-cell clones were isolated by seeding individual cells in 96-well plates (5 cells/mL, 100 μL per well). About 10 days later, monoclonal cells were marked, monitored for growth, and sub-cultured.

### Transient overexpression

When cells grew to a confluency of ~50%, a specified amount of plasmid was transfected into cells using Turbofectin 8.0 (Origene) by following the manufacturer’s instruction. Briefly, plasmids and Turbofectin 8.0 were mixed at a ratio of 1:3 (w/v) to Opti-MEM, followed by incubation at room temperature for 15 min. The mixture was then added dropwise to cultured cells and incubated for 48 h at 37°C.

### Cell proliferation assay

The growth curve assay was performed using the Cell Counting Kit-8 (CCK-8) (Sigma-Aldrich, #96992) according to the manufacturer’s instruction. Briefly, 100 μL of cells (2,500 cells/mL for PC3 and DU145 and 10,000 cells/mL for 22Rv1) were plated in 96-well plates at day 0. A mixture of 10 μL CCK-8 reagent and 10 μL Dulbecco’s phosphate-buffered saline (DPBS) was added into each well at indicated time points. Plates were incubated in a humidified incubator at 37°C for 2 h, followed by measuring the absorbance at 450 nm using a microplate reader (800 TS, BioTek).

### Migration assay

Migration assay was performed essentially as described^36^. The outside membrane of Transwell inserts (8 μm pore size) was coated with 50 μL of 15 μg/mL Collagen Type I. A total of 5 × 10^4^ control or RIPK2-KO PC3 cells in 200 μL serum-free medium were seeded into the upper chamber, followed by adding 600 μL 10% FBS-containing medium to the lower chamber. After incubation at 37°C for 6 h, cells were fixed by 4% paraformaldehyde and stained by 0.12 mg/mL crystal violet solution for 15 min. Cells on the inside membrane of Boyden chambers were removed by cotton swabs. Migrated cells on the outside membrane were rinsed with deionized water for three times and imaged under an All-in-one Keyence microscope. The total area (μm^2^) of migrated cells were automatically calculated using the Keyence image analyzer.

### Invasion assay

Invasion assay was conducted essentially as described^36^. The inside membrane of Transwell inserts (8 μm pore size) was coated with 50 μL Matrigel matrix (1:50 diluted). For each sample, 200 μL (1×10^6^ cells/mL) of control or RIPK2-KO PC3, DU145, or 22Rv1 cells suspended in serum-free medium were seeded in the upper chamber. 600 μL of 10% FBS-containing medium was added to the lower chamber. After incubation at 37°C for 24 h, cells were fixed, stained, cleaned, imaged, and quantified as described in the migration assay.

### Clonogenic assay (anchorage-dependent colony formation assay)

Two milliliters of PC3 (250/mL), DU145 (250/mL), or 22Rv1 (1,000/mL) cells were seeded in 6-well plates and cultured in 10% FBS-containing media, which were replaced every 2-3 days. After 10-14 days, cells were fixed in 4% paraformaldehyde for 15 min and rinsed with DPBS for three times. Cells were stained with 120 μg/mL crystal violet solution. Cell colonies were quantified under an All-in-one microscope (Keyence) as described in the migration assay.

### Soft agar assay (anchorage-independent colony formation assay)

Soft agar assay was performed essentially as described^37^. Briefly, 1.5 mL of 0.5% Nobel agar solution was plated in each well of 6-well plates as the bottom layer. 1.5 mL of 6.67 × 10^3^/mL (for PC3 and DU145) or 1.33×10^4^/mL (for 22Rv1) cells in 0.3% noble agar solution were plated in 6-well plates as the top layer. Cells were cultured in DMEM supplemented with 10% FBS, 2mM L-glutamine, 100 units/mL penicillin and 100μg/mL streptomycin. The medium was replaced every 2-3 days. After 2-3 weeks, cell colonies were stained with 0.1% crystal violet solution for 20 minutes and washed with deionized water for five times. Cell colonies were quantified under an All-in-one microscope (Keyence) as described in the migration assay.

### Animal studies

All experimental protocols and procedures were approved by the Institutional Animal Care and Use Committee at Cedars-Sinai Medical Center. All relevant ethical regulations, standards, and norms were rigorously adhered to. For *In vivo* tumor growth assay, control or RIPK2-KO 22Rv1 cells were adjusted to 1.5 × 10^7^ cells/mL in DPBS, followed by mixing with Matrigel at a 1:1 ratio (v/v). For each male SCID/Beige mouse (7-week old; Charles River), 100 μL mixture was subcutaneously injected into both flanks. Tumor length and width were measured with a caliper for three times each week, and tumor volumes were calculated using the formula of (length × width^2^)/2. At the endpoint, mice were euthanized and tumor xenografts were collected and weighted with a scale. For *In vivo* metastasis assay, pBMN (CMV-copGFP-Luc2-Puro) (Addgene) was stably transfected into control and RIPK2-KO 22Rv1 cells. Two weeks later, luciferase-expressing cells were sorted with a FACSAria III (BD Biosciences) using the same gate for GFP. Control and RIPK2-KO 22Rv1 cells expressing similar levels of luciferase were selected and expanded. To each male SCID/Beige mouse (7-week old; Charles River), 100 μL of 1 × 10^7^ cells/mL in DPBS was intracardially injected into the left cardiac ventricle. Each week, 150 μL of 30 mg/mL D-luciferin was intraperitoneally injected into each mouse, followed by measuring tumor metastasis with an IVIS Spectrum In Vivo Imaging System (PerkinElmer). Luciferase activity was quantified by the Living Image software (Caliper Life Sciences, v4.3.1). For GSK583 treatment, to each male SCID/Beige mouse (8-week old; Charles River), 100 μL of 2.5 × 10^6^ cells/mL in DPBS was intracardially injected. Seven days later, mice were randomized into two groups—one group was administered by oral gavage with GSK583 (10 mg per kg, n=7), and the other with vehicle control (3% DMSO, n=5) every day. Metastases were visualized by bioluminescence imaging every week as described above.

### Myc luciferase assay

The c-Myc activity was assessed using the Myc Reporter kit (BPS Biosciences) and the Dual-Luciferase Reporter System (Promega) according to the manufacturers’ instructions. Briefly, 100 μL (1.5 × 10^5^ cells/mL) control and RIPK2-KO PC3 cells were seeded into 96-well plates. After overnight incubation, when cells reached ~50% confluency, 1 μL of Reporter A (60 ng/μL) in the Myc Reporter kit was transfected into cells using Turbofectin 8.0. After 48 h, cells were lysed in 25 μL Passive Lysis Buffer (provided in the Dual-Luciferase Reporter kit). 20 μL of cell lysate was transferred to 96-well plates and placed in a 96-well microplate luminometer (GloMax-Multi, Promega). 100 μL Luciferase Assay Reagent II and 100 μL Stop & Glo Reagent (both provided in the Dual-Luciferase Reporter kit) were sequentially injected, and firefly and Renilla luciferase activities were automatically measured. c-Myc activities were determined by the ratios of firefly to Renilla luciferase activities.

### Label-free proteomics

Label-free proteomics was performed essentially as described^38^. Briefly, after cell lysis, 60 μg protein per sample was digested into tryptic peptides using the filter-aid sample preparation (FASP) method^39^. About 1 μg peptides in 10 μL solution were separated on a 50-cm EASY-Spray column using a 200-min LC gradient at the flow rate of 150 nL/min. Separated peptides were analyzed with an LTQ Orbitrap Elite (Thermo Scientific) in a data-dependent manner. The acquired MS data (24 RAW files) were searched against the Uniprot_Human database (released on 01/22/2016, containing 20,985 sequences) using the Andromeda algorithm^40^ in the MaxQuant (v1.5.5.1) environment^41^. A stringent 1% FDR was used to filter the identifications of peptide-spectrum matches, peptide, and protein groups. Note that in bottom-up proteomics analysis, different proteins identified by the same set of shared peptides cannot be distinguished, so they are collapsed into a “protein group” to minimize redundant identifications. The mass spectrometry proteomics data have been deposited to the ProteomeXchange Consortium (http://proteomecentral.proteomexchange.org) via the PRIDE^42^ partner repository with the database identifier PXD18890, where more detailed experimental information was included. Perseus (v1.6.6.0)^43^ was applied to perform quality assessment and statistical analysis. Proteins identified from the reverse decoy and contaminating protein sequence databases as well as those with site-only identifications were removed. For statistical comparison, all LFQ intensity values were log2-transformed, and only proteins with ≥3 valid values in each group were used. Unpaired two-tailed Welch’s *t*-test followed by Benjaminin-Hochberg adjustment^44^ was used to calculate *p* and *q* values, respectively. To compute the combined *p* and *q* values, Stouffer’s method followed by Benjaminin-Hochberg adjustment was applied.

### Interactome analysis by immunoprecipitation-mass spectrometry (IP-MS)

Cells were cultured in 150-mm dishes until reaching the confluency of ~50%, and then transfected with 15 μg of indicated plasmids for 48 h. After cell lysis, 500 μL of 2 mg/mL protein solution was pre-cleared by incubating with 60 μL of 50% immobilized protein A/G Plus agarose bead slurry (Thermo Scientific, #20423) for 2 h at 4°C. The pre-cleared protein solution was incubated with 3 μg anti-FLAG antibody (Sigma Aldrich, #F1804) or IgG (Millipore, #12-371) overnight at 4°C. The next day, 60 μL of 50% immobilized protein A/G gel slurry was added to each sample, followed by incubation on a vertical shaker for 2 h at 4°C. After washing for five times, bound proteins were eluted with 50 μL of 2× Laemmli sample buffer containing 5% β-mercaptoethanol, by heating at 95°C for 5 min. Eluted proteins were analyzed by gel-enhanced liquid chromatography-tandem mass spectrometry (GeLC-MS/MS) essentially as described^38,45^. Briefly, eluted proteins were resolved by short-range SDS-PAGE, reduced, alkylated, and digested with trypsin (1:50, w/w) for 16 h in the gel. Tryptic peptides were analyzed using an EASY-nLC 1200 connected to an Orbitrap Fusion Lumos (Thermo Scientific) operated in a data-dependent manner. The acquired MS data (20 RAW files) were searched against the Uniprot_Human database (released on 03/30/2018, containing 93,316 sequences) using MaxQuant (v1.5.5.1). The data have been deposited to the PRIDE with the database identifier PXD18870, where more detailed experimental information was provided. Perseus (v1.6.6.0) was applied to perform quality assessment and statistical analysis as described in the label-free proteomics analysis. For each comparison, only proteins with ≥3 valid values in at least one group were used. Missing data were imputed from normal distribution by Perseus, using the default values (width 0.3; down shift 1.8). The *p* and *q* values were computed as described above.

### Phosphoproteomics

From regularly cultured control and RIPK2-KO PC3 cells (described in the label-free proteomics section), 1 mg protein was reduced, alkylated, and digested with trypsin in Amicon Ultra-4 centrifugal filter units (Millipore, #UFC803024) using the FASP method. To the resulting peptide solution, 1.5 mL acetonitrile, 7.5 mL of Incubation Buffer (60% acetonitrile, 3% trifluoroacetic acid), and 2 mg equilibrated TiO2 beads were sequentially added and mixed, followed by incubation for 60 min. TiO2 beads were washed with 1 mL of 60% acetonitrile, 3% trifluoroacetic acid, 50 mM citric acid three times (20 min per time) and 1 mL of 80% acetonitrile, 0.1% trifluoroacetic acid for 1 min once. Phosphopeptides were eluted with 100 μL of 50% acetonitrile, 14% ammonium hydroxide and then 100 μL of 80% acetonitrile, 5.6% ammonium hydroxide (5 min incubation per elution). Peptide solution resulting from the two elution steps was combined and dried down in a SpeedVac. Enriched phosphopeptides were analyzed using an EASY-nLC 1200 connected to an Orbitrap Fusion Lumos operated in a data-dependent manner. The acquired MS data (9 RAW files) were searched against the Uniprot_Human database (released on 01/22/2016) using MaxQuant (v1.5.5.1). The data have been deposited to the PRIDE with the database identifier PXD18871, where more detailed experimental information was included. Perseus (v1.6.6.0) was applied to perform quality assessment and statistical analysis, as described in the interactome analysis. To compute relative phosphorylation level changes, relative phosphosite intensities were normalized against the relative abundance of corresponding proteins, which were quantified in the above-mentioned label-free proteomic analysis. To quantify kinase activity changes in each comparison *(i.e.,* Ctrl *vs.* gRNA10_4 or Ctrl *vs.* gRNA10_12), the KSEA algorithm was employed as described^46^. Briefly, a correctly formatted dataset containing phosphosites with *P* < 0.05 were inputted into the KSEA App^28^ interface, where the PhosphoSitePlus plus NetworKIN dataset was selected. Only kinases with ≥ 5 substrate sites were selected for the quantification of kinase activities. For each kinase, the log2(Ctrl/KO) ratios of all substate sites were averaged to infer kinase activity changes.

### Western blot

Membranes were probed with antibodies against RIPK2 (1:1,000, Cell Signaling Technology #4142 or Santa Cruz Biotechnology #sc-166765), c-Myc (1:5,000, Abcam #ab32072), phospho-c-Myc (Ser62) (1:1,000, Cell Signaling Technology #13748S), ubiquitin (K48-linkage specific) (1:1,000, Cell Signaling Technology, #12805), FLAG (1:5,000, Sigma #F1804), MKK7 (1:1,000, Cell Signaling Technology #4172S or 1:2,000, Santa Cruz Biotechnology #sc-25288), MKK7 (phospho-Ser271) (1:1,000, Aviva Systems Biology #OAAF05547), JNK (1:1,000, Cell Signaling Technology #9252S or 1:2,000 Santa Cruz Biotechnology sc-7345), JNK (phospho-T183/Y185) (1:1,000, Cell Signaling Technology #9251S), mCherry (1:1,000, Abcam #ab213511), GFP (1:1,000, Abcam #ab290), β-actin (1:5,000, Sigma Aldrich #5441), or GAPDH (1:1,000, Cell Signaling Technology #3683). Signal was visualized with secondary HRP-conjugated antibodies (1:5,000, Cell Signaling Technology #7074S or #7076S) and chemiluminescent detection.

### c-Myc ubiquitination assay

Control and RIPK2-KO PC3 Cells were cultured in 100-mm dishes until reaching the confluency of ~80%, and then treated with 10 μM MG132 for 4 hours before cell lysis. Immunoprecipitation was performed as described in the Interactome analysis section, except that an anti-c-Myc antibody (Abcam #ab32072) was used. Eluted proteins were probed by Western Blot as indicated.

### Hematoxylin and eosin (H&E) staining and immunohistochemistry (IHC)

H&E and IHC were carried out by the Cedars-Sinai Biobank & Translational Pathology core by following standardized protocols. For IHC staining of tissue microarrays, the anti-RIPK2 antibody (Sigma Aldrich #HPA015273) was used at 1:100 dilution. Stained slides were digitized using Aperio AT Turbo (Leica Biosystems). Cancer areas and normal glands were annotated by an expert pathologist (X.Y.) in the images, which were then exported for image analysis in the Leica Tissue IA software package (Leica Biosystems). Protein expression was quantified by the mean 3,3’-diaminobenzidine (DAB) staining intensity of pixels in the annotated tumor areas. DAB staining was automatically deconvolved from hematoxylin by the software.

### Immunofluorescence and fluorescence imaging

For immunofluorescence imaging, 2 mL (1.25 × 10^5^/mL) of HEK293T cells with RIPK2-KO were plated into 6-well plates and grown on poly-L-lysine pre-treated coverslips. Cells were transfected with NES-RIPK2m4-3×FLAG or NLS-RIPK2m4-3×FLAG (1.0 μg per plasmid) for 48 h. Cells were then fixed and permeabilized in methanol/acetone (1:1, v/v) for 20 min at – 20°C, washed by PBS twice, and blocked in 2% BSA for 1 h at 37°C. The primary antibody against FLAG (1:500, Sigma Aldrich #F1804) was diluted in 2% BSA and incubated at 4°C overnight. Following 3× PBS washes, fluorochrome-conjugated secondary antibody (1:1,000, Cell Signaling Technology #4408) was diluted in 2% BSA and incubated with the samples for 1 h at 37°C in the dark. After 3× PBS washes, the coverslips were transferred to slides mounted in Mounting Medium with DAPI (Millipore, #DUO82040), and the cells were viewed under an All-in-one fluorescence microscope (BZ-X700, Keyence). For fluorescence imaging, 2 mL (1.25 × 10^5^/mL) of HEK293T cells with the knockout of both RIPK2 and MKK7 genes were plated and grown as mentioned above. Cells were co-transfected with 1) mCherry and eGFP, 2) MKK7-mCherry and eGFP, 3) mCherry and RIPK2-eGFP, or 4) MKK7-mCherry and RIPK2-eGFP (0.5 μg per plasmid) for 48 h. Cells were fixed and visualized as described in the immunofluorescence imaging.

### Proximity ligation assay (PLA)

PLA was performed according to the manufacturer’s instruction (Duolink, #DUO92101, Sigma). Primary antibodies were diluted in Duolink antibody diluent as follows: rabbit anti-RIPK2 (Cell Signaling Technology, #4142) at 1:500, mouse anti-MKK7 (Santa Cruz Biotechnology, #sc-25288) at 1:100, and mouse anti-PRKDC (Santa Cruz Biotechnology, #sc-5282) at 1:100. Imaging was performed with an All-in-one fluorescence microscope (BZ-X700, Keyence) under TexasRed and DAPI filters. The numbers of PLA signals (shown in red) and cells (nuclei, shown in blue) from four to five random fields were quantified for each sample with Image J (v1.52p)^47^.

### Structural modeling

The structural model of the RIPK2-MKK7 complex was generated from available structures in the RCSB Protein Data Bank (PDB)^48^. Briefly, the 3D structure of the RIPK2 kinase domain was used from an available structure (PDB code 6ES0)^49^, and that of the MKK7 kinase domain was generated by homology modelling^50^, using the structure of MEK1 (PDB code 1S9I)^51^ as a template. Putative dimerization models of kinase domains were generated with Rosetta^52^ and ZDOCK^53^. Top 10 models from each program were further subjected to molecular minimization and 1.2 ns short molecular dynamics using Desmond (Schrodinger, SBGrid Consortium^54^), in order to optimize molecular interaction and estimate energetics of the complex. Finally, the energetically stable protein complex was selected.

### Correlation analysis

To perform correlation analysis, we employed the PCTA prostate cancer (n=2,113)^6^, TCGA Firehose Legacy prostate cancer (n=499), and TCGA PanCancer Atlas^13^ (32 studies) cohort data. For gene expression abundance, PCTA provides median-centered and quantile normalized expression values^55^. For the TCGA Firehose Legacy prostate cancer cohort, median-centered log2(FPKM+1) values were computed for each gene. For the TCGA PanCancer Atlas, RSEM (RNA-Seq by Expectation Maximization) values were log-transformed using log2(RSEM+1) and median centered by genes. Given these expression values, gene-set activation score was computed by using weighted Z-score method^56^. Spearman’s method was used to compute correlation coefficients between genes and/or gene-sets, using the PCTA portal (thepcta.org) or RStudio (v1.2.5033).

### Statistical analysis

Statistical analyses were performed in RStudio (v1.2.5033). All statistical tests were two-sided with a significance level of *P* < 0.05. All statistics and reproducibility information are reported in the figure legends. For animal studies, sample sizes wer defined on the basis of past experience to achieve 80% power. For ethical reasons, the minimum number of animals necessary to achieve the scientific objectives was used.

### Reporting summary

Further information on research design is available in the Nature Research Reporting Summary linked to this paper.

## Data availability

The described label-free proteomics, interactome, phosphoproteomics data have been deposited in the PRIDE repository under the following accession numbers: PXD18890 (proteomics), PXD18870 (interactome), PXD18871 (phosphoproteomics).

## Acknowledgments

We thank Drs. Dolores Di Vizio, Jayoung Kim, Gina (Chia-Yi) Chu, Stephen Freedland, Hisashi Tanaka, and their group members for helpful suggestions. We are grateful to Dr. Mandana Zandian for lab assistance. We thank Dr. Robert Vessella for the tissue microarray that was supported by the Department of Defense Prostate Cancer Biorepository Network (PCBN) (W81XWH-14-2-0183). This work was supported in part by the Biobank and Translational Research Shared Resource of Cedars-Sinai Cancer. W.Y. was supported by NCI R01 (1R01CA218526, 1R01CA232574), Department of Defense Exploration-Hypothesis Development Award (W81XWH-15-1-0167), Cedars-Sinai Development of Prostate Cancer Fund, Cedars-Sinai Precision Health Award, and UCLA CTSI Core Voucher Award. Y.Y. was supported by the Department of Defense (DoD) – Early Investigator Research Award (W81XWH-18-1-0476).

## Author contributions

W.Y., Y.Y., and B.Z. conceived and designed the study. Y.Y. and B.Z. carried out most experiments. C.Q. contributed to the mouse experiments. A.V. contributed to Western blot and PCTA analyses. A.C. and R.M. performed structural modeling. X.Y., B.S.K., and A.G. contributed to the tissue microarray analysis. E.P., N.K., M.R.F. interpreted results and edited the manuscript. S.Y. contributed to the PCTA analysis. W.Y., Y.Y., and B.Z. performed most data analysis, generated figures, and wrote the manuscript. All authors made intellectual contributions and reviewed the manuscript.

## References

1. Bray, F. et al. Global cancer statistics 2018: GLOBOCAN estimates of incidence and mortality worldwide for 36 cancers in 185 countries. CA. Cancer J. Clin. 68, 394–424 (2018).

2. Small, E. J. Redefining hormonal therapy for advanced prostate cancer: Results from the LATITUDE and STAMPEDE studies. Cancer Cell 32, 6–8 (2017).

3. Nuhn, P. et al. Update on systemic prostate cancer therapies: Management of metastatic castrationresistant prostate cancer in the era of precision oncology. Eur. Urol. 75, 88–99 (2019).

4. Siegel, R. L., Miller, K. D. & Jemal, A. Cancer statistics, 2020. CA. Cancer J. Clin. 70, 7–30 (2020).

5. Cerami, E. et al. The cBio cancer genomics portal: An open platform for exploring multidimensional cancer genomics data. Cancer Discov. 2, 401–404 (2012).

6. You, S. et al. Integrated classification of prostate cancer reveals a novel luminal subtype with poor outcome. Cancer Res. 76, 4948–4958 (2016).

7. Nguyen, D. T. et al. Pharos: Collating protein information to shed light on the druggable genome. Nucleic Acids Res. 45, D995–D1002 (2017).

8. Grasso, C. S. et al. The mutational landscape of lethal castration-resistant prostate cancer. Nature 487, 239–43 (2012).

9. Robinson, D. et al. Integrative clinical genomics of advanced prostate cancer. Cell 161, 1215–1228 (2015).

10. Abida, W. et al. Genomic correlates of clinical outcome in advanced prostate cancer. Proc. Natl. Acad. Sci. U. S. A. 166, 11428–11436 (2019).

11. He, S. & Wang, X. RIP kinases as modulators of inflammation and immunity. Nat. Immunol. 19, 912–922 (2018).

12. Zare, A. et al. RIPK2: New elements in modulating inflammatory breast cancer pathogenesis. Cancers (Basel). 10, 1–17 (2018).

13. Weinstein, J. N. et al. The Cancer Genome Atlas Pan-Cancer analysis project. Nat. Genet. 45, 1113–1120 (2013).

14. Zhao, S. G. et al. Associations of luminal and basal subtyping of prostate cancer with prognosis and response to androgen deprivation therapy. JAMA Oncol. 3, 1663–1672 (2017).

15. Haile, P. A. et al. The identification and pharmacological characterization of 6-(tert -Butylsulfonyl)-N -(5-fluoro-1 H -indazol-3-yl)quinolin-4-amine (GSK583), a highly potent and selective inhibitor of RIP2 kinase. J. Med. Chem. 59, 4867–4880 (2016).

16. Canning, P. et al. Inflammatory signaling by NOD-RIPK2 is inhibited by clinically relevant type II kinase inhibitors. Chem. Biol. 22, 1174–1184 (2015).

17. Sanjana, N. E., Shalem, O. & Zhang, F. Improved vectors and genome-wide libraries for CRISPR screening. Nat. Methods 11, 783–784 (2014).

18. Doench, J. G. et al. Optimized sgRNA design to maximize activity and minimize off-target effects of CRISPR-Cas9. Nat. Biotechnol. 34, 184–191 (2016).

19. Brown, L. C., Lu, C., Antonarakis, E. S., Luo, J. & Armstrong, A. J. Androgen receptor variant-driven prostate cancer II: advances in clinical investigation. Prostate Cancer Prostatic Dis. (2020) doi:10.1038/s41391-020-0215-5.

20. Halabi, S. et al. Meta-Analysis evaluating the impact of site of metastasis on overall survival in men with castration-resistant prostate cancer. J. Clin. Oncol. 34, 1652–1659 (2016).

21. Zhang, F. et al. Whole-body physiologically based pharmacokinetic model for nutlin-3a in mice after intravenous and oral administration. Drug Metab. Dispos. 39, 15–21 (2011).

22. Van Riggelen, J., Yetil, A. & Felsher, D. W. MYC as a regulator of ribosome biogenesis and protein synthesis. Nat. Rev. Cancer 10, 301–309 (2010).

23. Dang, C. V. MYC on the path to cancer. Cell 149, 22–35 (2012).

24. Whitfield, J. R., Beaulieu, M. E. & Soucek, L. Strategies to inhibit Myc and their clinical applicability. Front. Cell Dev. Biol. 5, 1–13 (2017).

25. Liberzon, A. et al. The molecular signatures database hallmark gene set collection. Cell Syst. 1, 417–425 (2015).

26. El Gammal, A. T. et al. Chromosome 8p deletions and 8q gains are associated with tumor progression and poor prognosis in prostate cancer. Clin. Cancer Res. 16, 56–64 (2010).

27. Farrell, A. S. & Sears, R. C. MYC degradation. Cold Spring Harb. Perspect. Med. 4, 1–16 (2014).

28. Wiredja, D. D., Koyutürk, M. & Chance, M. R. The KSEA App: a web-based tool for kinase activity inference from quantitative phosphoproteomics. Bioinformatics 33, 3489–3491 (2017).

29. Noguchi, K. et al. Regulation of c-Myc through phosphorylation at Ser-62 and Ser-71 by c-Jun N-terminal kinase. J. Biol. Chem. 274, 32580–7 (1999).

30. Mathiasen, D. P. et al. Identification of a c-Jun N-terminal kinase-2-dependent signal amplification cascade that regulates c-Myc levels in ras transformation. Oncogene 31, 390–401 (2012).

31. Guan, B., Wang, T.-L. & Shih, I.-M. ARID1A, a factor that promotes formation of SWI/SNF-mediated chromatin remodeling, is a tumor suppressor in gynecologic cancers. Cancer Res. 71, 6718–27 (2011).

32. Sanjana, N. E., Shalem, O. & Zhang, F. Improved vectors and genome-wide libraries for CRISPR screening. Nat. Methods 11, 783–784 (2014).

33. Doench, J. G. et al. Optimized sgRNA design to maximize activity and minimize off-target effects of CRISPR-Cas9. Nat. Biotechnol. 34, 184–191 (2016).

34. Sancak, Y. et al. The Rag GTPases bind raptor and mediate amino acid signaling to mTORC1. Science 320, 1496–1501 (2008).

35. Jin, C. et al. Safe engineering of CAR T cells for adoptive cell therapy of cancer using long-term episomal gene transfer. EMBO Mol. Med. 8, 702–11 (2016).

36. Justus, C. R., Leffler, N., Ruiz-Echevarria, M. & Yang, L. V. In vitro cell migration and invasion assays. J. Vis. Exp. (2014) doi:10.3791/51046.

37. Borowicz, S. et al. The soft agar colony formation assay. J. Vis. Exp. e51998 (2014) doi:10.3791/51998.

38. Zhou, B. et al. Low-background acyl-biotinyl exchange largely eliminates the coisolation of non-S-acylated proteins and enables deep S-acylproteomic analysis. Anal. Chem. 91, 9858–9866 (2019).

39. Wiśniewski, J. R., Zougman, A., Nagaraj, N. & Mann, M. Universal sample preparation method for proteome analysis. Nat. Methods 6, 359–362 (2009).

40. Cox, J. et al. Andromeda: a peptide search engine integrated into the MaxQuant environment. J. Proteome Res. 10, 1794–1805 (2011).

41. Cox, J. & Mann, M. MaxQuant enables high peptide identification rates, individualized p.p.b.-range mass accuracies and proteome-wide protein quantification. Nat. Biotechnol. 26, 1367–72 (2008).

42. Vizcaíno, J. A. et al. 2016 update of the PRIDE database and its related tools. Nucleic Acids Res. 44, D447–56 (2016).

43. Tyanova, S. et al. The Perseus computational platform for comprehensive analysis of (prote)omics data. Nat. Methods 13, 731–740 (2016).

44. Benjamini, Y. & Hochberg, Y. Controlling the false discovery rate: a practical and powerful approach to multiple testing. J. R. Stat. Soc. Ser. B 289–300 (1995).

45. Yang, W. et al. Integration of proteomic and transcriptomic profiles identifies a novel PDGF-MYC network in human smooth muscle cells. Cell Commun. Signal. 12, 44 (2014).

46. Casado, P. et al. Kinase-substrate enrichment analysis provides insights into the heterogeneity of signaling pathway activation in leukemia cells. Sci. Signal. 6, 1–14 (2013).

47. Schneider, C. a, Rasband, W. S. & Eliceiri, K. W. NIH Image to ImageJ: 25 years of image analysis. Nat. Methods 9, 671–675 (2012).

48. Bernstein, F. C. et al. The protein data bank: A computer-based archival file for macromolecular structures. Arch. Biochem. Biophys. 185, 584–591 (1978).

49. Suebsuwong, C. et al. Activation loop targeting strategy for design of receptor-interacting protein kinase 2 (RIPK2) inhibitors. Bioorg. Med. Chem. Lett. 28, 577–583 (2018).

50. Arnold, K., Bordoli, L., Kopp, J. & Schwede, T. The SWISS-MODEL workspace: A web-based environment for protein structure homology modelling. Bioinformatics 22, 195–201 (2006).

51. Ohren, J. F. et al. Structures of human MAP kinase kinase 1 (MEK1) and MEK2 describe novel noncompetitive kinase inhibition. Nat. Struct. Mol. Biol. 11, 1192–1197 (2004).

52. Moretti, R., Lyskov, S., Das, R., Meiler, J. & Gray, J. J. Web-accessible molecular modeling with Rosetta: The Rosetta Online Server that Includes Everyone (ROSIE). Protein Sci. 27, 259–268 (2018).

53. Pierce, B. G. et al. ZDOCK server: Interactive docking prediction of protein-protein complexes and symmetric multimers. Bioinformatics 30, 1771–1773 (2014).

54. Morin, A. et al. Collaboration gets the most out of software. Elife 2013, 1–6 (2013).

55. You, S. et al. A systems approach to rheumatoid arthritis. PLoS One 7, e51508 (2012).

56. Levine, D. M. et al. Pathway and gene-set activation measurement from mRNA expression data: The tissue distribution of human pathways. Genome Biol. 7, R93 (2006).

