## Extended Data Figures for "RIPK2 Stabilizes c-Myc and is an Actionable Target for Inhibiting Prostate Cancer Metastasis"

[1]Department of Surgery, Cedars-Sinai Medical Center, Los Angeles, California, United States
[2]Department of Biomedical Sciences, Cedars-Sinai Medical Center, Los Angeles, California, United States
[3]Department of Pathology and Laboratory Medicine, Cedars-Sinai Medical Center, Los Angeles, California, United States
[4]Department of Medicine, Cedars-Sinai Medical Center, Los Angeles, California, United States
[5]Samuel Oschin Comprehensive Cancer Institute, Cedars-Sinai Medical Center, Los Angeles, California, United States
[6]Department of Urology, Icahn School of Medicine at Mount Sinai, New York, New York, United States
[7]Department of Medicine, University of California Los Angeles, Los Angeles, California, United States
[8]These authors contributed equally
[†]Present address: Department of Pathology, University of Utah, Salt Lake City, Utah, United States

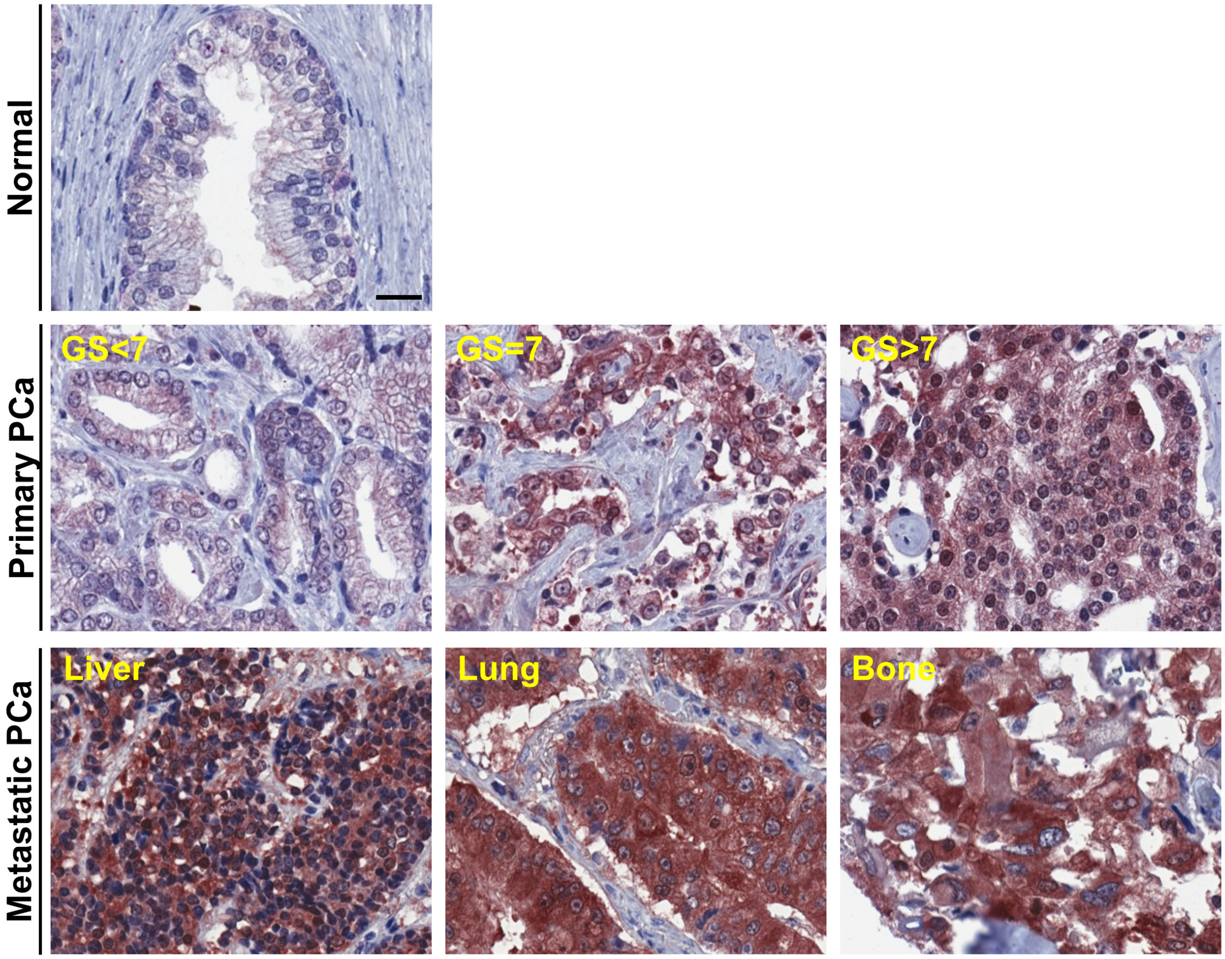


**Extended Data Fig. 1 | Representative immunohistochemical staining results of RIPK2 protein expression in human prostate cancer tissue microarrays.** Scale bar = 25 µm. GS, Gleason score; PCa, prostate cancer.

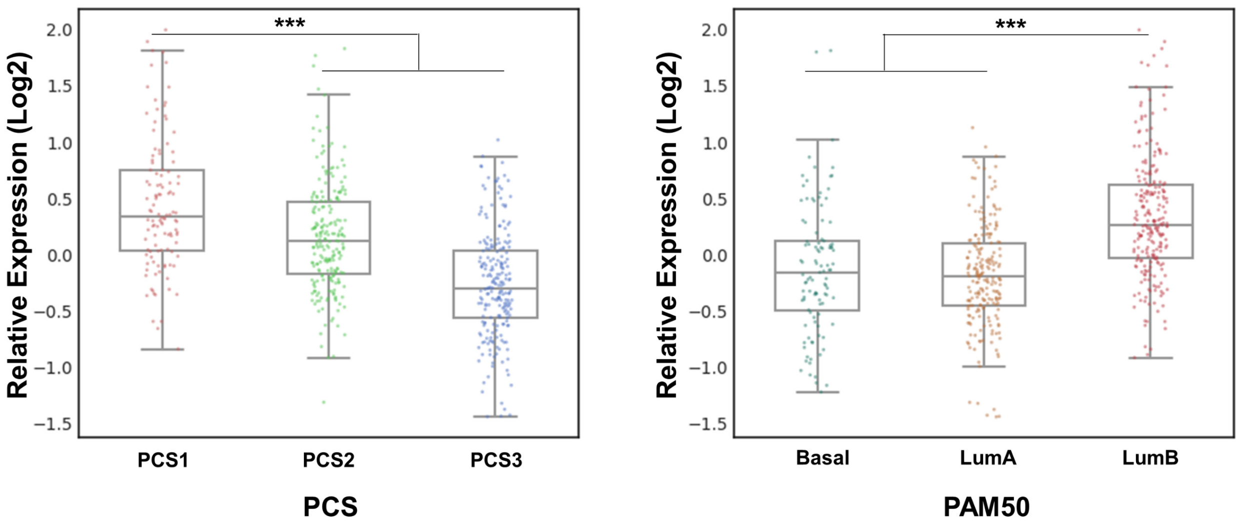


**Extended Data Fig. 2 | Boxplot of RIPK2 mRNA levels in three PCS (left) and PAM50 (right) subtypes of prostate cancer**. Here, the TCGA Firehose Legacy cohort (n=499) was used. *P* values were determined by unpaired two-tailed Student's *t*-test. *** $P < 0.001$.

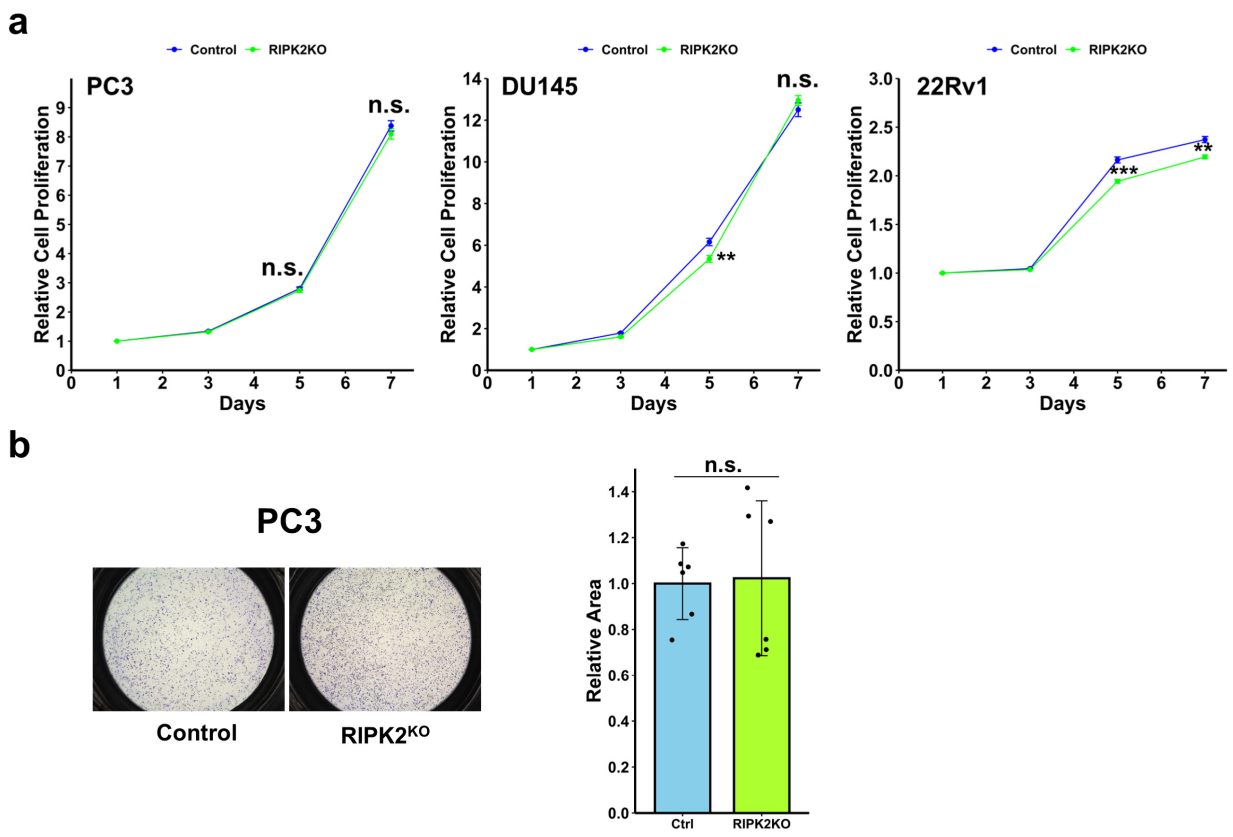


**Extended Data Fig. 3 | RIPK2-knockout (RIPK2$^{KO}$) has negligible effect on prostate cancer cell proliferation and migration. a,** Growth curves of prostate cancer PC3, DU145, and 22Rv1 cells with RIPK2$^{KO}$, compared with their respective control cells (n=10 for each cell line). Data were shown as mean ± standard deviation. Mann-Whitney *U* test; in day 5, PC3: $P$ = 0.28; DU145: $P$ = 3.89E-3; 22Rv1: $P$ = 1.68E-3; in day 7, PC3: $P$ = 0.31; DU145: $P$ = 0.17; 22Rv1: $P$ = 2.44E-4. **b,** Migration of control and RIPK2$^{KO}$ PC3 cells. Data were shown as mean ± standard deviation of six biological replicates. Unpaired two-tailed Student's *t*-test, $P$ = 0.88. In the figure, ** represents $P < 0.01$, *** $P < 0.001$, and n.s. not significant.

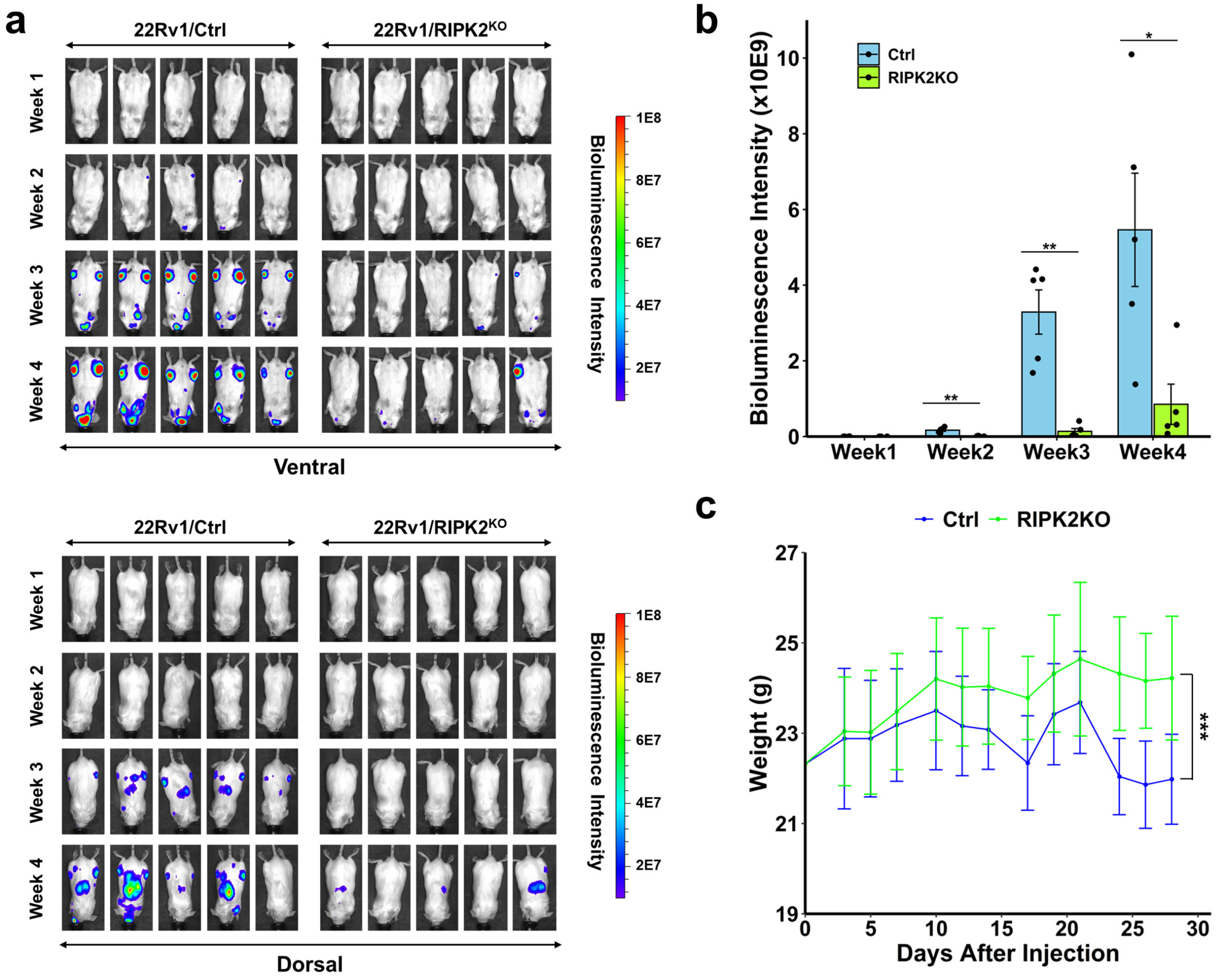


**Extended Data Fig. 4 | Comparison of mice bearing metastatic 22Rv1 control or RIPK2-KO tumors.** **a,** Bioluminescence images of mice (n=5 per group) from the ventral (top) and dorsal (bottom) sides at different weeks following intracardiac injection of control and RIPK2-KO 22Rv1 cells. **b,** Bar plot of the total bioluminescence intensities at different weeks after intracardiac injection. Data represent mean ± standard error of the mean. Mann-Whitney *U* test. * $P < 0.05$ and ** $P < 0.01$. **c,** Changes in mouse weight following intracardiac injection. Data represent mean ± standard error of the mean. Two-way ANOVA test. *** $P < 0.001$.

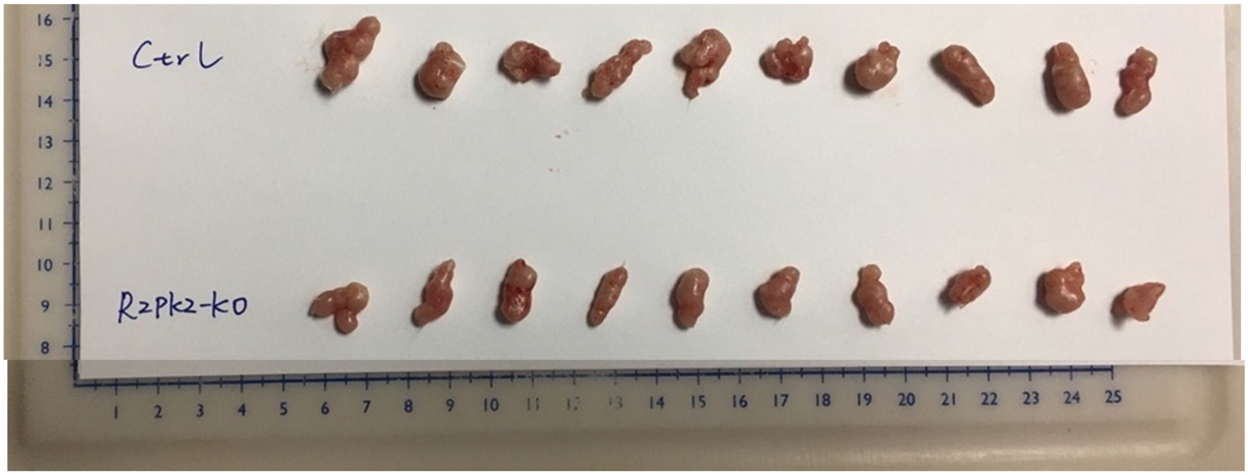


**Extended Data Fig. 5 | Image of 22Rv1 xenograft tumors.** The top row shows xenograft tumors derived from 22Rv1 cells transfected with a non-targeting control gRNA, while the bottom row shows xenograft tumors derived from 22Rv1 cells with RIPK2 knockout. Tumors were harvested 26 days after subcutaneous injection.

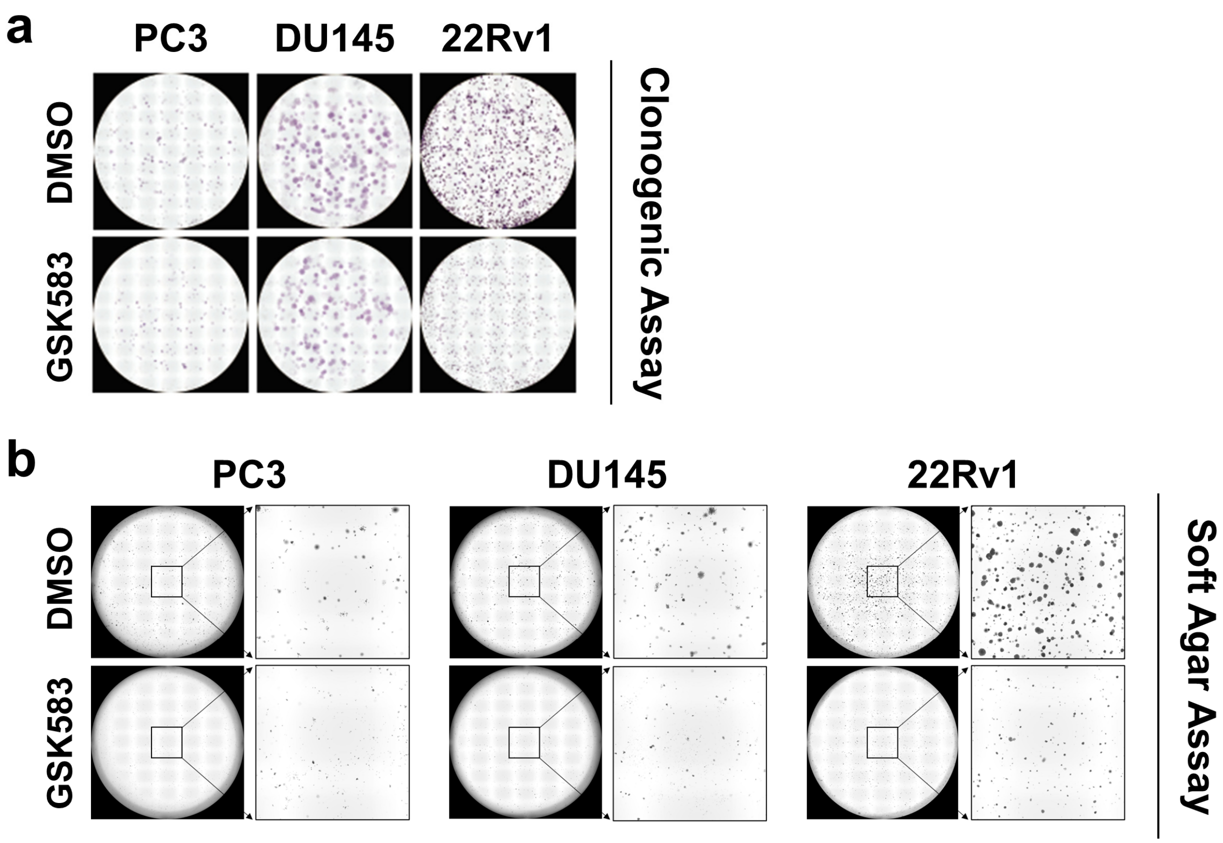


**Extended Data Fig. 6 | Effects of GSK583 treatment on the anchorage-dependent and -independent colony formation of prostate cancer PC3, DU145, and 22Rv1 cells. a,** Clonogenic assay to measure anchorage-dependent colony formation (n=6 for each cell line). **b,** Soft agar assay to measure anchorage-independent colony formation (n=6 for each cell line). See Fig. 2i and 2j for bar plots.

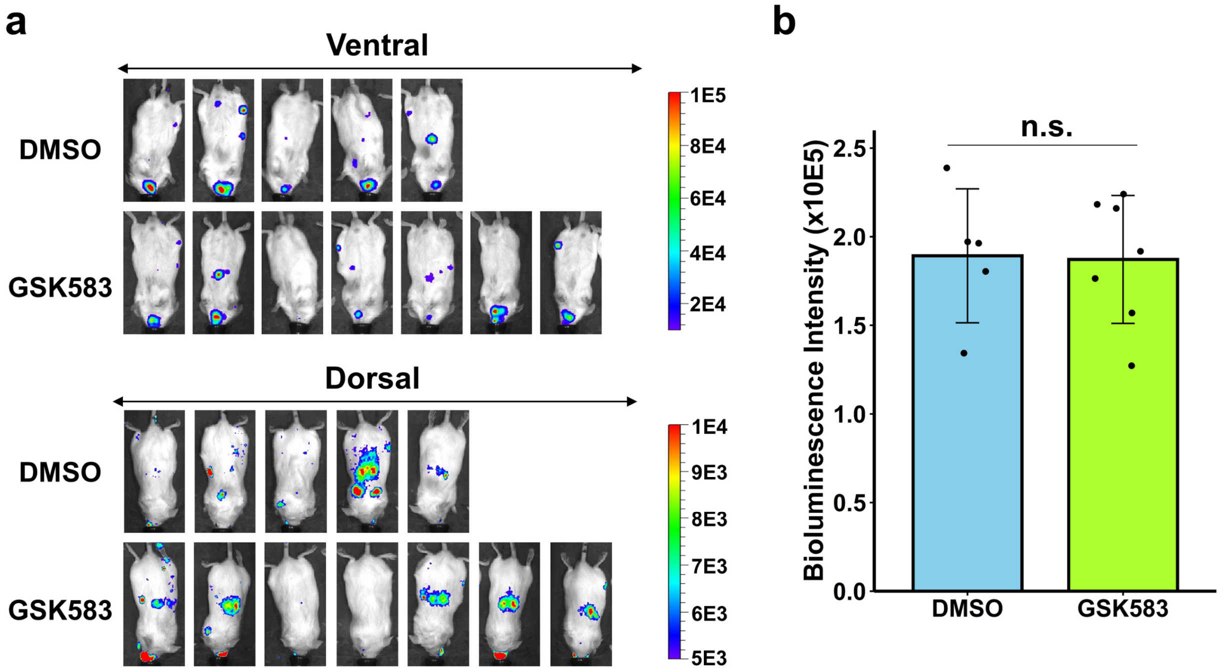


**Extended Data Fig. 7 | Comparison of the bioluminescence intensities in mice randomized into the control and GSK583 treatment groups. a,** Bioluminescence images of mice 7 days after intracardiac injection of luciferase-labeled 22Rv1 cells. **b,** Bar plot of the total bioluminescence intensities. Data represent the mean ± standard error of the mean. Mann-Whitney *U* test; $P = 0.88$; n.s. represents not significant.

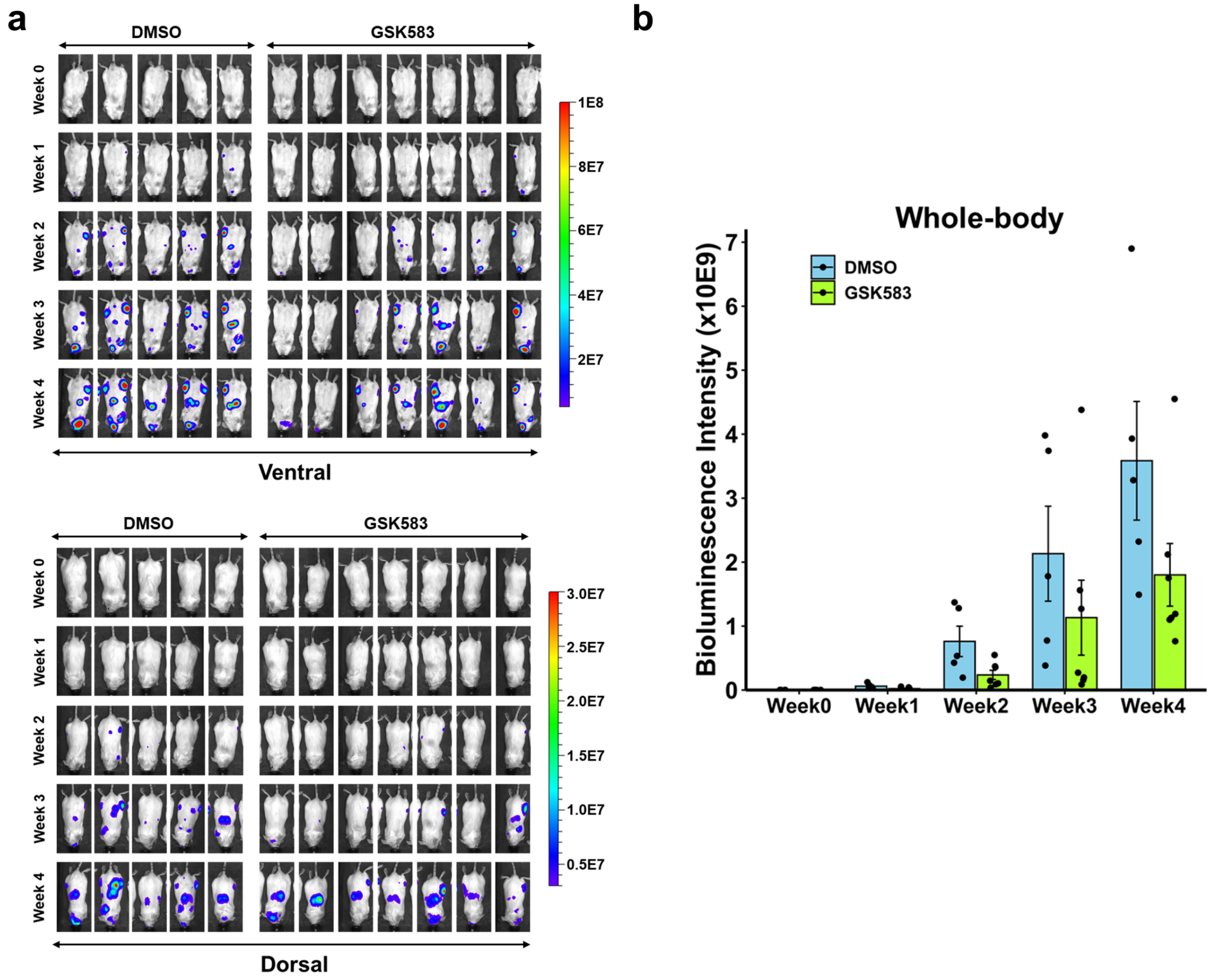


**Extended Data Fig. 8 | Bioluminescence imaging of mice bearing metastatic 22Rv1 tumors that were treated with GSK583 (10 mg per kg) or vehicle control. a,** Bioluminescence images of treated mice (vehicle, n=5; GSK583, n=7) from the ventral (top) and dorsal (bottom) sides at different weeks, starting at day 7 following intracardiac injection. **b,** Bar plot of the total bioluminescence intensities of treated mice at different weeks. Data represent the mean ± standard error of the mean.

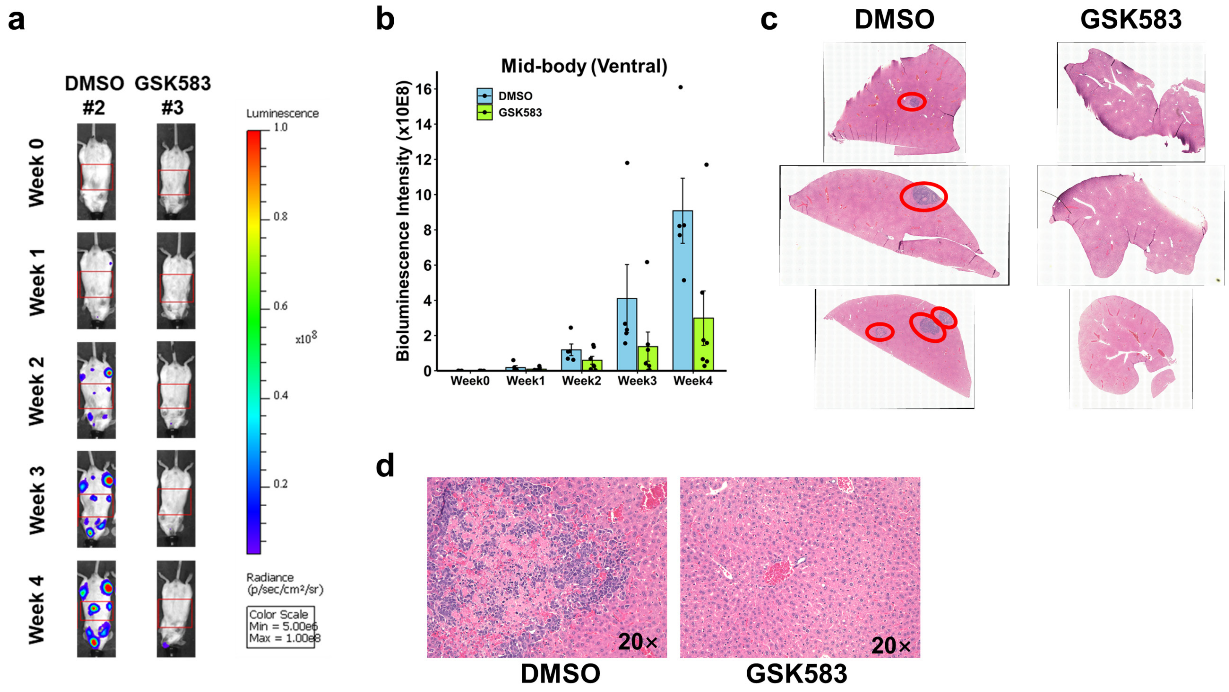


**Extended Data Fig. 9 | The effect of GKS583 treatment on liver metastasis. a,** Representative bioluminescence images of the mid-body sections (ventral side, in red boxes) of metastasis-bearing mice treated with GSK583 (10 mg per kg) or DMSO control. The mid-body sections are rich in liver metastasis. **b,** Bar plot of bioluminescence intensities of the mid-body sections of metastasis-bearing mice treated with GSK583, compared with vehicle control. Data represent the mean ± standard error of the mean. **c,** Representative hematoxylin and eosin (H&E)-stained images of livers from three GSK583-treated mice and three control mice. **d,** Representative 20× enlarged H&E images of livers from a GSK583-treated mouse and a control mouse.

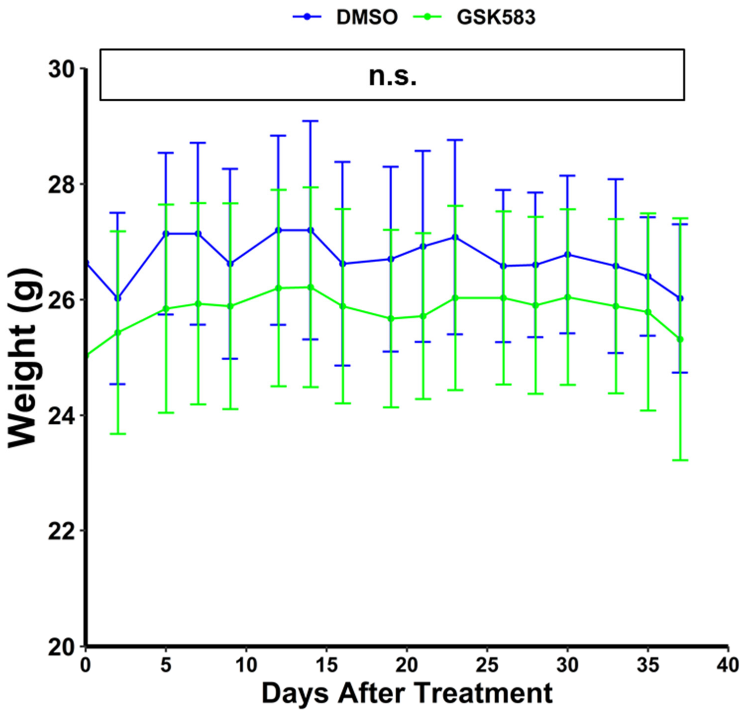


**Extended Data Fig. 10 | GSK583 treatment did not significantly affect mouse weight.** Data represent the mean ± standard deviation. Mann-Whitney *U* test; n.s. represents not significant.

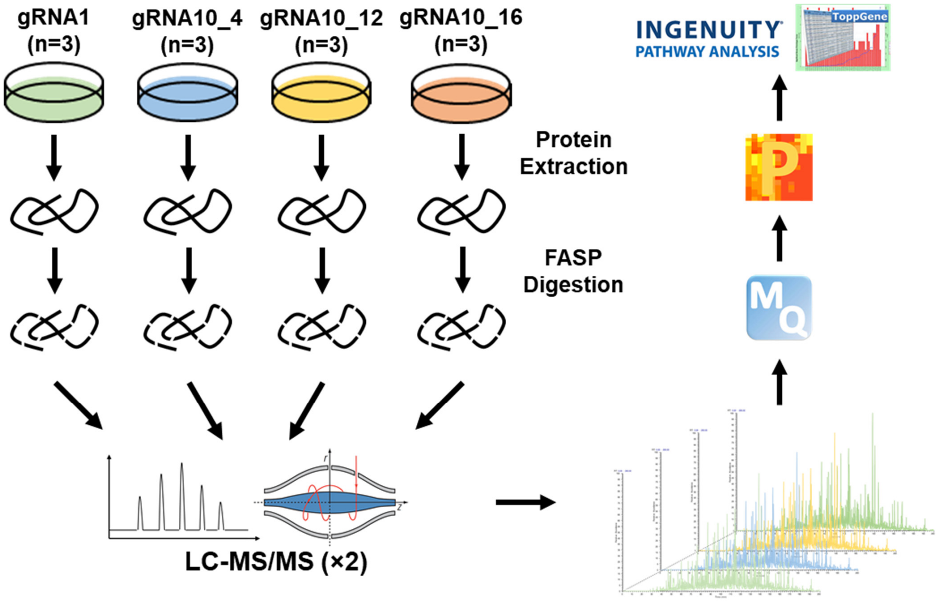


**Extended Data Fig. 11 | Schematic of label-free proteomics to identify proteins downstream of RIPK2.** Proteins were extracted from cells (three biological replicates in each group), reduced, alkylated, and digested into peptides by trypsin via filter-aided sample preparation (FASP). Tryptic peptides from each sample were analyzed by liquid chromatography-tandem mass spectrometry (LC-MS/MS) twice. The RAW files were processed using MaxQuant (MQ) for protein identification and quantification. The results were analyzed by Perseus (P) for quality assessment and statistical analysis. Identified differentially expressed proteins (DEPs) were analyzed by Ingenuity Pathway Analysis (IPA) to identify regulators upstream of the DEPs, and by ToppGene to identify significantly enriched gene ontology terms.

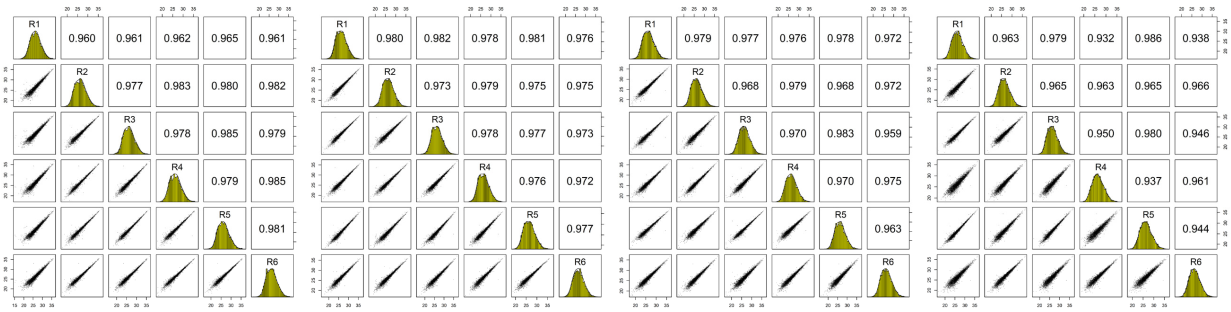


**PC3/Ctrl-gRNA** **PC3/gRNA10_4** **PC3/gRNA10_12** **PC3/gRNA10_16**

**Extended Data Fig. 12 | Pearson correlation coefficients between replicate runs of the label-free proteomics comparison of control and RIPK2-knockout PC3 cells.** PC3/Ctrl-gRNA: PC3 cells stably transfected with non-targeting control gRNA; PC3/gRNA10_4: PC3 cells stably transfected with RIPK2-targeting gRNA#10, single clone #4; PC3/gRNA10_12: PC3/gRNA10, single clone #12; PC3/gRNA10_16: PC3/gRNA10, single clone #16. Numbers in boxes: Pearson correlation coefficients; R1-R6: replicates #1-#6; scatter plots: distribution of log2-transformed label-free quantification (LFQ) intensities of protein groups.

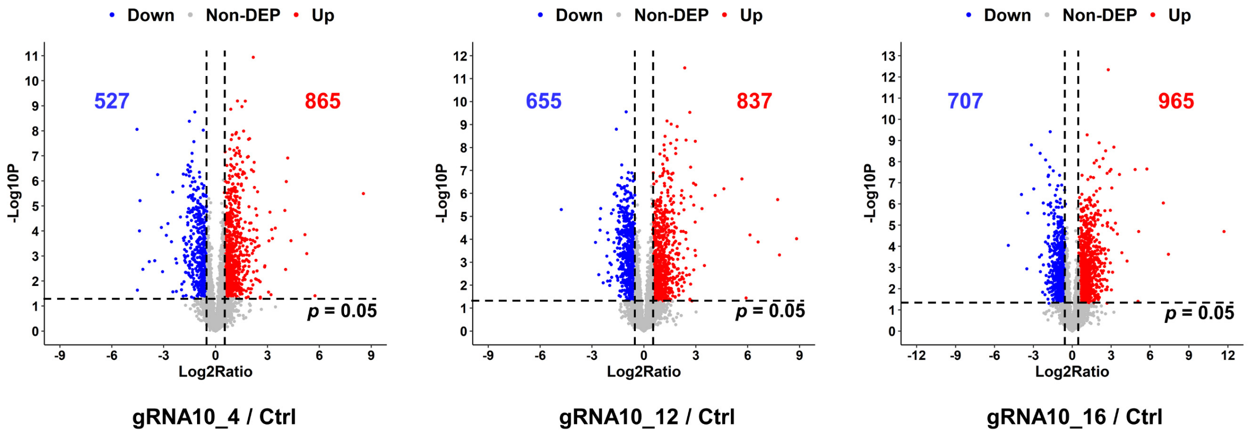


**Extended Data Fig. 13 | Volcano plots of protein groups quantified by label-free proteomics comparison of control and RIPK2-knockout PC3 cells.** Each dot represents a protein group. Blue dots represent protein groups significantly downregulated ($p < 0.05$, $\log_2$Ratio < -0.5) after RIPK2-knockout, compared with control cells. Red dots represent protein groups significantly upregulated ($p < 0.05$, $\log_2$Ratio > 0.5). Gray dots represent protein groups that were not significantly changed ($p \geq 0.05$ or $-0.5 \leq \log_2$Ratio $\leq 0.5$). The numbers in blue and red stand for the numbers of significantly downregulated and upregulated protein groups, respectively.

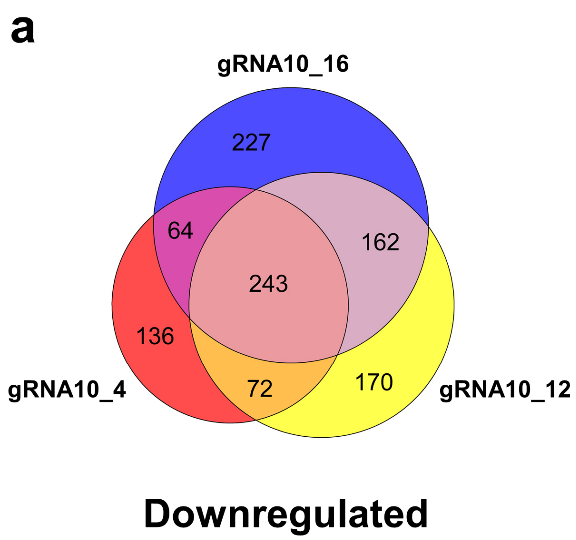


**Extended Data Fig. 14 | Venn diagrams of protein groups differentially expressed in three RIPK2-knockout PC3 clones, compared with control PC3 cells. a,** Venn diagram of protein groups significantly downregulated ($p < 0.05$, $\log_2$Ratio < -0.5) in the three PC3/RIPK2$^{KO}$ clones, compared with control PC3 cells. **b,** Venn diagram of protein groups significantly upregulated ($p < 0.05$, $\log_2$Ratio > 0.5) in the three PC3/RIPK2$^{KO}$ clones, compared with control PC3 cells.

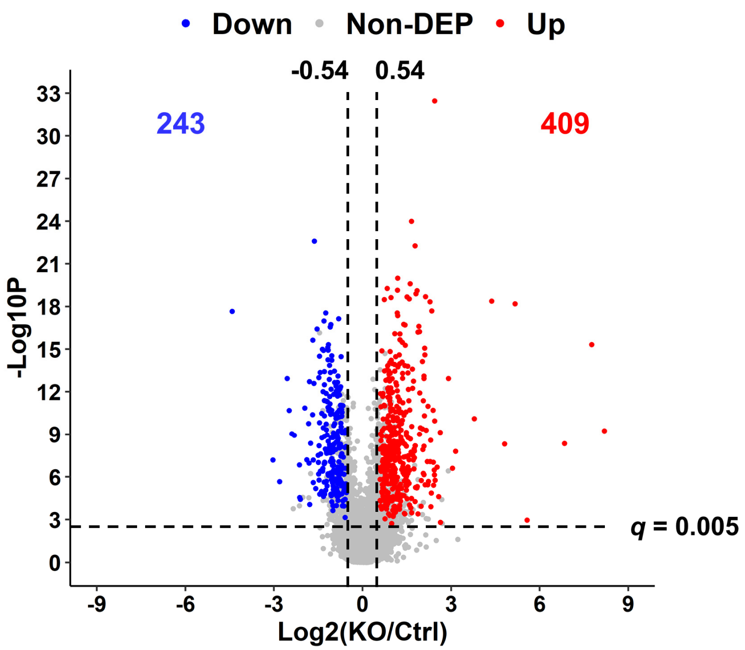


**Extended Data Fig. 15 | Volcano plot of the 652 protein groups that were consistently differentially expressed in the three PC3/RIPK2$^{KO}$ clones, compared with control PC3 cells.** Each dot represents a protein group. The $\log_2$(KO/Ctrl) ratios represent the mean values of $\log_2$Ratios in the three PC3/RIPK2$^{KO}$ clones compared with control cells. KO, RIPK2-knockout; Ctrl, control. *P* values represent the combined *p* values of the three comparisons shown in Extended Data Fig. 13. All of the 652 differentially expressed proteins have *q* values of $< 0.005$ and mean $\log_2$(KO/Ctrl) of $> 0.54$ in absolute value.

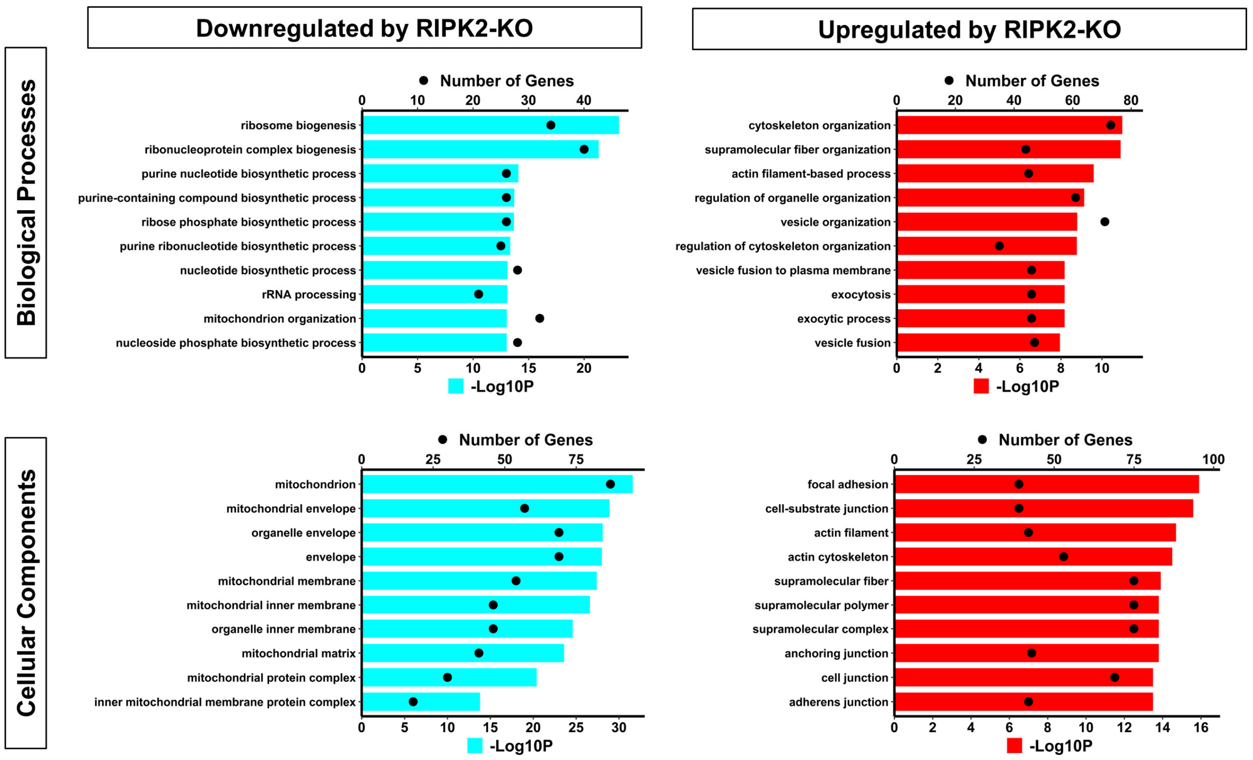


**Extended Data Fig. 16 | Bar graphs of the top 10 significantly enriched gene ontology terms.** The "Number of Genes", shown in black dots, represents the number of differentially expressed proteins corresponding to each gene ontology term. The bar colors represent downregulation (Cyan) and upregulation (red), and bar lengths correspond to the -$Log_{10}P$ values for the corresponding gene ontology terms.

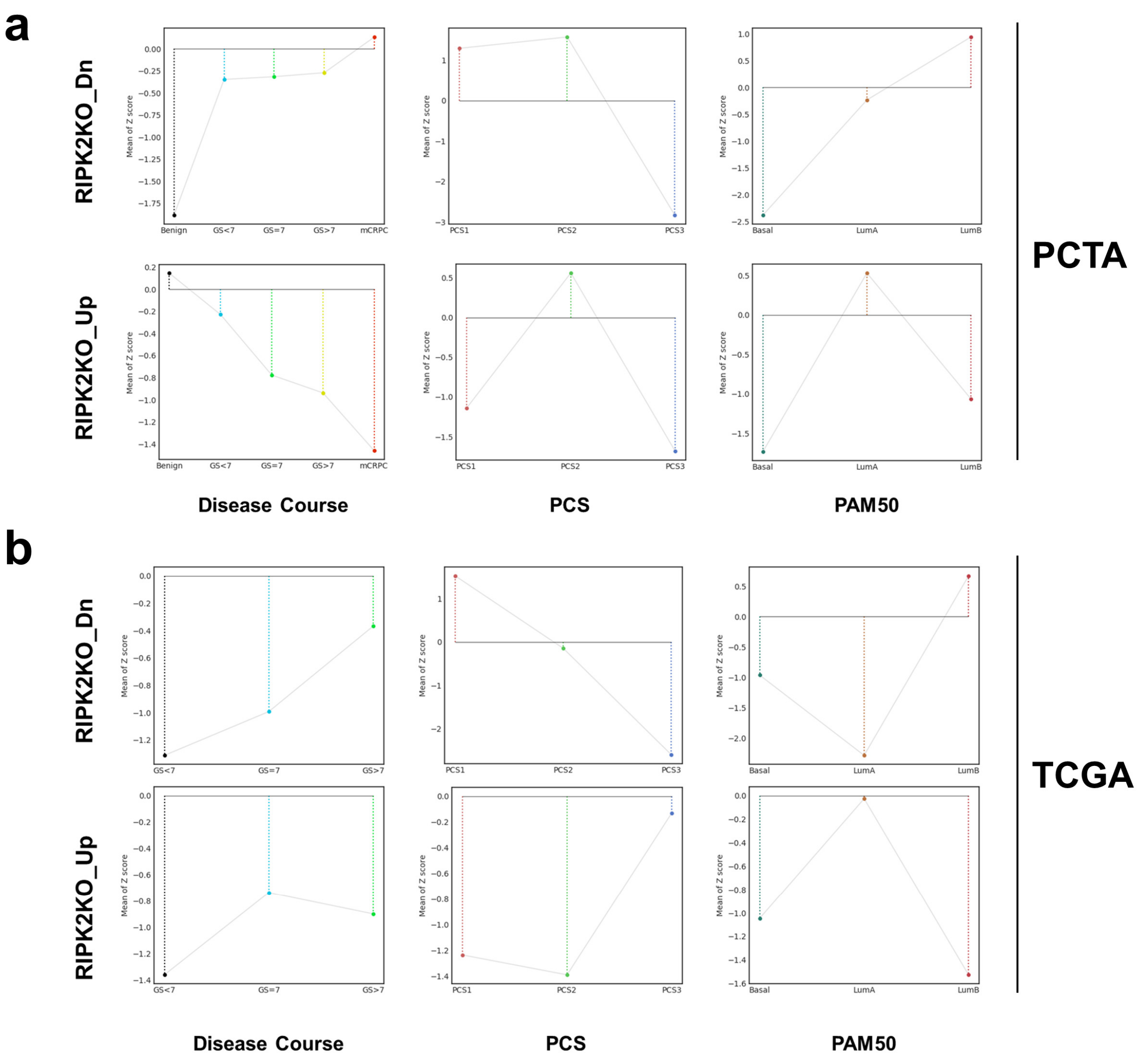


**Extended Data Fig. 17 | Line plots of the mean Z scores of the protein set downregulated or upregulated by RIPK2-knockout (RIPK2-KO). a,** Line plots for the PCTA cohort. **b,** Line plots for the TCGA Firehose Legacy prostate cancer cohort. RIPK2KO_Dn (RIPK2-induced): 243-protein set significantly downregulated by RIPK2-KO; RIPK2_Up (RIPK2-repressed): 409-protein set significantly upregulated by RIPK2-KO; GS, Gleason score; mCRPC: metastatic/castration-resistant prostate cancer; PCS: prostate cancer subtype; PAM50: prediction analysis of microarray 50.

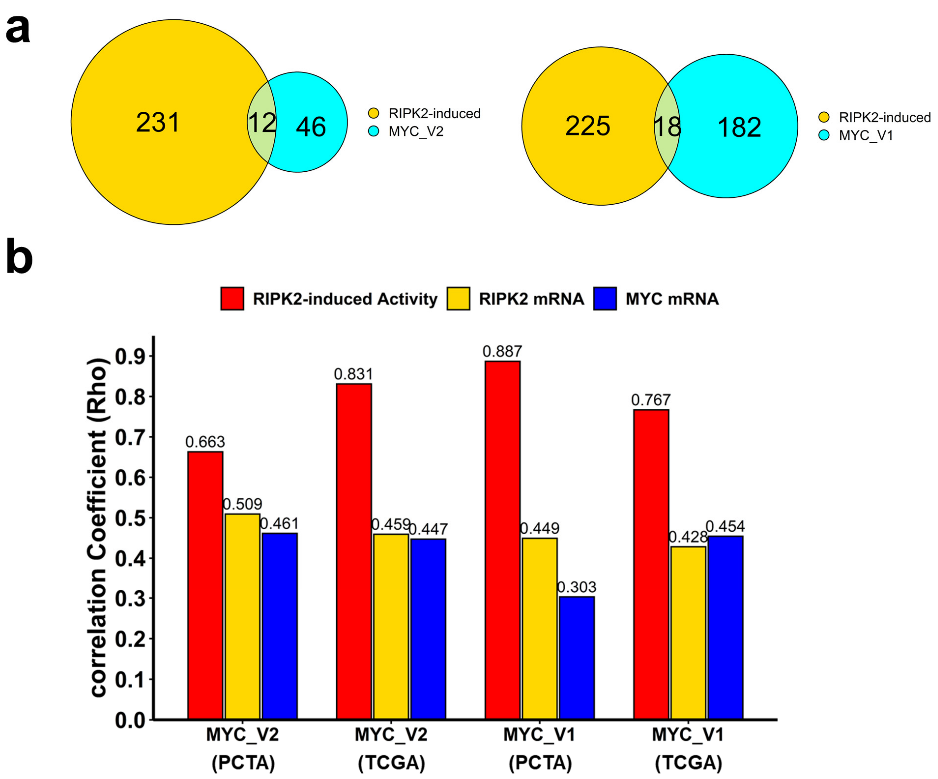


**Extended Data Fig. 18 | Comparison of the correlation between RIPK2-induced activity, RIPK2 mRNA, MYC mRNA, and MYC activity levels. a,** Venn diagrams of the genes contained in the RIPK2-induced signature and those in a Hallmark_MYC_Targets signature (V2, left; V1, right). **b,** Bar plot of the correlation coefficients (Spearman's Rho) in both PCTA and TCGA (Firehose Legacy) prostate cancer cohorts.

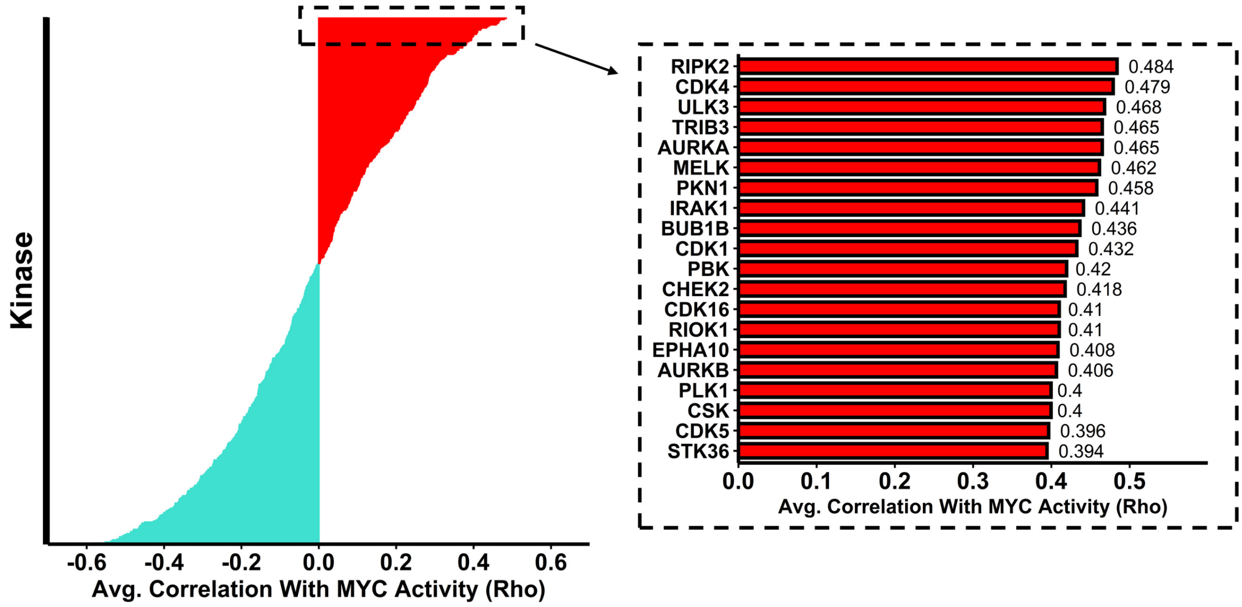


**Extended Data Fig. 19 | Bar graph of the average correlation coefficients (Spearman's rho) of RIPK2 mRNA levels with the human kinome in the PCTA and TCGA (Firehose Legacy) cohorts.** Red bars represent positive correlations, whereas turquoise bars represent negative correlations. Numbers adjacent to bars show the average Spearman's rho values in the two prostate cancer cohorts.

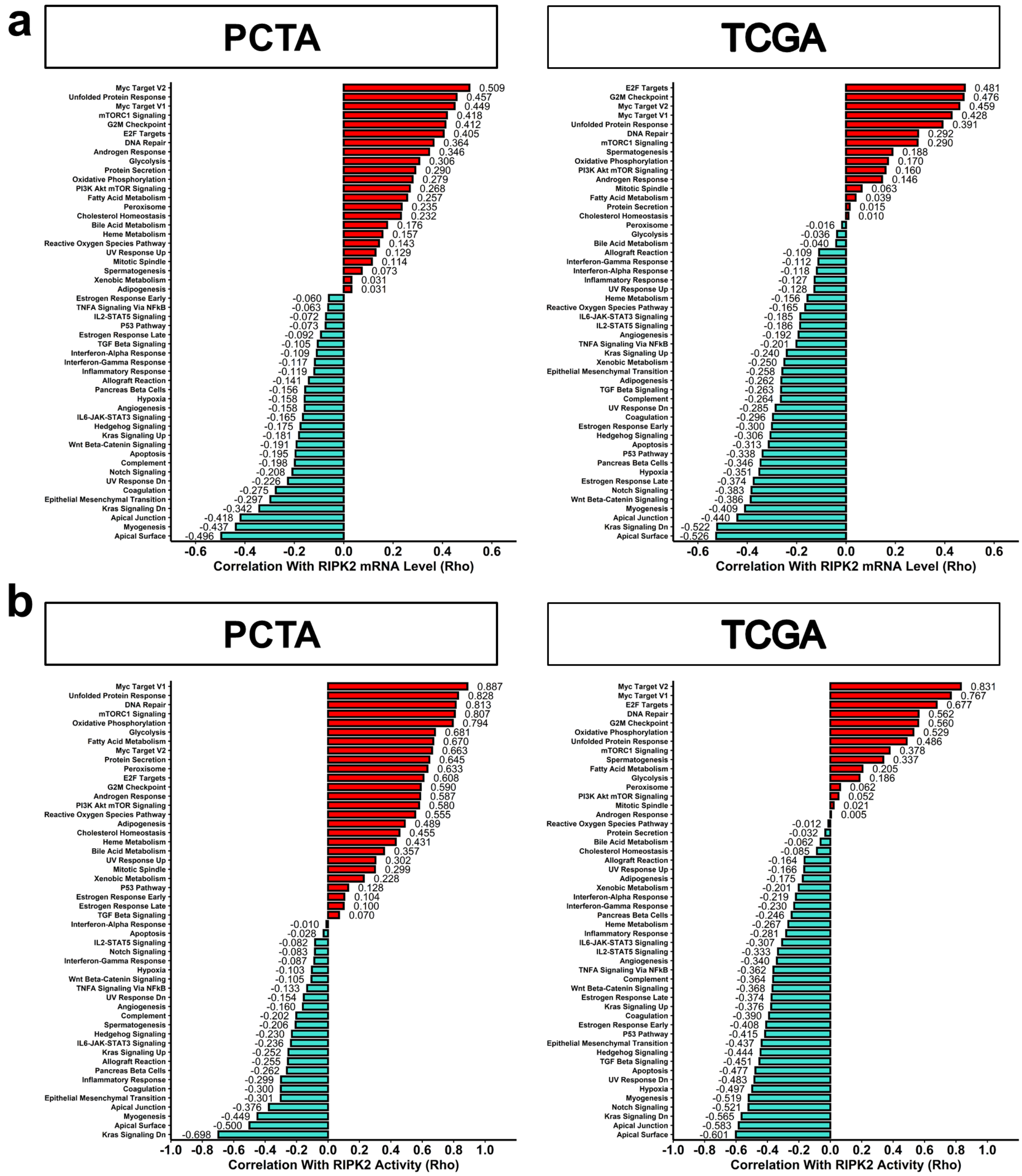


**Extended Data Fig. 20 | Bar graphs of the correlation coefficients (Spearman's rho) with the activity scores of the 50 MSigDB Hallmark gene sets. a,** Correlation with RIPK2 mRNA levels. **b,** Correlation with RIPK2-induced activity scores. Red bars represent positive correlations, whereas turquoise bars represent negative correlations. Numbers adjacent to bars show Spearman's rho values.

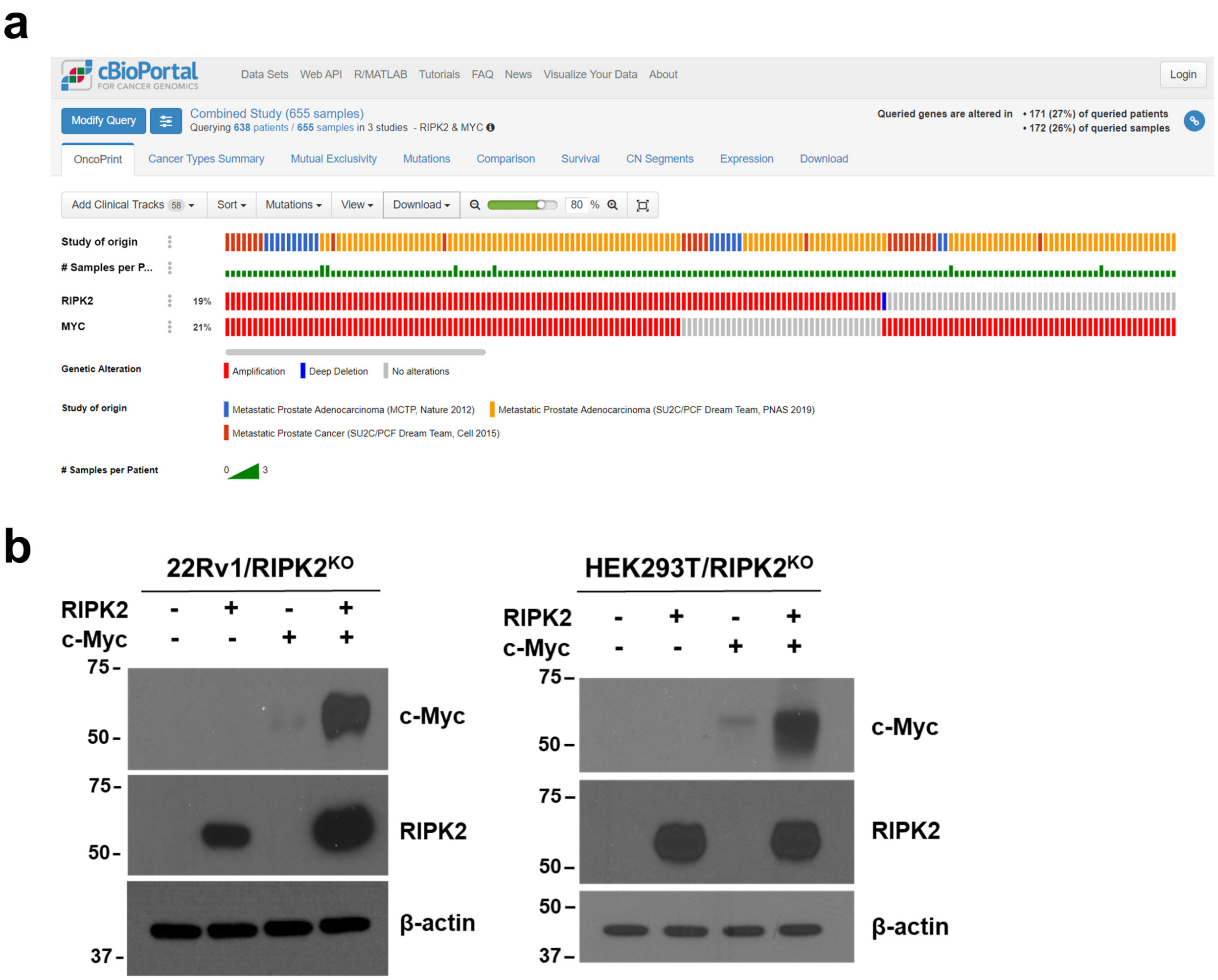


**Extended Data Fig. 21 | RIPK2 and MYC co-amplification and co-expression. a,** Frequency of RIPK2 and MYC amplification in three cohorts of metastatic castration-resistant prostate cancer (mCRPC) tissue specimens (n=655). **b,** Different levels of c-Myc protein increase in RIPK2-KO 22Rv1 (left) and HEK293T (right) cells in response to the ectopic expression of RIPK2 (0.5 μg plasmid) and/or c-Myc (0.5 μg plasmid).

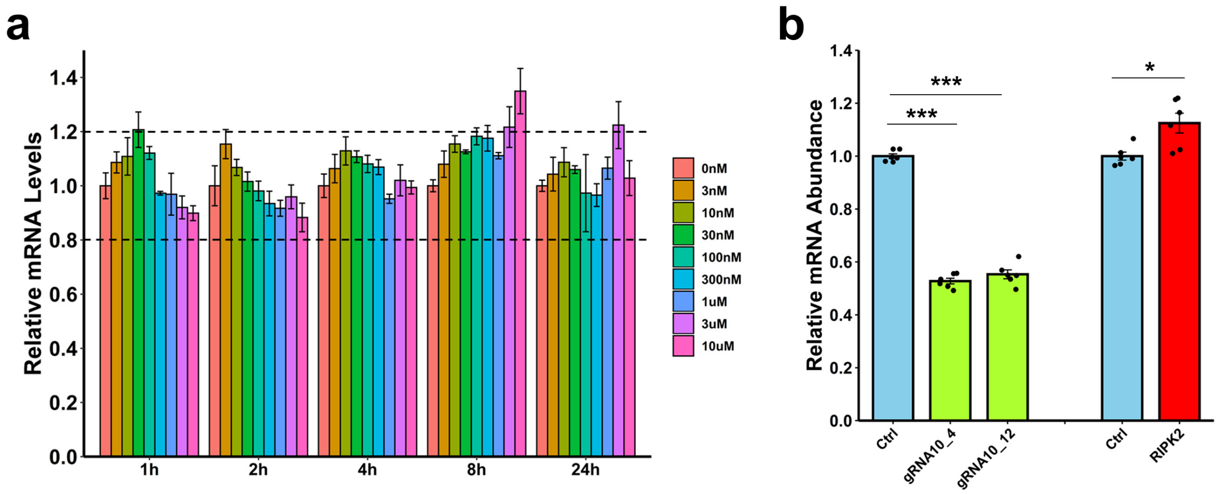


**Extended Data Fig. 22 | The effects of RIPK2 manipulation on c-Myc mRNA levels. a,** The changes of relative c-Myc mRNA abundance by different doses (0-10 μM) of GSK583 for different time (1-24 h). The data were normalized against GAPDH and then the control condition (*i.e.*, 0 nM) at each time point. **b,** The changes of relative c-Myc mRNA abundance by RIPK2 knockout or overexpression. The data were normalized against GAPDH and then the respective control conditions.

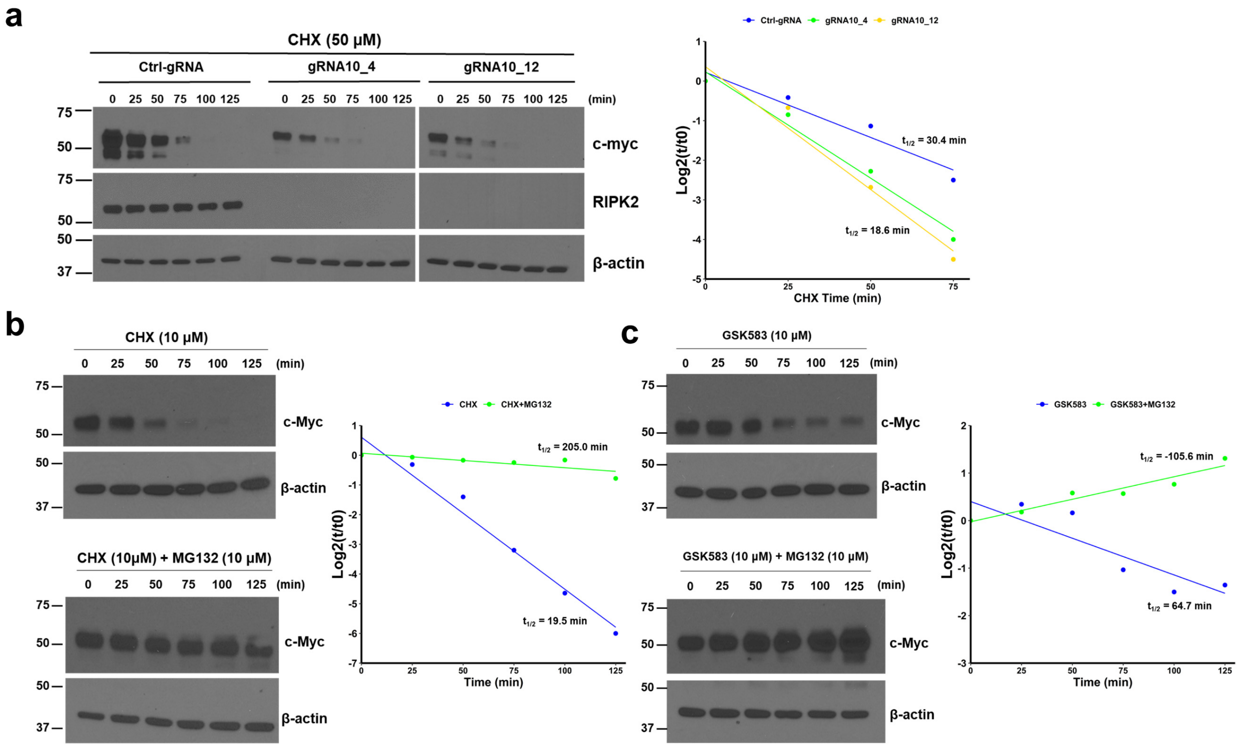


**Extended Data Fig. 23 | Genetic and pharmacologic inhibition of RIPK2 promotes the proteasomal degradation of c-Myc. a,** The c-Myc protein was degraded faster and had a shorter half-life in stable PC3/RIPK2$^{KO}$ single-cell clones #4 and #10, compared with control PC3 cells. CHX (cycloheximide) is a widely used protein translation inhibitor. **b,** c-Myc degradation in PC3 cells can be blocked by MG132, a widely used proteasomal degradation inhibitor. **c,** GSK583-induced c-Myc degradation in PC3 cells can be blocked by MG132.

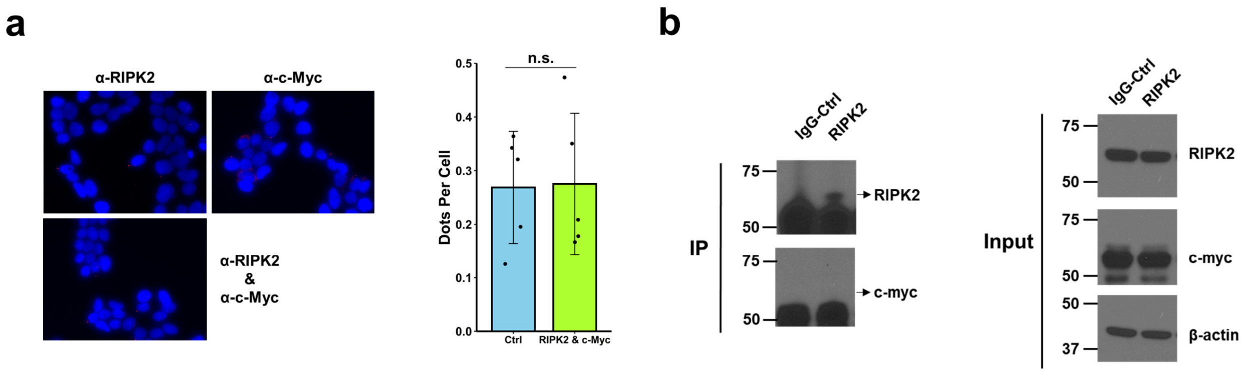


**Extended Data Fig. 24 | No protein-protein association of RIPK2 and c-Myc was detected. a,** Proximity ligation assay of RIPK2 and c-Myc proteins in parental HEK293T cells. In the bar graph, data represent the mean ± standard deviation and n.s. stands for not significant. The numbers of dots per cell under the control condition are the sum of dots per cell detected using an anti-RIPK2 or an anti-c-Myc antibody alone. **b,** Western blotting analysis of RIPK2 and c-Myc in immunoprecipitated RIPK2 protein complex from parental PC3 cells (left) or whole PC3 cell lysates (right).

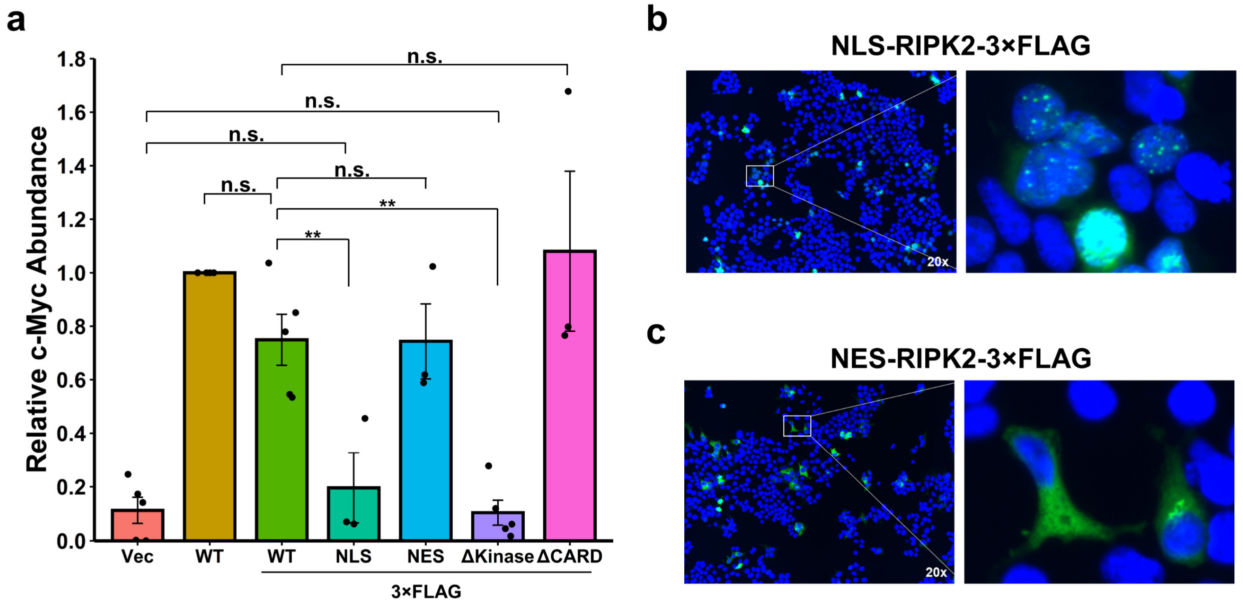


**Extended Data Fig. 25 | Ectopic expression of different constructs in RIPK2-KO HEK293T cells. a,** Densitometric quantification of the relative c-Myc protein abundance under different conditions. See Fig. 4a for representative Western blotting images. Data were first normalized against β-actin and then against the untagged wild-type (WT) condition and presented as mean ± standard error of the mean. Unpaired two-tailed Student's *t*-test. ** $P < 0.01$; n.s.: not significant. Vec, vector control; WT, wild-type RIPK2; NLS, nuclear localization signal-tagged RIPK2; NES, nuclear export signal-tagged RIPK2; ΔKinase, kinase domain-deleted RIPK2; ΔCARD, CARD domain-deleted RIPK2; 3×FLAG, with a C-terminal 3×FLAG tag. **b,** Immunofluorescence imaging analysis to confirm the nuclear localization of NLS-tagged RIPK2. **c,** Immunofluorescence imaging analysis to confirm the cytoplasmic localization of NES-tagged RIPK2.

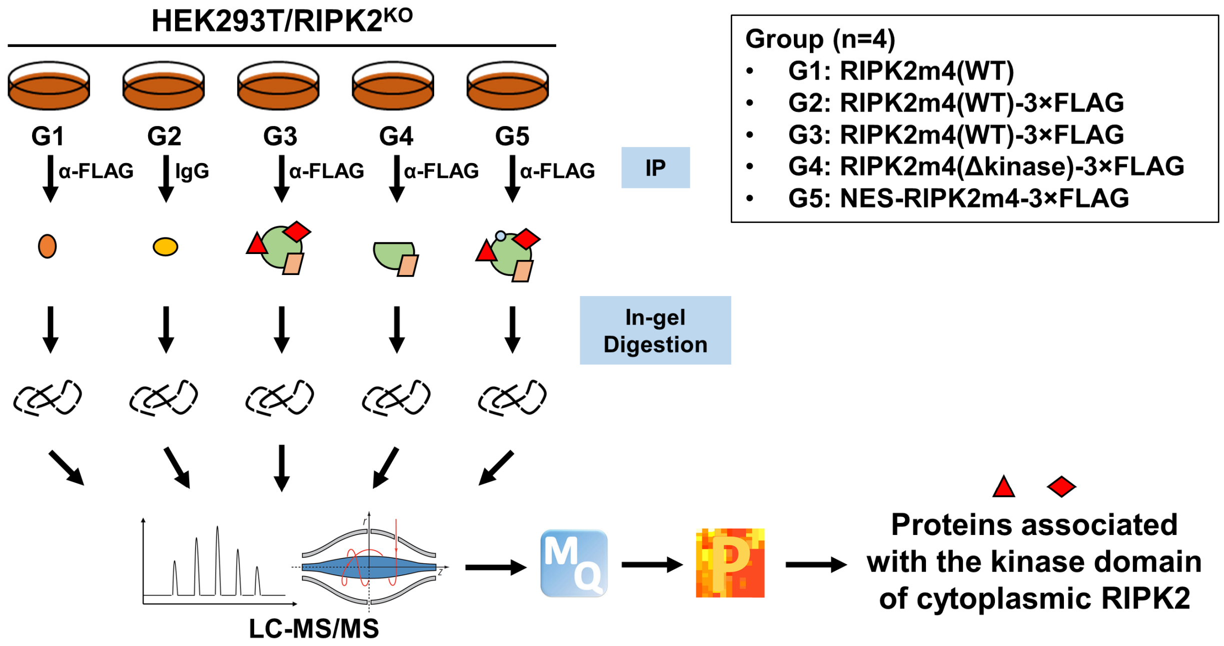


**Extended Data Fig. 26 | Schematic of RIPK2 interactome analysis.** Proteins were extracted from five groups of genetically manipulated HEK293T cells (n=4 per group). Among these, the G1, G2, and G4 groups were control groups: G1 was used to control for proteins associated with protein A/G-coated agarose beads and anti-FLAG antibody; G2 was used to control for proteins associated with protein A/G-coated agarose beads and protein alterations caused by ectopic expression of RIPK2 (wild-type)-3×FLAG; G4 was used to control for the identification of proteins associated with non-kinase-domain regions of RIPK2. The G3 and G5 groups were experimental groups: G3 was used to identify proteins associated with wild-type RIPK2; G5 was used to identify proteins associated with the cytoplasmic pool of RIPK2. Extracted proteins were subjected to immunoprecipitation using an anti-FLAG antibody (for G1, G3, G4, and G5 groups) or an IgG control (for the G2 group). Immunoprecipitated proteins went through in-gel protein reduction, alkylation, and digestion. Tryptic peptides were analyzed by liquid chromatography-tandem mass spectrometry (LC-MS/MS). The resulting RAW files were analyzed by MaxQuant (MQ) for protein identification and label-free quantification. Subsequently, Perseus was used to perform statistical analysis to identify significantly enriched proteins. IP: immunoprecipitation. WT: wild-type; Δkinase: deletion of the kinase domain; NES: nuclear export signal.

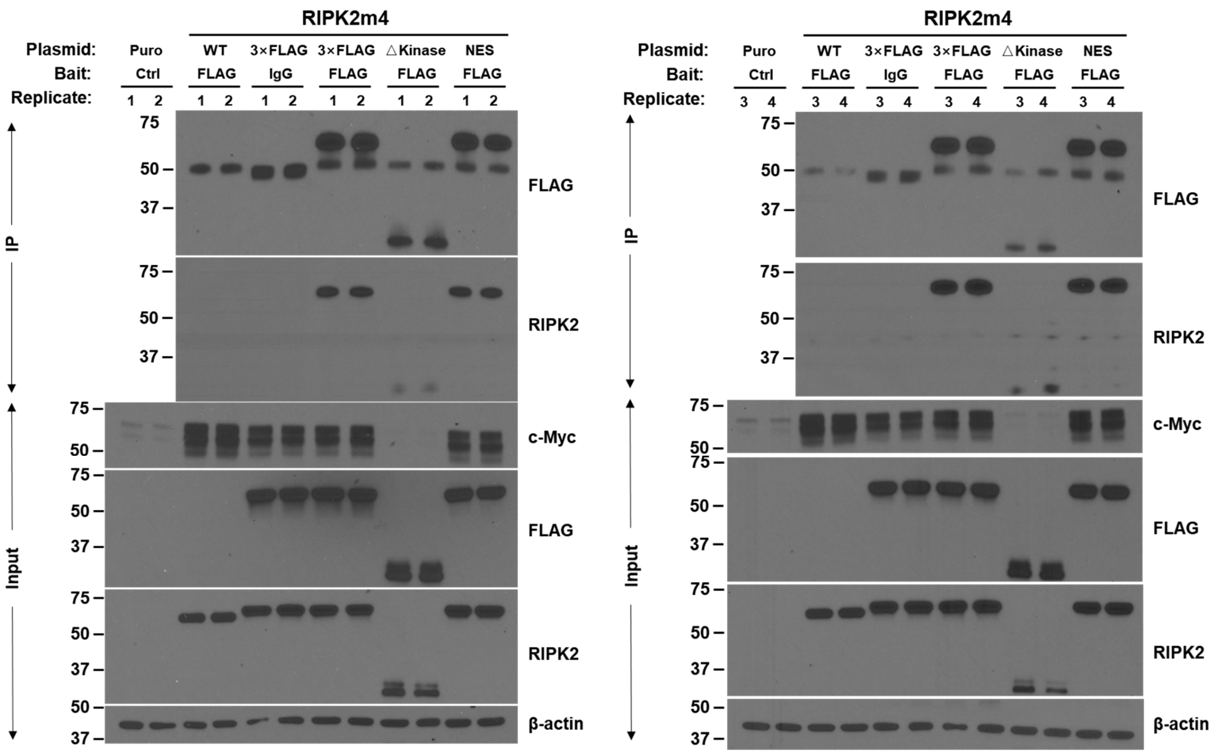


**Extended Data Fig. 27 | Western blot confirmation of successful immunoprecipitations.** HEK293T cells were transfected with empty vector, wild-type untagged RIPK2m4, 3×FLAG-tagged wild-type RIPK2m4, 3×FLAG-tagged NES-RIPK2m4, or 3×FLAG-tagged RIPK2m4-△Kinase. Protein complexes were immunoprecipitated with an anti-FLAG antibody or normal mouse IgG, followed by Western blotting analysis using indicated antibodies. See the Extended Data Fig. 26 for the experimental design.

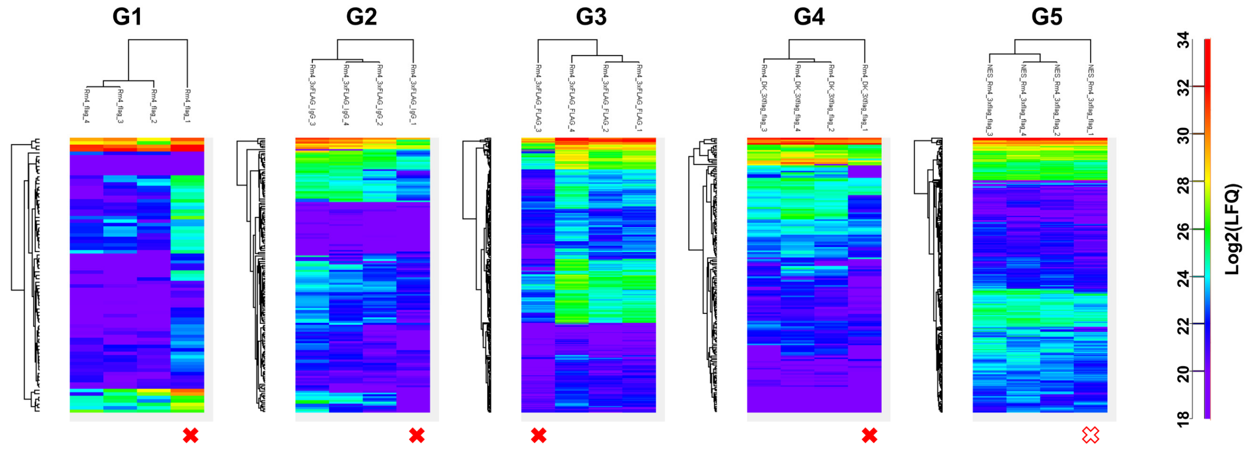


**Extended Data Fig. 28 | Determination of outlier RIPK2-interactome samples by unsupervised clustering of the log2-transformed LFQ intensities.** The outlier samples were excluded from further statistical analysis. G1-G5 corresponds to the G1-G5 groups shown in Extended Data Fig. 26.

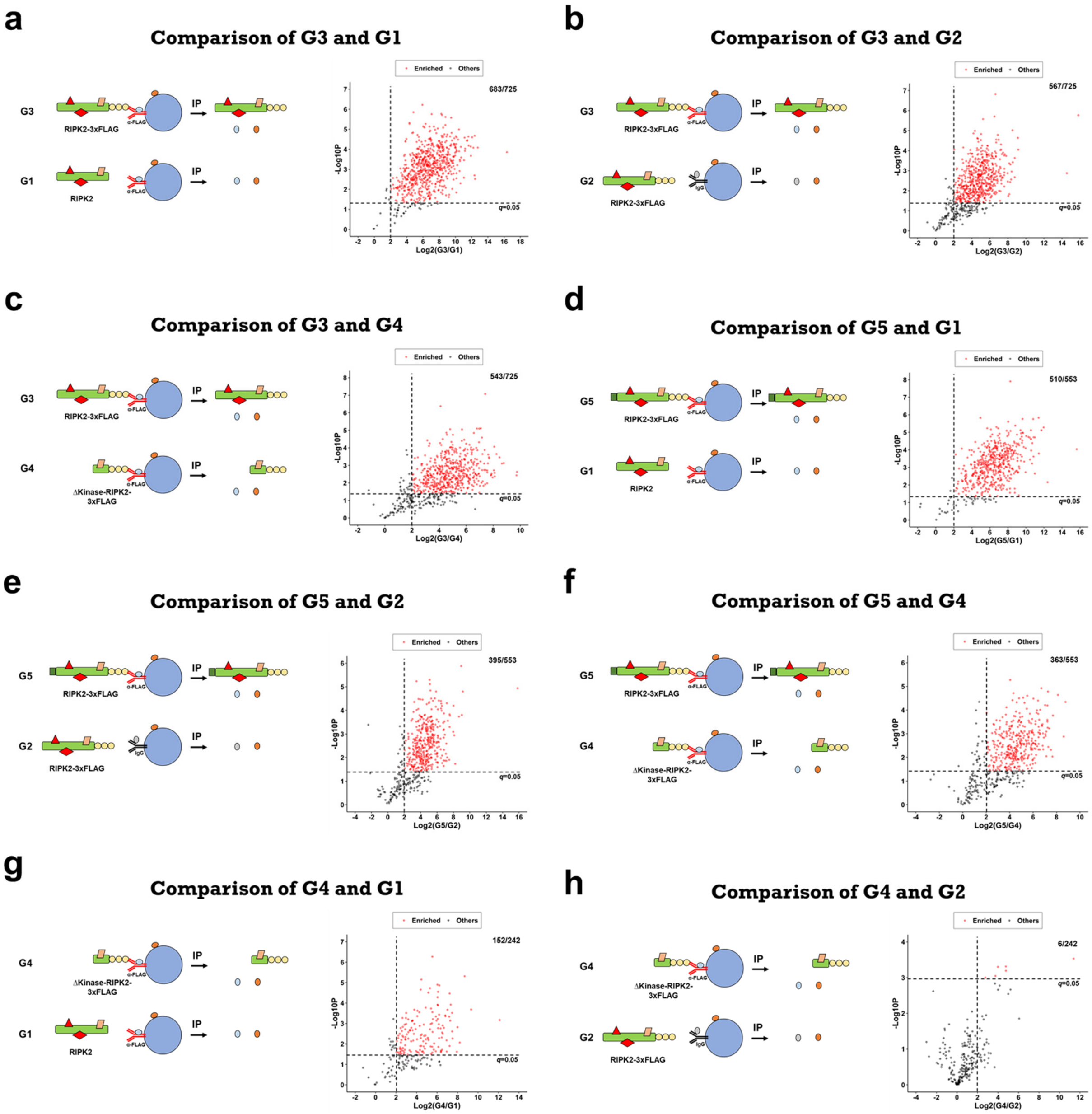


**Extended Data Fig. 29 | Label-free proteomic comparison of different immunoprecipitation groups. a-h**, Schematic for data analysis (left) and volcano plot (right) of significantly enriched protein groups ($q$ < 0.05, $Log_2Ratio > 2$, shown as red dots).

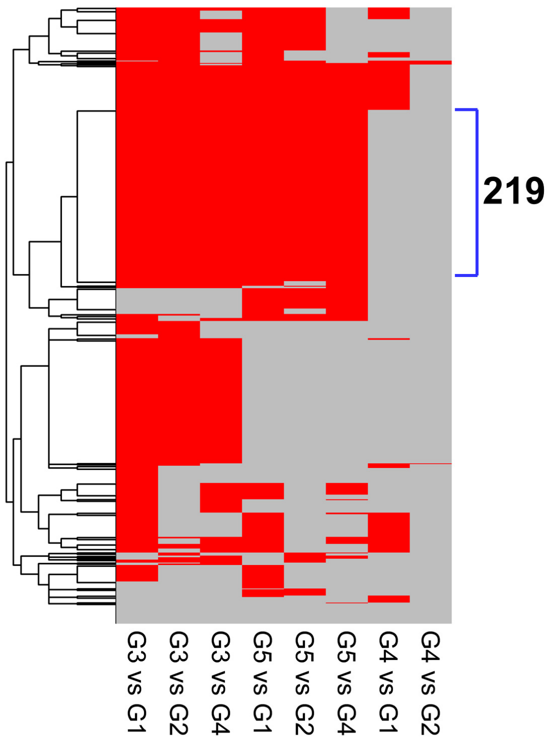


**Extended Data Fig. 30 | Identification of protein candidates associated with the kinase domain, but not other regions, of cytoplasmic RIPK2.** Two criteria were applied to identify the candidates. Firstly, proteins are significantly enriched ($q < 0.05$, $Log_2Ratio > 2$) in both experimental groups (*i.e.*, G3 and G5) compared with the three control groups (*i.e.*, G1, G2, and G4). Secondly, proteins are not significantly enriched in the control group G4 compared with the other control groups G1 and G2. The second criterion was used to further exclude the identification of proteins that may bind to the non-kinase-domain regions of RIPK2, which are not important for RIPK2's regulation of c-Myc.

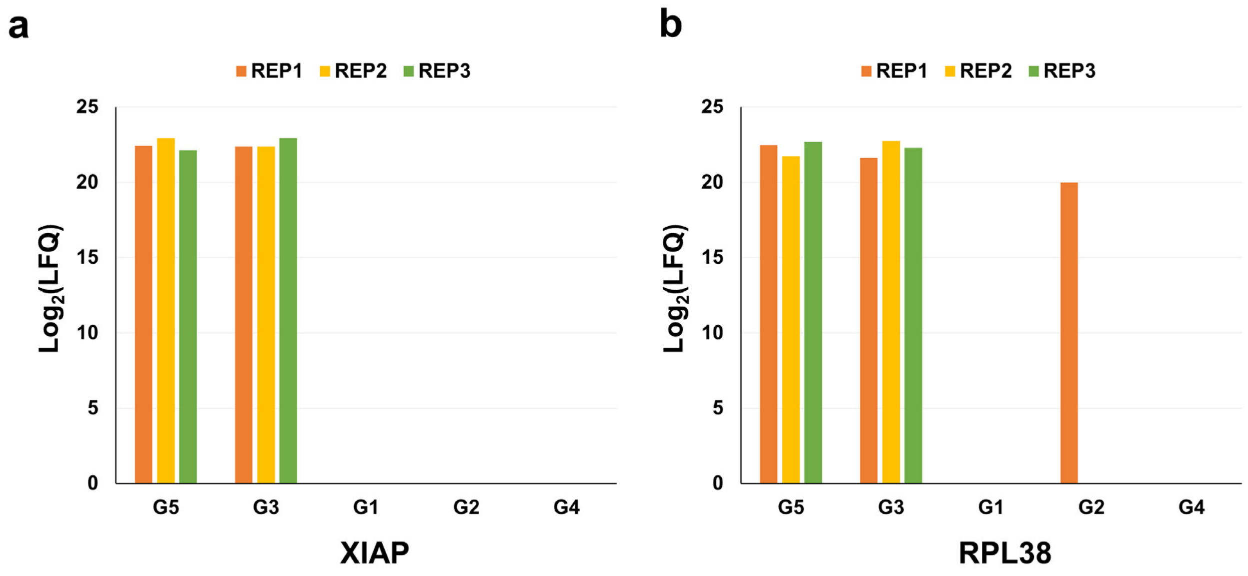


**Extended Data Fig. 31 | Bar plots of the log2-transformed LFQ intensities of XIAP (a) and RPL38 (b), two known RIPK2-interacting proteins.** See the Extended Data Fig. 26 for the five groups G1-G5. REP1-3 stands for replicate #1-#3.

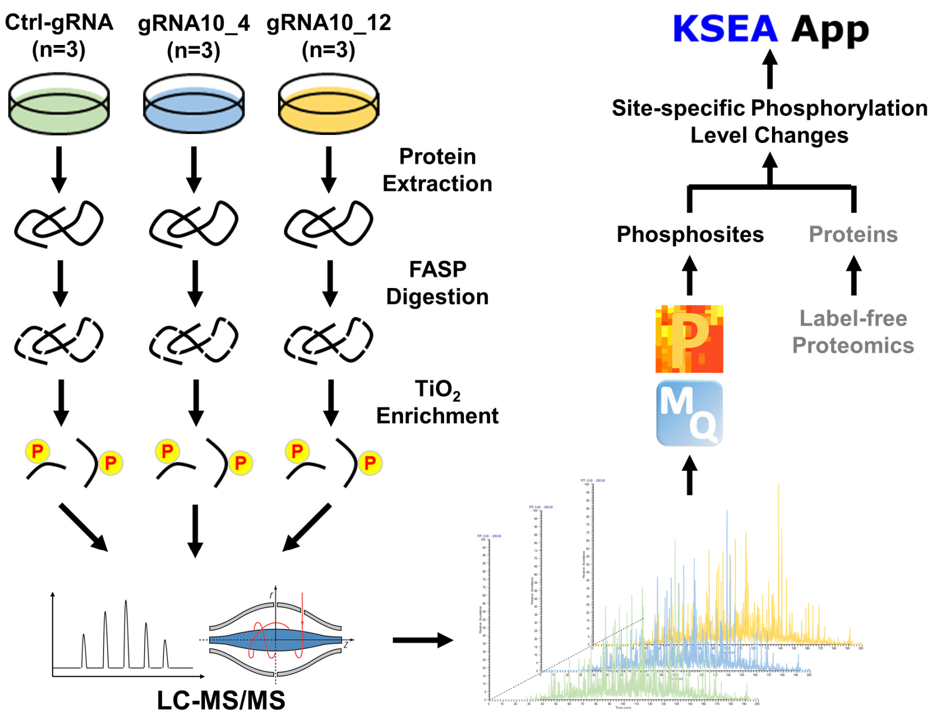


**Extended Data Fig. 32 | Schematic of label-free phosphoproteomics analysis to identify kinases downstream of RIPK2.** Proteins were extracted from control and RIPK2-KO (single-cell clones #4 and #12) PC3 cells, digested into tryptic peptides by filter-aided sample preparation (FASP), and subjected to titanium dioxide ($TiO_2$) enrichment of phosphopeptides. The enriched phosphopeptides were analyzed by liquid chromatography-tandem mass spectrometry (LC-MS/MS), and the resulting RAW files were analyzed with MaxQuant (MQ) to identify phosphopeptides and localize phosphosites. Perseus (P) was used to perform statistical analysis to identify differentially expressed phosphoproteins at the site-specific level. To compute the phosphorylation level changes, the relative LFQ intensities of phosphosites were normalized against the relative abundance of corresponding proteins, which was obtained from previous label-free proteomic analysis of the same cell clones. Finally, all significantly changed ($p < 0.05$) phosphosites were analyzed by the online KSEA App program to quantify the activity changes of inferred kinases.

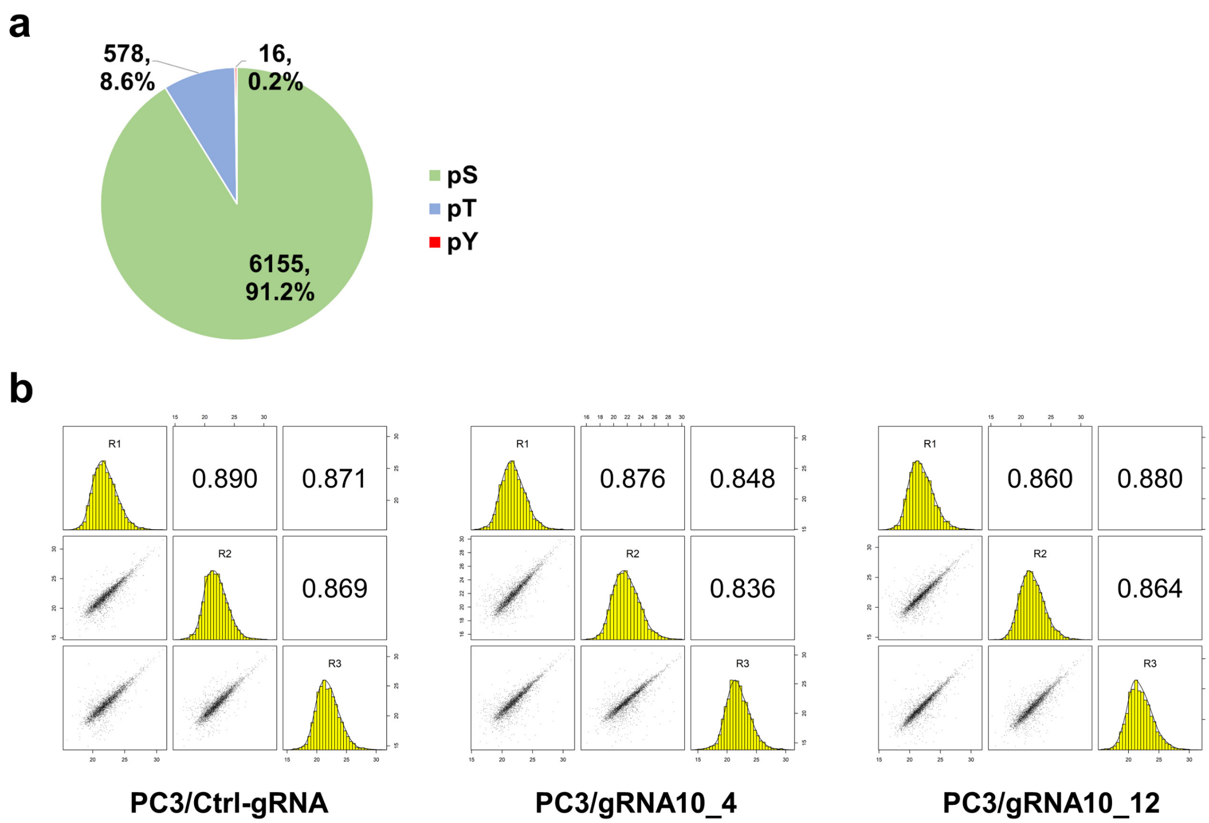


**Extended Data Fig. 33 | Quality assessment of the phosphoproteomics dataset. a,** Pie chart of identified phosphoserine (pS), phosphotheonine (pT), and phosphotyrosine (pY) sites identified from control and RIPK2-KO PC3 cells (identification false discovery rate < 1%, localization probability > 0.75). **b,** Pearson correlation coefficients between replicate runs of the label-free phosphoproteomic analysis. PC3/Ctrl-gRNA: PC3 cells stably transfected with non-targeting control gRNA; PC3/gRNA10_4: PC3 cells stably transfected with RIPK2-targeting gRNA#10, single clone #4; PC3/gRNA10_12: PC3 cells stably transfected with RIPK2-targeting gRNA#10, single clone #12; numbers in boxes: Pearson correlation coefficients; R1-R3: biological replicates #1-#3; scatter plots: distribution of log2-transformed LFQ intensities for phosphosites.

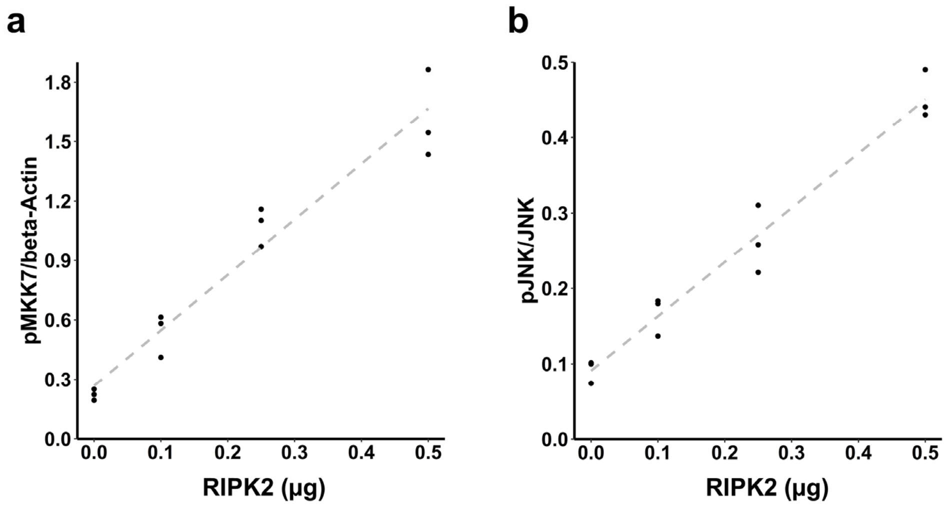


**Extended Data Fig. 34 | Scatter plots of pMKK7-S271 (a) and pJNK-T183/Y185 (b) levels in response to different RIPK2 dosages.** See Fig. 4f for representative Western blotting images.

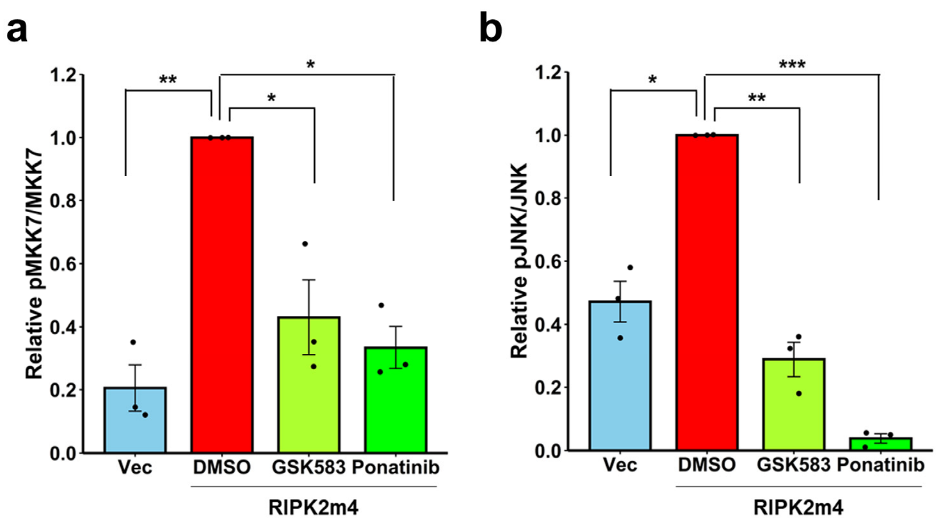


**Extended Data Fig. 35 | Bar plots of the relative MKK7 (a) and JNK (b) phosphorylation levels.** See Fig. 4g for representative Western blotting images. Data represent mean ± standard error of the mean. Unpaired two-tailed Student's *t*-test. * $P < 0.05$, ** $P < 0.01$, *** $P < 0.001$. pMKK7 stands for phospho-MKK7-S271; pJNK stands for phospho-JNK-T183/Y185; Vec, vector control.

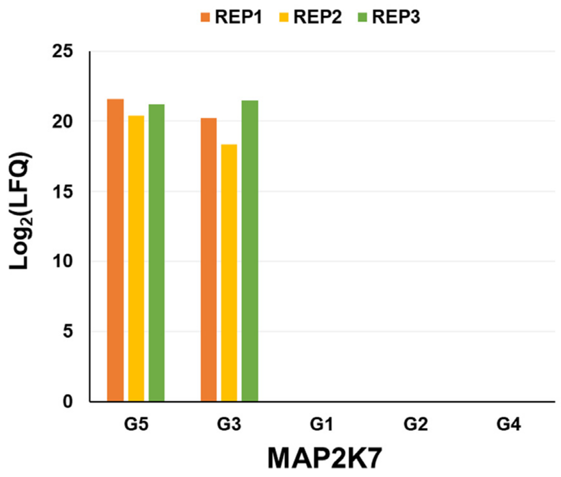


**Extended Data Fig. 36 | Bar plot of log2-transformed LFQ intensities of MAP2K7 (*i.e.*, MKK7) in different immunoprecipitation samples.** See the Extended Data Fig. 26 for the five groups G1-G5, among which G5 and G3 were experimental groups whereas G1, G2, and G4 were control groups. REP1-3 stands for replicate #1-#3.

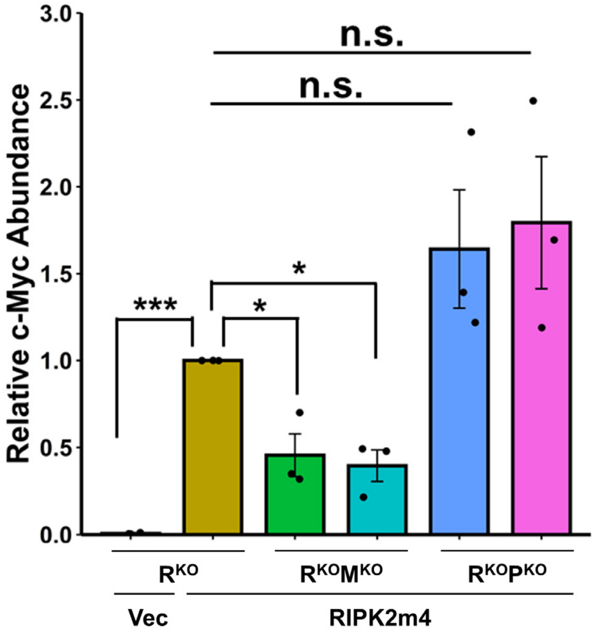


**Extended Data Fig. 37 | Bar graph of relative c-Myc protein abundance under indicated conditions.** See Fig. 4k for a representative Western blotting image. Data represent mean ± standard error of the mean. Unpaired two-tail Student's *t*-test; * $P < 0.05$; *** $P < 0.001$; n.s. not significant.

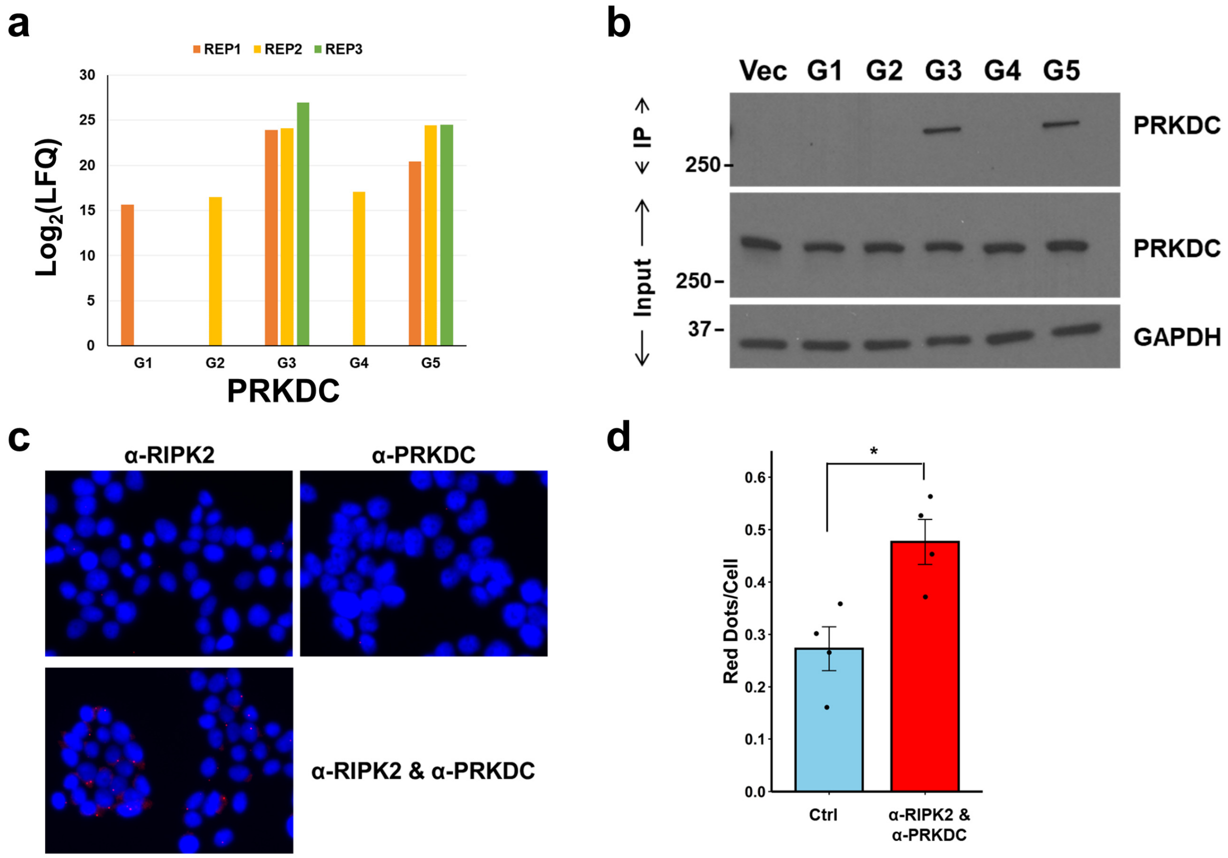


**Extended Data Fig. 38 | PRKDC colocalizes and associates with RIPK2. a,** Bar plot of the log2-transformed LFQ intensities of PRKDC in different immunoprecipitation samples. Here, G3 and G5 were experimental groups, whereas G1, G2, and G4 were control groups. See Extended Data Fig. 26 for more details. **b,** Western blotting analysis to probe PRKDC in protein complexes isolated by immunoprecipitation. **c,** Cellular images of the proximity ligation assay detecting the association of RIPK2 and PRKDC in parental HEK293T cells. **d,** Bar graph of the proximity ligation red dots per cell. The numbers under the control condition are the sum of dots per cell detected using an anti-RIPK2 or an anti-PRKDC antibody alone.

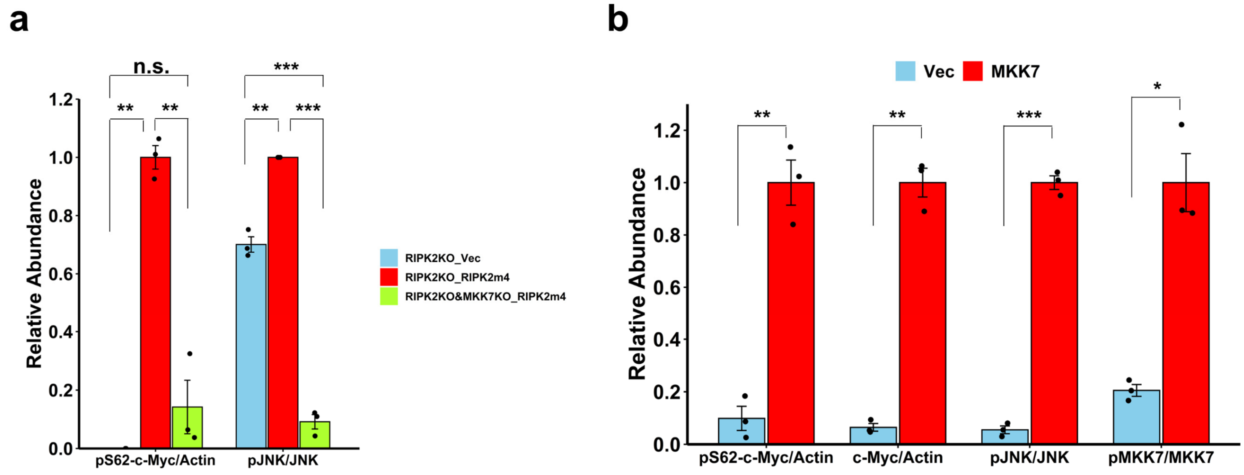


**Extended Data Fig. 39 | Bar graph of relative abundance of different proteins under indicated conditions. a,** Bar graph of relative abundance of pS62-c-Myc and pJNK-T183/Y185 in HEK293T cells under indicated conditions. See Fig. 4l for representative Western blotting images. **b,** Bar graph of relative abundance of pS62-c-Myc, c-Myc, pJNK-pT183/Y185, and pMKK7-S271 in HEK293T cells under indicated conditions. See Fig. 4m for representative Western blotting images. Data represent mean ± standard error of the mean. Unpaired two-tail Student's *t*-test; * $P < 0.05$; ** $P < 0.01$, *** $P < 0.001$; n.s. not significant.

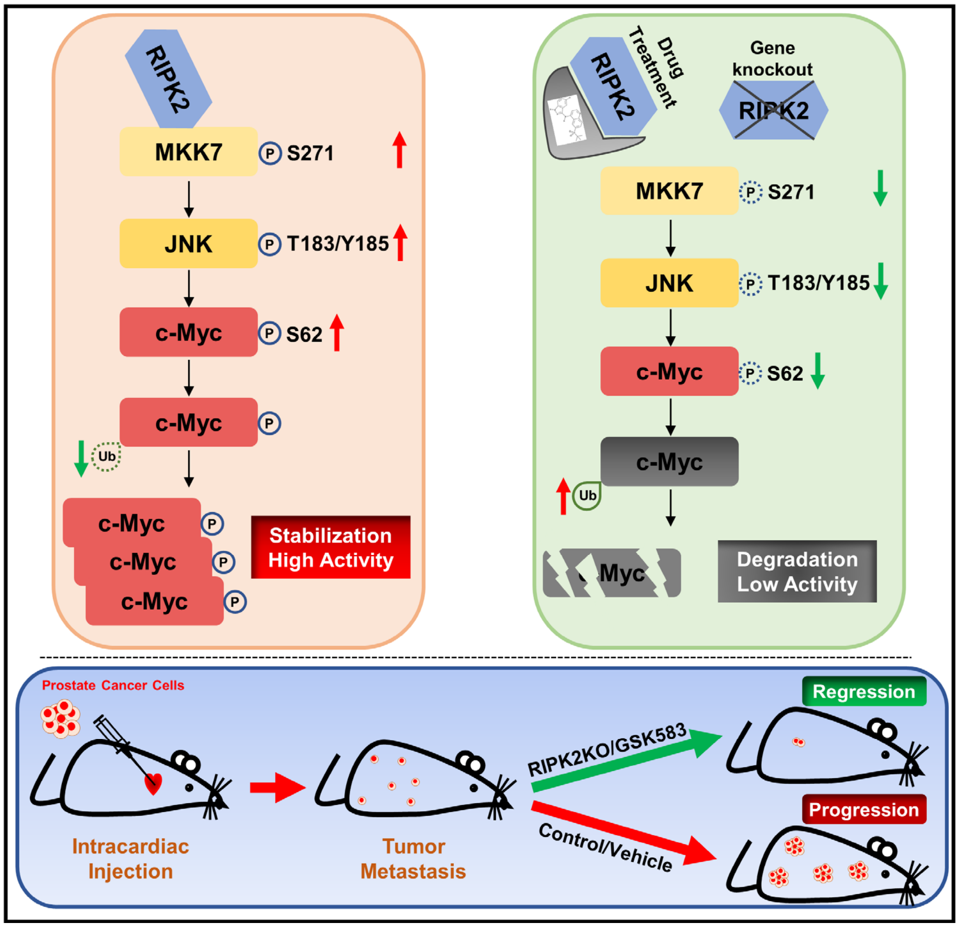


**Extended Data Fig. 40 | Schematic of targeting RIPK2 to destabilize c-Myc and suppress prostate cancer metastatic progression.** In prostate cancer cells, overexpressed RIPK2 associates with MKK7 and activates the MKK7/JNK phosphorylation pathway, resulting in the phosphorylation and stabilization of c-Myc protein and thus an increase in c-Myc activity. Genetic and pharmacological inhibition of RIPK2 suppress the MKK7/JNK/c-Myc phosphorylation pathway, promote the degradation of c-Myc, and attenuate c-Myc activity, consequently suppressing prostate cancer metastatic progression.

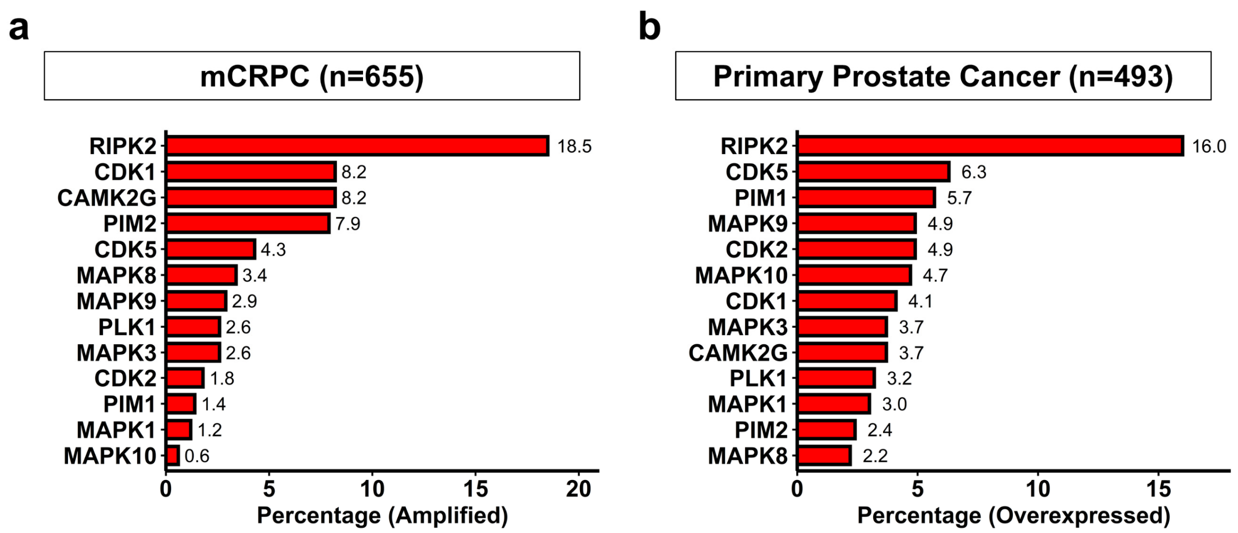


**Extended Data Fig. 41 | RIPK2 is much more frequently amplified and overexpressed than direct c-Myc-Ser62 kinases in human prostate cancer. a,** Bar plot of the frequency of gene amplification in mCRPC tissue specimens. **b,** Bar plot of the frequency of gene overexpression (Z>2, normalized against diploid samples) in TCGA PanCancer Atlas prostate cancer tissue specimens.

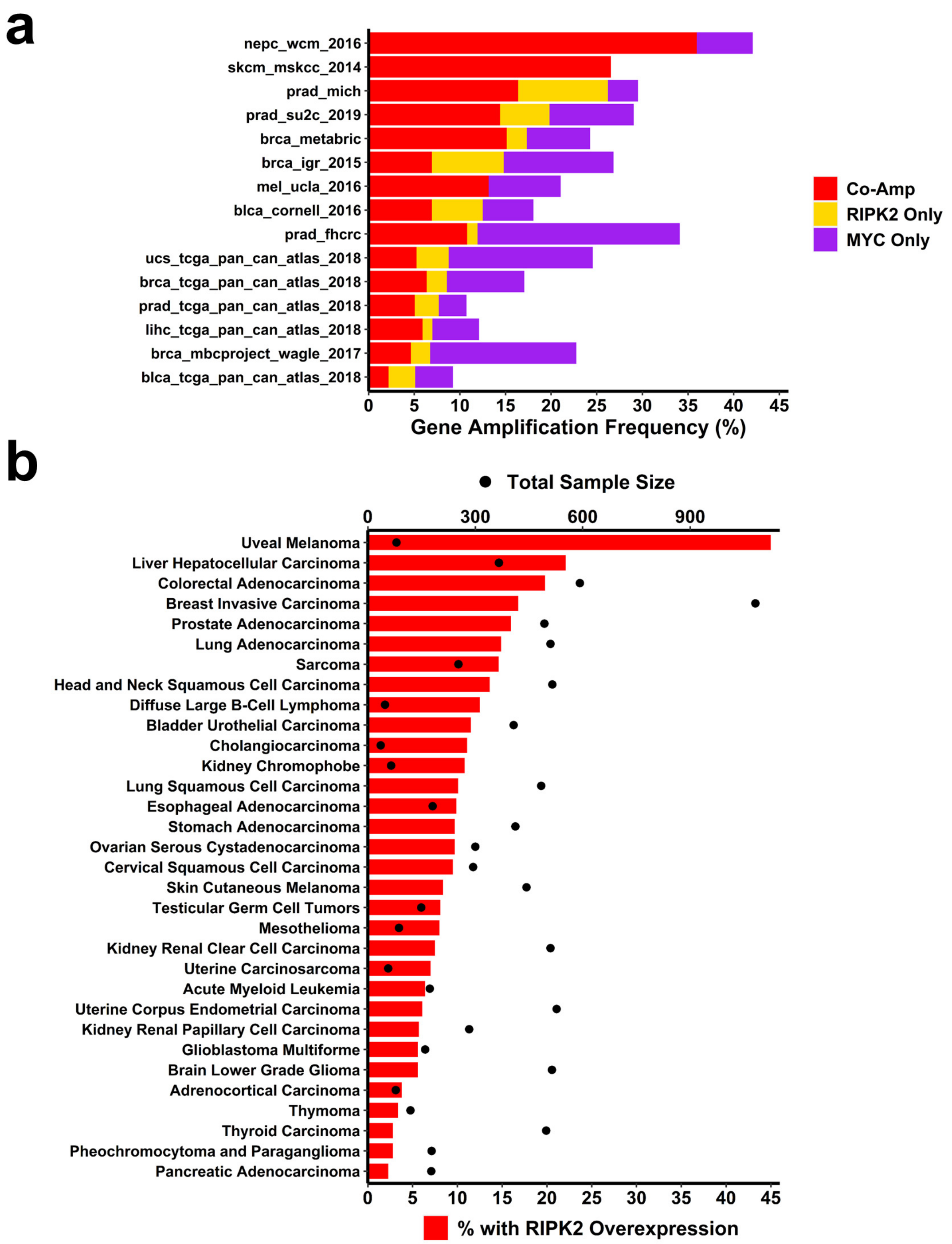


**Extended Data Fig. 42 | RIPK2 is frequently amplified or overexpressed in several other cancer types.** **a,** Bar plot of percentages of tumors with RIPK2 gene amplification in different cancer genomics studies. Red shows the co-amplification of RIPK2 and MYC, orange represents amplification of RIPK2 but not MYC, and purple indicates amplification of MYC but not RIPK2. **b,** Bar plot of percentages of tumors with RIPK2 overexpression (Z > 2, normalized against diploid samples) in the TCGA PanCancer Atlas studies. The "total sample size", shown in black dots, represents the number of tissue specimens with mRNA expression data. Bar lengths correspond to the percentages of tumors with RIPK2 overexpression (Z > 2).

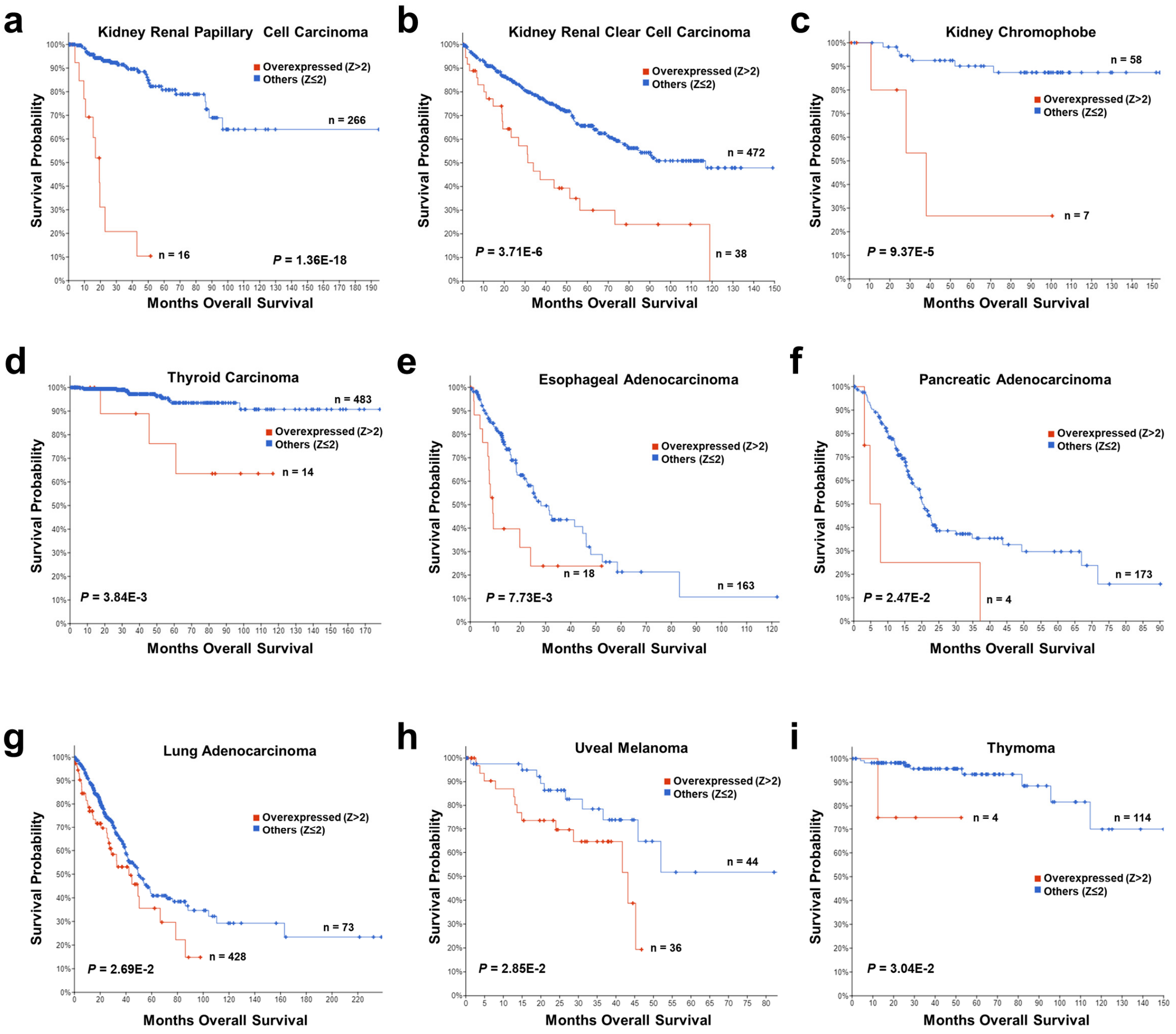


**Extended Data Fig. 43 | RIPK2 overexpression is associated with significantly worse overall survival in some other cancer types. a,** Kidney renal papillary cell carcinoma (n=282). **b,** Kidney renal clear cell carcinoma (n=510). **c,** Kidney chromophobe (n=65). **d,** Thyroid carcinoma (n=497). **e,** esophageal adenocarcinoma (n=181). **f,** Pancreatic adenocarcinoma (n=177). **g,** Lung adenocarcinoma (n=501). **h,** Uveal melanoma (n=80). **i,** Thymoma (n=118). Kaplan-Meier overall survival analysis of cancer patients (TCGA PanCancer Atlas) stratified based on whether RIPK2 is significantly ($Z > 2$, normalized against diploid samples) overexpressed. Two-sided log-rank test.

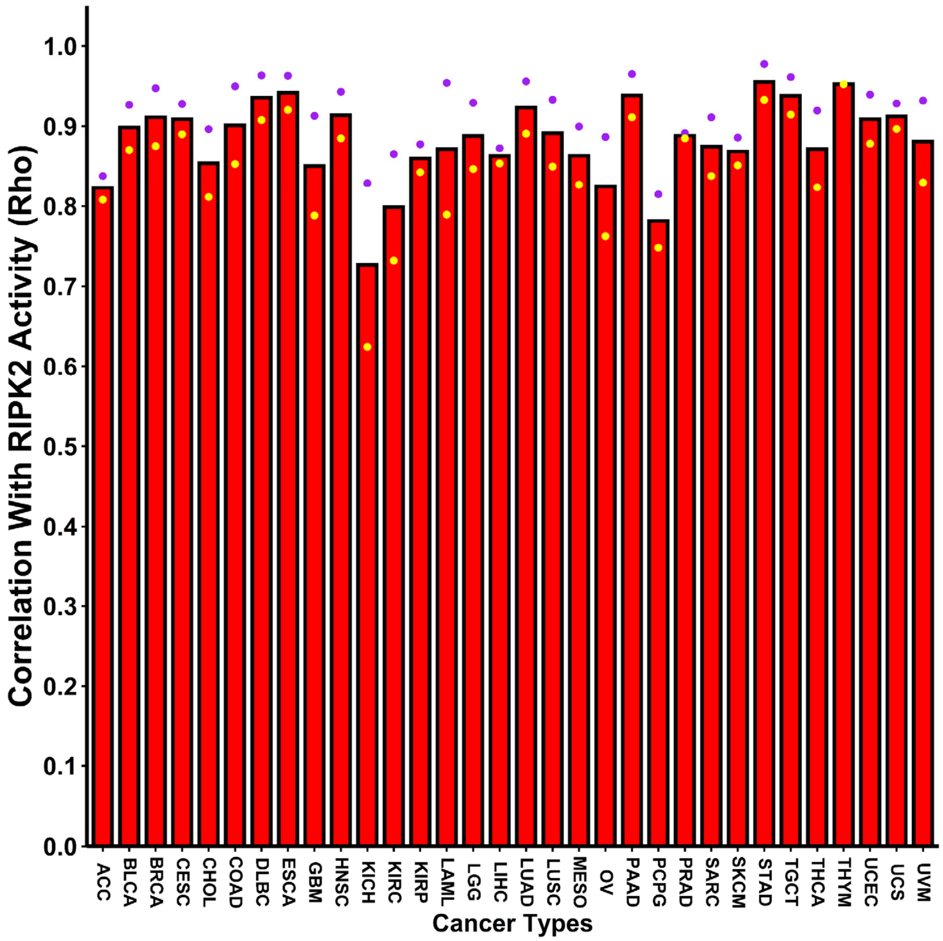


**Extended Data Fig. 44 | RIPK2 activity scores are strongly associated with MYC activity scores across different cancer types.** Bar heights represent the average Spearman's correlation coefficients of RIPK2-induced activity Z scores with the activity Z scores of two MYC target gene sets (Hallmark_MYC_Targets_V1 and Hallmark_MYC_Targets_V2). Purple and yellow dots indicate the correlation coefficients of RIPK2-induced activity Z scores with the MYC_V1 and the MYC_V2 activity Z scores, respectively.
